# The modular evolution of multiheme cytochromes *c* bucks the general trend observed in proteins

**DOI:** 10.1101/2025.05.29.656830

**Authors:** Ricardo Soares, Catarina M. Paquete, Ricardo O. Louro

## Abstract

Multiheme cytochromes *c* are versatile enzymes which contribute to key steps in diverse biogeochemical cycles of elements and are also key players in microbial electrochemical technologies. We previously showed that these enzymes evolve by grafting and pruning of cytochrome *c* modules. Here, we extended our analysis and found that grafting can also involve the incorporation of domains of other protein families and of small peptide sequences that affect the heme coordination environment. We show that the addition of hemes to multiheme cytochromes *c* occurs exclusively by integrating heme-binding peptides and that gain of hemes is equally probable along the protein sequence. By contrast, heme loss occurs by the loss of heme-binding peptides and the accumulation of point mutations. Notably, heme-binding motif loss is disproportionately more prevalent at the position nearer to the N-terminus. This observation contrasts with the general trend observed in proteins, which are usually more conserved at the N-terminus, and likely reflects the way in which multiheme cytochromes *c* are assembled. Overall, this work completes the picture of how multiheme cytochromes *c* evolved to achieve the diversity of structures and functions found in nature, and sets them apart from other proteins with respect to the drivers for their evolution. This has contributed to the diversity of roles that they play in various biogeochemical cycles and has implications for engineering artificial variants to enhance biotechnological applications.

## Introduction

It is widely recognised that extant proteins originated by the fusion of small peptides (Eck and Dayhoff 1966; Söding and Lupas 2003; Romero Romero et al. 2016), and this *modus operandi* later extended to fusions of protein domains, leading to larger and more complex structures (Kummerfeld and Teichmann 2005; Pasek et al. 2006). Further sequence divergence by selection and drift, subsequently led to the great diversity of extant proteins (Jayaraman et al. 2022). Numerous evolutionary studies have focused on heme proteins (Hardison 2012; Sello et al. 2015; Sello et al. 2015; Pillai et al. 2020; Hansen et al. 2021; Harris et al. 2022), including *c*-type cytochromes ( Dickerson, 1971; Li et al., 2024; Margoliash, 1963). However, the evolution of multiheme *c*-type cytochromes (MHC), which are, in general, more complex, has received little attention.

MHC are metalloproteins found in prokaryotes that contain multiple covalently bound heme *c* cofactors. In recent years, MHC have caught the attention of the scientific community owing to their pivotal role in diverse biogeochemical cycles of the elements (Lycus et al. 2023) (Fonseca et al. 2013; Costa et al. 2019; Edwards, White, et al. 2020; Tikhonova et al. 2023) (Igarashi et al. 1997; Einsle et al. 1999; M. Akram et al. 2019), catalytic reactions such as fumarate reduction (Chen et al. 1994; Paquete et al. 2014), and sensing the presence of small molecules (Catarino et al. 2010). Additionally, electroactive bacteria that contain complete extracellular electron transfer pathways composed of MHC have gathered significant attention in the context of the development of bioelectrochemical systems and bioelectronics applications (Ing et al. 2018; Fonseca et al. 2021). The establishment of the covalent bonds between the heme and the polypeptide is catalysed by dedicated cytochrome *c* maturation machinery found in the periplasmic space of prokaryotes. It recognises specific heme-binding motifs in the amino-acid sequence, such as the CxxCH heme binding motif and its variants (Verissimo and Daldal 2014; Paquete et al. 2019; Mendez et al. 2022). Each heme is coordinated by the two cysteines via thioether bonds, and the iron is axially coordinated by the histidine found in the motif. This coordination of the heme *c* imposes a strong constraint for the folding of MHC that is absent in proteins without covalently linked co-factors(Dolla et al. 1995; Florens et al. 1995; Chen et al. 2022). Furthermore, the covalent binding of the hemes *c* shields them from entropically driven dissociation (Arnesano et al. 1999; Allen et al. 2003). As a consequence, all known multiheme cytochromes with more than three hemes contain *c*-type hemes (Cantor and Schimmel 1980; Edwards, Richardson, et al. 2020), and display a large number of heme *c* cofactors in a relatively short polypeptide chain with ratios as low as 20 amino-acid residues per heme (Sharma et al. 2010; Soares et al. 2024). However, this imposes evolutionary constraints in which the traditional structural and functional fitness landscapes do not apply (Gilson et al. 2017).

The number of hemes in MHC varies substantially between the different families (Soares et al. 2024). So far, the MHC with the highest number of hemes per polypeptide chain for which the structure has been experimentally determined is HmcA with 16 hemes (Czjzek et al. 2002; Matias et al. 2002), but genome mining shows MHC coding sequences with more than 100 heme-binding motifs per polypeptide (Leu et al. 2020; Wasmund et al. 2024). Additionally, some MHC families possess a high number of paralogues, demonstrating a high degree of flexibility and diversification (Soares et al. 2024). This makes MHC an interesting target to study the dynamic mechanisms for metalloprotein evolution. Gene fusion and fission events have marked the evolution of MHC and represent step changes that, by domain shuffling, provided an important source for diversification (Klotz et al. 2008; Soares et al. 2022). However, little is known about the mechanisms for more gradual changes that can shape the evolution of MHC, such as the addition or removal of just one heme, as is often found in homologous MHC (Edwards et al. 2012). Here, we took advantage of the current revolution in protein structuromics (Perrakis and Sixma 2021; Thornton et al. 2021) and combined it with a thorough phylogenetic analysis of the evolution of MHC to address this open question and provide foundations to understand the global mechanisms that shape the evolution of MHC. We found that the fusion of small peptide sequences or domains containing other cofactors is an evolutionary mechanism for the diversification of MHC. In addition to these processes, we also found that fission is more prevalent than fusion, in particular at the N-terminal.

## Results and Discussion

### Different MHC families exhibit varying degrees of diversity

To study the mechanisms that shaped the gradual evolution of MHC, we focused on every family that has been experimentally characterised in terms of structure and function (for reviews, see (Bayar et al., 2025; Paquete et al., 2019)). We collected highly homologous sequences for each of these families. While for some, we were only able to collect three highly homologous protein sequences, for others, we could obtain more than 1,000 (Table 1). Some families are homogeneous in the number of heme-binding motifs in the polypeptide sequence within our dataset, namely NrfA, NrfB, MccA, STC, MtrC, cytochrome *c*554, and HAO, while other families (FccA, ActA6, ActA5, OTR, MtrA, NrfH, PufC, cytochrome *c*7, cytochrome *c*3, OmcE, 16HmcA, GSU1996, IhOCC, OmcS, ONR, UndA, Lpc552 AvECn and PcECN) displayed heterogeneity in the number of heme-binding motifs within our dataset (fig. 1). Excluding protein families that were poorly represented (< 10 highly homologous sequences), the average amino acid substitution distance within each protein family was approximately 50%. NrfH showed the highest diversity with a distance of 68%, while OmhA, for which we could only collect four protein sequences from a single genus, *Carboxydothermus*, showed the lowest diversity, containing an average distance of 17% between the sequences of the dataset (Table 1).

**Fig. 1:**
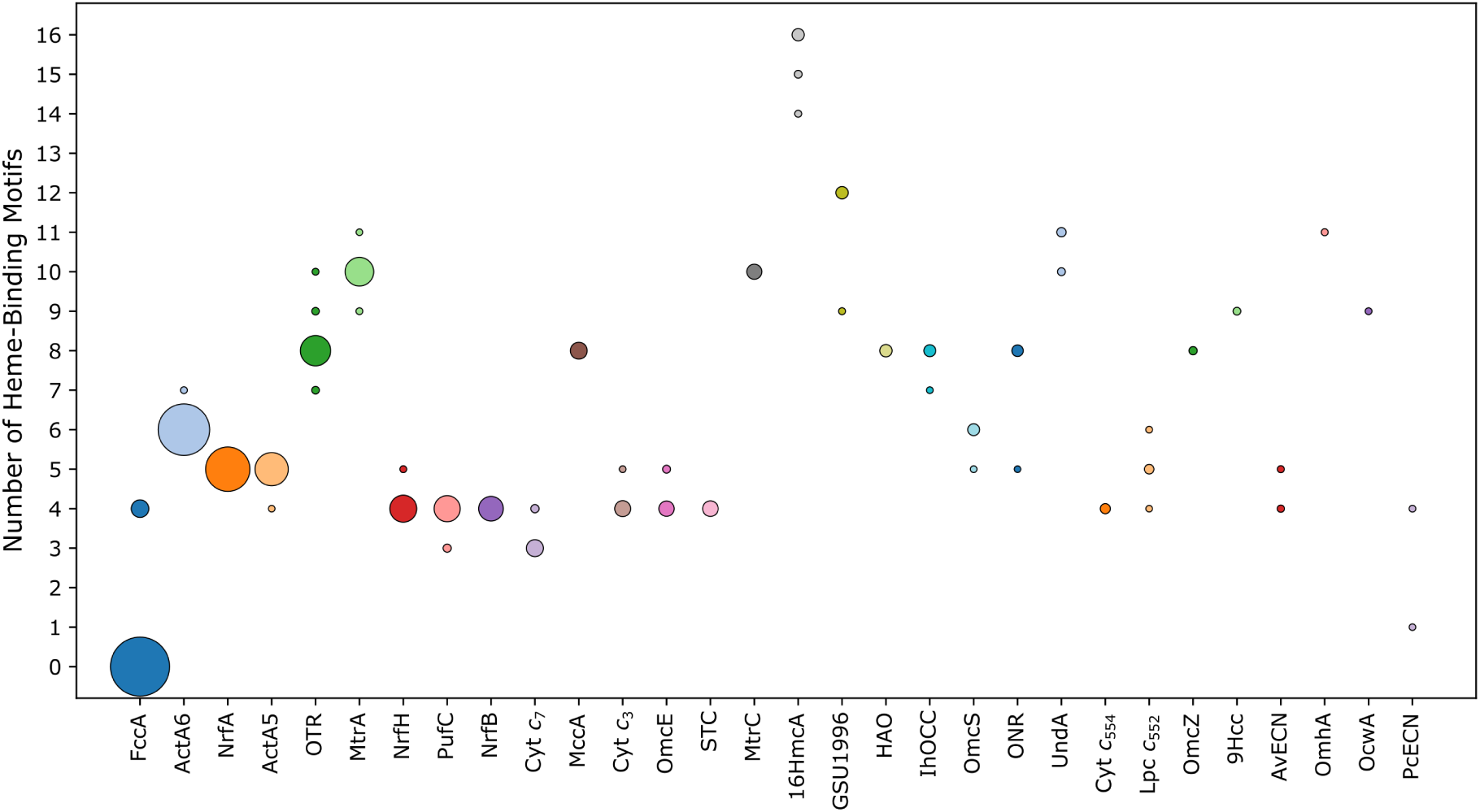
Bubble chart showing the distribution of the number of heme-binding motifs for each group of homologous MHC that was collected from the RefSeq select database. MHC groups are ordered by dataset size (Table 1), and the circle sizes are scaled according to the number of sequences containing a particular number of heme-binding motifs.

**Table 1.** Number of MHC sequences collected from the RefSeq select database for each protein family. The number of heme-binding motifs (HBM) and average amino acid substitution distance (**x*̄***) are shown. The number of sequences that compose each subgroup containing a different number of heme-binding motifs within a given family is shown within parentheses.

| Protein | № sequences | № HBM | $\bar{x}$ distance ± SE |
| --- | --- | --- | --- |
| FccA | 1956 | 0 (1816); 4 (140) | 0.663 ± 0.011 |
| ActA6 | 1379 | 6 (1376); 7 (3) | 0.525 ± 0.014 |
| NrfA | 1011 | 5 | 0.589 ± 0.012 |
| ActA5 | 559 | 5 (558); 4 (1) | 0.560 ± 0.020 |
| OTR | 480 | 7 (9); 8 (460); 9 (8); 10 (3) | 0.551 ± 0.013 |
| MtrA | 414 | 9 (3); 10 (410); 11 (1) | 0.567 ± 0.018 |
| NrfH | 364 | 4 (361); 5 (3) | 0.683 ± 0.022 |
| PufC | 350 | 3 (13); 4 (337) | 0.549 ± 0.016 |
| NrfB | 297 | 4 | 0.601 ± 0.022 |
| Cyt *c*<sub>7</sub> | 144 | 3 (130); 4 (14) | 0.495 ± 0.031 |
| MccA | 128 | 8 | 0.400 ± 0.011 |
| Cyt *c*<sub>3</sub> | 114 | 4 (112); 5 (2) | 0.629 ± 0.027 |
| OmcE | 113 | 4 (102); 5 (11) | 0.649 ± 0.021 |
| STC | 106 | 4 | 0.468 ± 0.026 |
| MtrC | 98 | 10 | 0.520 ± 0.011 |
| 16HmcA | 71 | 14 (4); 15 (10); 16 (57) | 0.593 ± 0.014 |
| GSU1996 | 62 | 9 (4); 12 (58) | 0.449 ± 0.016 |
| HAO | 59 | 8 | 0.459 ± 0.013 |
| IhOCC | 55 | 7 (2); 8 (53) | 0.541 ± 0.013 |
| OmcS | 54 | 5 (2); 6 (52) | 0.533 ± 0.015 |
| ONR | 49 | 5 (1); 8 (48) | 0.444 ± 0.012 |
| UndA | 35 | 10 (11); 11 (24) | 0.578 ± 0.011 |
| Cyt *c*<sub>554</sub> | 32 | 4 | 0.412 ± 0.019 |
| Lpc c552 | 29 | 4 (1); 5 (26); 6 (2) | 0.460 ± 0.016 |
| OmcZ | 14 | 8 | 0.543 ± 0.014 |
| 9Hcc | 10 | 9 | 0.375 ± 0.017 |
| AvECN | 10 | 4 (6); 5 (4) | 0.572 ± 0.021 |
| OmhA | 4 | 11 | 0.173 ± 0.011 |
| OcwA | 3 | 9 | 0.432 ± 0.015 |
| PcECN | 3 | 1 (1); 4 (2) | 0.191 ± 0.016 |

Given that changes in the number of hemes in MHC can impact the heme-core arrangement and introduce new functions, we further explored the MHC diversity and evolution by phylogenetic reconstruction. We used the sequences of each protein family with at least 10 homologous sequences that showed heterogeneity in the number of heme-binding motifs (fig. S1-S16). We restricted our analysis to events that were associated with high statistical support (ultra-fast bootstrap ≥ 95 % and SH-aLRT ≥ 80 % (Minh et al. 2022)).

### MHC domains can fuse with domains of other protein superfamilies

Our analysis reveals that MHC can fuse with other families of redox proteins. This is exemplified by the flavocytochrome *c*3 (FccA), which in *Shewanella* spp. is a moonlighting protein that functions as a fumarate reductase and also participates in extracellular electron transfer (Fonseca et al. 2013; Paquete et al. 2014). In FccA, the tetraheme MHC module was fused to the N-terminus of a flavoprotein, placing heme 4 at a short distance from the FAD cofactor, which is compatible with the fast electron transfer required for the catalytic reaction (Rothery et al. 2003). The FccA family shows an evolutionary origin from ancestral flavoproteins without heme-binding motifs (fig. 2; fig. S3). The closest homologs were predicted to be periplasmic, but in a sister branch, we also found sequences predicted to be cytoplasmic (supplementary files). Interestingly, in the case of *S. gelidii* and *Turicimonas muris,* this process was reversed by more recent fission events of the MHC module. BLASTp searches against the PDB (supplementary files) for the four most related flavoproteins that serve as an outgroup before the fusion event (WP_116270682.1, WP_049686820.1, WP_013256151.1 and WP_139259373.1) showed homology towards flavoproteins that reduce fumarate and urocanate (Kim et al. 2018; Venskutonytė et al. 2021).

**Fig. 2.**
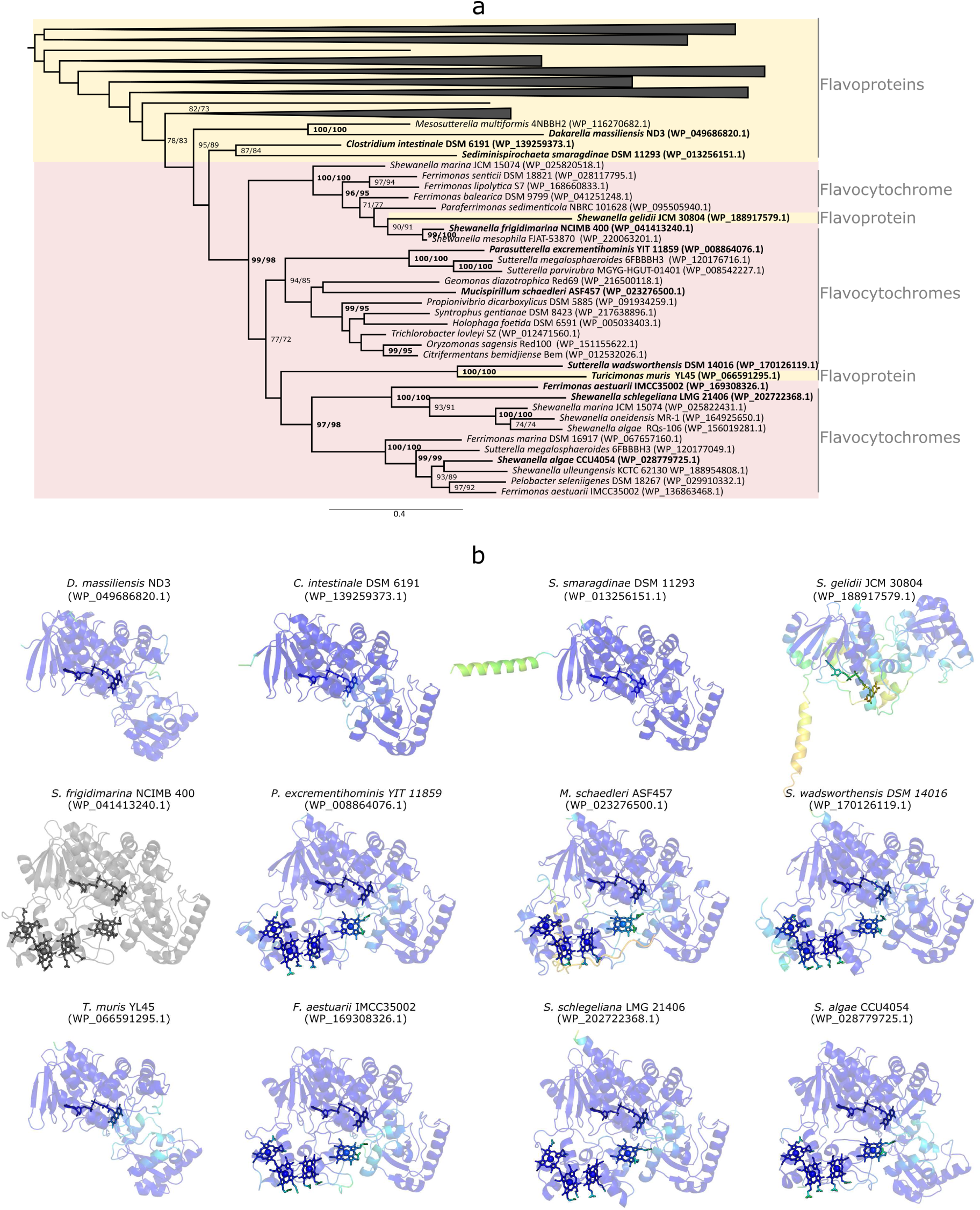
FccA evolution from ancestral flavoproteins that gained an N-terminal MHC module containing four hemes. **a)** Maximum likelihood tree constructed using FccA sequences. Ancestral nodes were collapsed. The SH-likelihood and Ultrafast Bootstrap support are shown near each node. Only confidence values above 70% are shown, and values equal to or above 80/95% are represented in bold. Each sequence is labelled with the corresponding protein family. **b)** Structural models from the flavocytochrome representative sequences represented in bold in the phylogenetic tree of panel a) and closely related flavoproteins predicted using AlphaFold3. The *S. frigidimarina* NCIMB 400 reference structure is represented in grey, while predicted structures are colored from red to blue according to pLDTT confidence values.

### The nature of the distal ligands can be modulated by fusion of non-cytochrome peptides

We also thoroughly analysed the conservation of the distal ligands of the hemes within each MHC family. In our dataset, we found homologous sequences of the decaheme MtrA that lack the canonical distal histidine that coordinates heme 1. The evolutionary reconstruction of the MtrA family shows a branch where both the heme-binding motif and the distal ligand of heme 1 changed from ancestral nodes (fig. 3a). The heme-binding motif changed from Cx2CH to Cx3CH, and the distal ligand from a Histidine located 6 amino-acids after the second heme-binding motif to a Cysteine located 24 amino-acids after that heme-binding motif (figfig. 3b). We used AlphaFold3 to predict the structures of the MtrA homologues with the unusual heme 1 (Cx3CH + distal Cys) and five closely related sequences of a sister branch that conserves the common heme-binding motif and the histidine as the distal ligand (Cx2CH + distal His) of heme 1. The predicted structures showed a pLDDT > 70 for 70-83 % of the sequence, which gave us confidence to align the heme core of these structures against the reference structure of MtrA (PDB:6R2Q (MtrA from *Shewanella baltica* OS185)). We observe that the Cx3CH + distal Cys predicted structures contain an extension of the α-helix that contains the original distal ligand and, in addition, contain also an extra loop containing the new distal Cysteine ligand, when compared with the MtrA reference structure. This new distal ligand coordinates the iron from a different orientation due to the extension of the α-helix (fig.3c). Heme 1 in the predicted structures of MtrA containing Cx3CH + distal Cys deviates more from the reference structure than heme 1 in the predicted structures of MtrA variants containing the Cx2CH + distal His, 1.90 ± 0.24 Å and 0.85 ± 0.32 Å, respectively for the average deviation distance of the iron and nitrogen atoms (fig. 3c and 3d). In addition, heme 1 of the predicted structures of MtrA containing Cx3CH + distal Cys is the most deviated of all hemes (fig. 3d). We repeated the same protocol using RoseTTAFold2, which showed similar results, confirming congruence among different structure prediction methods (fig. S17).

**Fig. 3.**
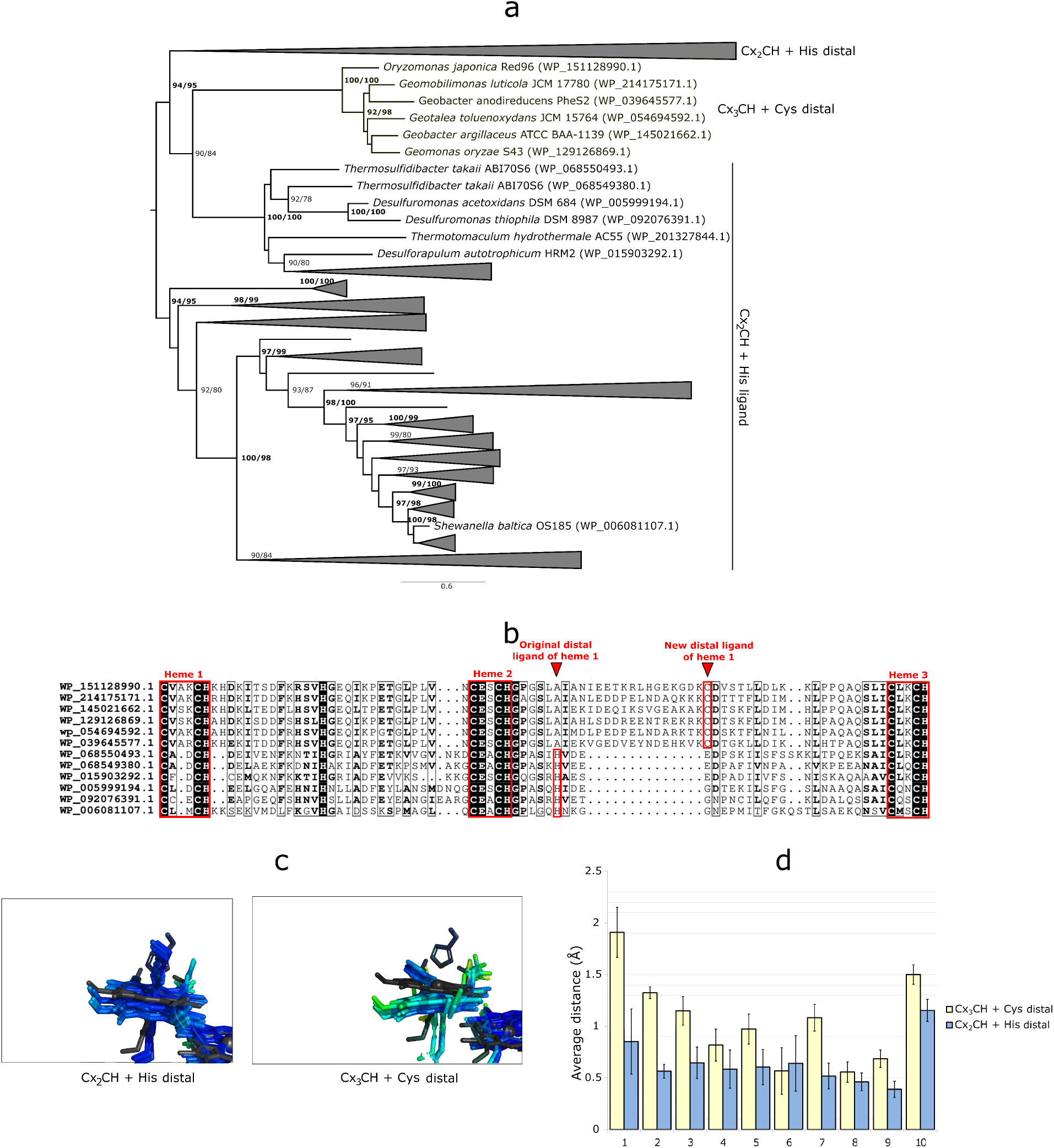
Evolution of MtrA reveals the emergence of an alternative heme-binding motif (CX3CH) and a new distal ligand for heme 1. **a)** Maximum likelihood tree of MtrA sequences. Ancestral nodes were collapsed. Near each node, the SH-likelihood and Ultrafast Bootstrap support are shown. Only confidence values above 70% are shown, and values equal to or above 80/95 % are reported in bold. Each sequence is labelled according to the nature of heme 1 coordination. **b**) N-terminal portion of the alignment of MtrA homologues that contain the alternative heme 1 and closely related sequences containing the canonical heme 1. **c)** Heme core alignment showing heme 1 arrangement of the canonical (left panel) *vs* alternative heme 1 (right panel). The *S. baltica* OS185 reference structure is represented in dark grey, while predicted structures are colored from red to blue according to the predicted local distance difference test (pLDDT) confidence values. **d)** Average distance between the heme rings (iron and nitrogen atoms) of the predicted structures of the Cx3CH + distal Cys and Cx2CH + distal His and the reference MtrA structure (PDB: 6R2Q_A). Error bars represent confidence intervals at 95%.

While distal Cys coordination is less frequent in *c*-type cytochromes than histidine or methionine (Li et al. 2011), there are some well-known examples, such as SoxAX (Kilmartin et al. 2011), PufC (Alric et al. 2004), DsrJ (Grein et al. 2010), TsdA (Denkmann et al. 2012), and NaxS (Mohd Akram et al. 2019). In these cytochromes, the distal Cysteine is important for the function, with most of them playing a role in the catalytic transformation of nitrogen or sulfur compounds (Kilmartin et al. 2011; Denkmann et al. 2012; Mohd Akram et al. 2019). Cysteine is a stronger field ligand than Histidine or Methionine, and therefore hemes with His/Cys coordination have typically lower redox potential than those with His/His and His/Met coordination (Adachi et al. 1991; Raphael and Gray 2002; Alric et al. 2004). In this sense, the MtrA homologues with Heme 1 axially coordinated by His/Cys may have a different function from the MtrA from *Shewanella* spp.. This hypothesis is reinforced by the syntenic gene organisation of operons with MtrA homologues containing the Cx3CH + Cys ligand sequences. These operons do not contain the classical MtrCAB genes found in *Shewanella* spp. Instead, these operons contain an MtrB homologue, an ABC transporter, and response regulator genes (fig. S18), suggesting that they participate in the active transport of some redox-active metabolite.

### Fusion of non-cytochrome peptide sequences contributes to remodelling the heme core arrangement

To further analyse the effect of the changes in the heme binding motif and distal coordination, we made structural predictions on MtrA based on three types of *in silico* mutants for the Cx3CH + distal Cys sequences. We changed *in silico* the heme-binding motif Cx3CH to the canonical Cx2CH (mutCx2CH + distal Cys); we removed the extra 14 amino acids and changed the Cysteine ligand to a Histidine ligand (mutCx3CH + distal His); and only removed the extra 14 amino acids (mutCx3CH + distal Cys) (fig. 4a). Only mutCx3CH + distal His and mutCx3CH + distal Cys present noticeable differences from the original Cx3CH + distal Cys predicted structures, with heme 1 showing less deviation from heme 1 of the experimental reference structure of MtrA (fig. 4b). In the case of mutCx3CH + distal Cys, the cysteine is not predicted to coordinate heme 1 (fig. 4b).

**Fig. 4:**
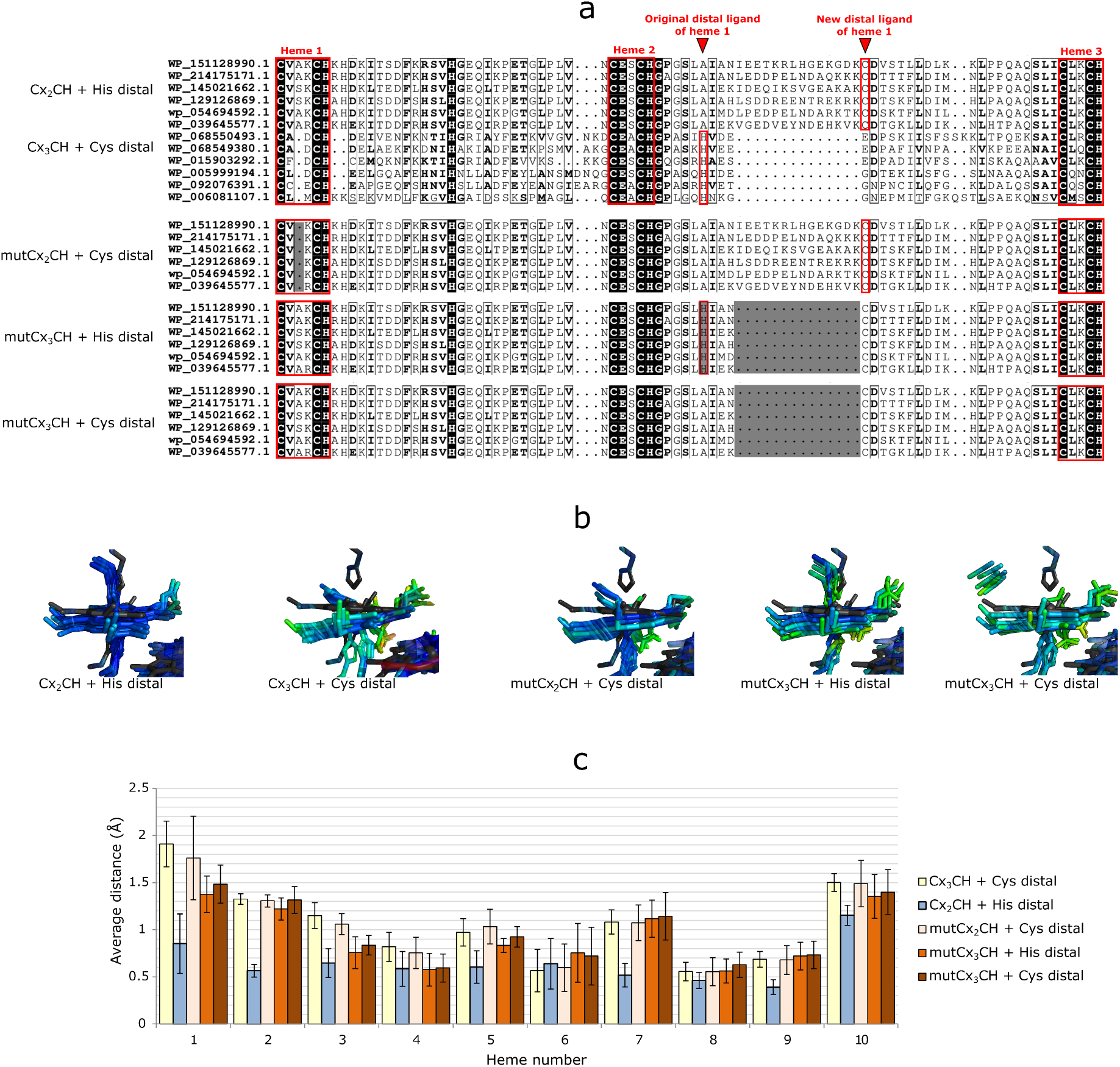
Comparison of the heme core alignment between *in silico* mutants and the reference MtrA structure (PDB: 6R2Q_A) using AlphaFold 3. **a)** alignment of the N-terminal portion of the wild-type sequences and *in silico* mutant sequences. Heme binding motifs and distal ligands are highlighted with red boxes. Mutations are highlighted with a grey shading. **b)** Architecture of the heme 1 environment for the MtrA wild-type sequences and *in silico* mutants when all hemes are aligned to the MtrA experimental structure (in black). Predicted structures are colored from red to blue according to the pLDTT confidence values. **c)** Average distance between the heme rings (iron and nitrogen atoms) of the predicted structures of the wild-type and *in silico* mutants and the reference MtrA structure (PDB: 6R2Q_A). Error bars represent confidence intervals at 95%.

The change of heme arrangement and structure near heme 1 of MtrA homologues that was found here may have significant implications, as the heme arrangement controls the electron transfer flow and the heme reduction sequence within each MHC (Fonseca et al. 2009; Paquete et al. 2010; Fonseca et al. 2012a; van Wonderen et al. 2021; Guberman-Pfeffer and Herron 2025). The two recurring geometric arrangements of pairs of hemes found in MHC are generally designated as stacked and T-shaped (Iverson et al. 1998). Nonetheless, these arrangements encompass a high diversity of possible angles and distances within heme pairs that reveal the modulation of this aspect of MHC by evolutionary pressure or some degree of evolutionary freedom (Wang et al. 2022). Concerning evolutionary freedom, it has been shown that electrostatic interactions among hemes within MHC, in the absence of redox-linked conformational changes, are essentially dominated by the iron-to-iron distance following a Coulombic decay enhanced by a Debye–Hückel shielding factor (Fonseca et al. 2012b). Therefore, changes in the heme arrangements by fusion of small peptide sequences may play an important role in influencing the redox potential of the hemes and shaping both intra- and inter-molecular electron transfer rates, as well as interactions with physiological partners.

### Loss of heme-binding motifs is more frequent than gain

As shown in fig. 1, some MHC conserved the number of heme-binding motifs, while others were prone to diversification. The families of cytochrome *c*3, *c*7, and UndA (fig. S1, S2, and S14) show heterogeneity in the number of heme-binding motifs within the homologous sequences that we collected. However, for those cases, we could not obtain well-resolved phylogenetic trees in the branches that correspond to the change in the number of heme-binding motifs that would allow us to define the chronological order of events with statistical confidence. In the case of OTR, 4 out of 5 events of heme-binding motif loss/gain were also unresolved. For the six families of MHC showing gain of a heme-binding motif (i.e., FccA, NrfH, OmcE, Lpc *c*552, ActA6, and MtrA), the incorporation of the extra heme-binding motif(s) appears to be a step change and not a cumulative transition. This conclusion is based on the observation that the extra heme-binding motifs define alignment gap regions of the closely related homologues without the extra heme-binding motif(s) (fig. S3, S4, S5, S8, S9, S13). This strongly suggests that these events happened by fusion of peptide sequences containing *c*-type heme binding sequences. We were not able to detect any known insertion sequence element by ISEScan and TransposonPSI that would allow us to relate this to any characterised transposable element. By contrast, events of loss of heme-binding motifs (i.e., FccA, PufC, Lp*c*552, ActA(5), IhOCC, ONR, OTR, MtrA, GSU1996, HmcA,) appear to occur by two mechanisms. In some cases, these events are associated with gap regions (fig. S3, S10, S11, S12, S13, and S15), which suggests that they occurred by the loss of heme *c* binding peptide segments. However, in other cases, they are not associated with gap regions, which suggests that they occurred by the accumulation of point mutations (fig. S6, S7, S8, S11, and S13). Overall, we were able to reconstruct the chronology of 23 events that fit our stringent criteria, where 17 were a loss of a heme-binding motif, and 6 were a gain of a heme-binding motif. This suggests that overall, MHC evolution is a two-step process of gaining complexity by fusion of modules and then refining by fission or mutations. N-terminal loss of heme-binding motifs (11 events) was more frequent than loss within the middle of the sequences (6 events) or at the C-terminal (2 events) (fig. 5). By contrast, the gain of heme-binding motifs did not show a bias concerning the position in the polypeptide sequence and we could identify two events of heme-binding motif gain for each relative position within the polypeptide sequence that fit within our critera of high statistical support (fig. 5).

**Fig. 5.**
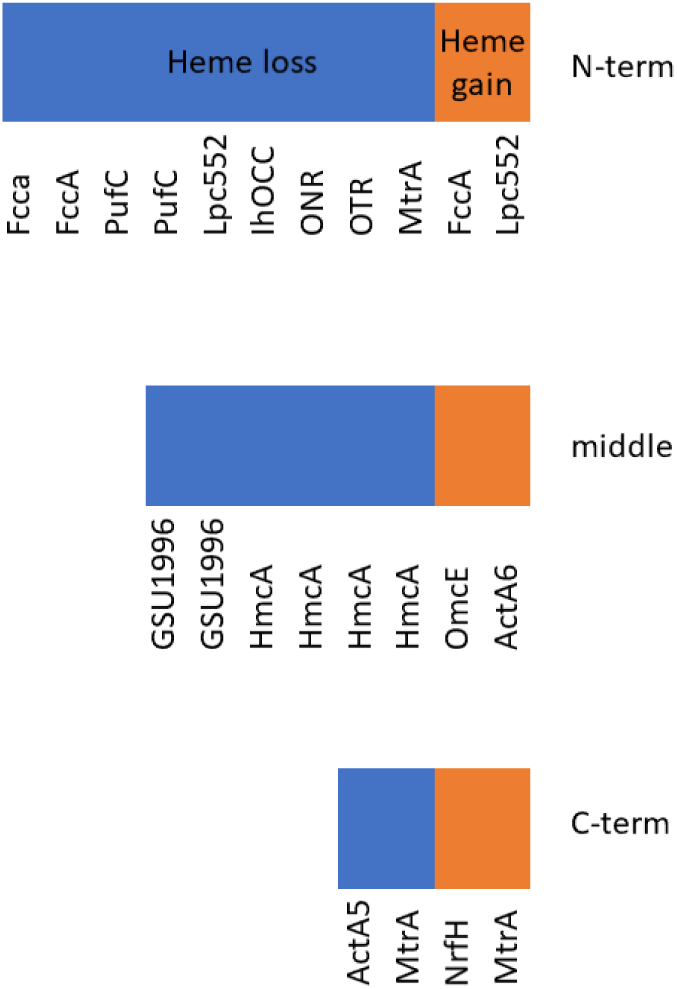
Loss and gain of heme binding motifs parsed according to the location in the polypeptide sequence. N- and C-terminal hemes correspond to the first and last heme, respectively. Middle hemes correspond to hemes between the first and last hemes.

## Conclusions

In a previous work, we found that the group of homologous MHC involved in the biogeochemical cycles of iron (OcwA, OmhA), sulfur (MccA), and nitrogen (NrfA, ONR, HAO, HDH, and IhOCC) (Soares et al. 2022) evolved by the process of fusion and fission of redox modules containing multiple hemes (Soares et al. 2022). This provides a mechanism for the rapid evolution of cytochromes with a number of hemes adequate to the needs of the microbial metabolism, where they are to be integrated. In this work, we map additional mechanisms across a broad range of unrelated MHC. The information gathered in this work paints a more diverse picture of the global mechanisms that drive MHC evolution. It reveals that in addition to fusion and fission of cytochrome modules (Soares et al. 2022), events that occur in well-resolved branches containing multiple MHC homologues fall in four categories: fusion with other protein domains; insertion of peptides that impact the heme core; loss or gain of peptides containing a heme binding motif; and loss of heme-binding motifs by accumulation of point mutations. In particular, it shows differences between the gain and loss of heme binding motifs. Indeed, in all events of gain of heme-binding motifs for which we could find robust phylogenetic statistics, the extra motif is located in a gap region of the closest related sequences that do not contain this extra portion. This indicates that all of these events result from the fusion of peptides containing a *c*-type heme-binding site. By contrast, we found that the events of loss of heme-binding motifs appear to occur by two mechanisms: by gene fission and by the accumulation of point mutations. The mechanism for cytochrome *c* maturation is likely at the basis of this evolutionary asymmetry. Indeed, to bind a heme *c* it is necessary to have a motif containing several aminoacids in a specific sequence, recognised by the heme lyase(Schulz et al. 1998; Verissimo and Daldal 2014). By contrast, a single mutation in any of the cysteines or the histidine of the binding motif is enough to dramatically diminish heme lyase activity (Kranz et al. 2009; Babbitt et al. 2015).

The observation of a bias for more frequent loss of heme binding motifs in the N-terminus was a surprise given that a large survey of single domain proteins showed that the general trend for loss of peptide segments revealed that this happens predominantly in the C-terminus (Weiner 3rd et al. 2006). In prokaryotes, the phenomenon of co-translational folding, where folding of the nascent polypeptide is initiated as soon as it emerges from the ribosome, has been implicated in the establishment of structural asymmetries along the polypeptide sequence (McBride and Tlusty 2021). However, in the case of MHC, co-translational folding needs to be explicitly avoided, because the apo-cytochrome must cross through the SEC system unfolded, to reach the periplasmic space and be assembled as a holo-cytochrome (Page et al. 1998; Stevens et al. 2011). We propose that this fundamental difference in the way MHC are matured is the root cause of the bias towards loss of heme binding domains closer to the N-terminus. It also sets MHC apart from other proteins in the way they evolve to acquire new structure and function, with important implications for their roles in various biogeochemical cycles and for engineering artificial variants to enhance biotechnological applications.

## Materials and methods

### Sequence collection

The sequences of multiheme cytochromes with a structure deposited in the PDB (accession date 03/22/2024) and containing 3 or more hemes were used as query sequences in BLASTp (Altschul et al., 1990) searches. Each sequence was individually used to find homologous sequences containing at least 30% identity, 75 % coverage and an *e*-value of 1^-2^, in both query-subject directions. All sequences that were labelled “partial sequences” in the RefSeq database were removed. Each sequence dataset was aligned using MAFFT 7 (Katoh and Standley 2013) with the L-INS and “Leave gappy regions” options. Overall mean distances were calculated for each dataset using the p-distance model (Nei and Kumar 2000), uniform rates, and pairwise deletion methods within MEGA 11 (Tamura et al. 2021). Standard errors were calculated using the Bootstrap method with 1,000 replications. Heme-binding motifs were identified and counted using the design scripts. Operons were predicted using the Operon Mapper web-server (Taboada et al. 2018). Annotation was provided by the NCBI RefSeq database (Haft et al. 2018) and InterProScan web server searches (Jones et al. 2014).

### Phylogenetic analysis

Datasets containing more than 200 sequences were reduced using MMseqs2 (Steinegger and Söding 2017) with a clustering threshold of 75% identity and 75% coverage. The dataset of FccA, which contained the highest number of sequences mainly composed of flavoproteins with no heme-binding motifs, was further reduced for the flavoprotein using MMseqs but with 50% identity and 75% coverage as clustering thresholds. Aligned sequences were inspected and manually edited, when necessary, in MEGA 11. Low-quality alignment positions were trimmed using TrimAI 1.3 with the “Gappy out” option within the phylemon 2 platform (Sánchez et al. 2011). Phylogenetic trees were generated using IQ-TREE 2 (Minh et al. 2020) on the CIPRESS platform (Miller et al. 2010). ModelFinder (Kalyaanamoorthy et al. 2017) was used for the selection of the best evolutionary model. The best model was selected according to the Bayesian information criterion (BIC). For generation of branch support values, ultra-fast bootstrap (Hoang et al. 2018) and SH-aLRT (Guindon et al. 2010) statistical methods were used. Confidence values were based on 1,000 replications for each method. Heme-binding motif(s) gain/loss events were considered when the respective node containing the change on heme-binding motifs was supported by confidence values of ultra-fast bootstrap ≥ 95 % and SH-aLRT ≥ 80 % (Minh et al. 2022). Unusual heme-binding motifs were subjected to confirmation by structural prediction using AlphaFold3 (Abramson et al. 2024). Truncation (N or C terminal loss) events were confirmed by retrieving the respective DNA sequence and also the DNA sequence of 5’ and 3’ nearby genes or pseudogenes and translating it into protein using ExPASy translate tool (Gasteiger et al. 2003). All generated trees are reported in the supplementary information and shown in Fig S1-S16, containing the NCBI RefSeq accession codes, the number of heme-binding motifs and the multiple sequence alignments for the transition events of gain/loss of heme-binding motifs. The same phylogenetic trees reporting the species and full taxonomy information at each tip are provided in Supplementary File 6. BLASTp searches (Altschul et al., 1990) against the PDB were performed for the FccA sequences containing no heme-binding motifs on 13/02/2025. Subcellular localisation and presence of signal peptides were predicted using SignalIP 5.0(Almagro Armenteros et al. 2019) and PSORTb version 3.0.3 (Yu et al. 2010). All phylogenetic trees (rooted, species taxonomy, full taxonomy and the corresponding number of hemes) are available in supplementary files.

### 3D structure prediction and analysis

Structures were predicted using the AlphaFold 3 web server (Abramson et al. 2024). Axial ligand conservation was inspected, and its predicted structural implications were analysed when the change in axial ligand coordination resulted in abrupt structural changes regarding the hemecore arrangement. Alternatively, MHC structures were predicted using RoseTTAFold 2 (Baek et al. 2023) within the Google Colaboratory environment with default parameters, except for the increase to the generation of 5 models with 24 recycles each. Heme *c* was incorporated as described in (Soares et al. 2024). Structures were further water refined using the HADDOCK web server (Honorato et al. 2024). The best structures according to the predicted local distance difference test (pLDDT) that did not have overlapping hemes were used. The 33 less mobile atoms of the hemes *c* rings were aligned using the pair_fit function of PyMOL (TM) version 2.5.7. Distances were calculated for the nitrogen and iron atoms of each aligned pair of hemes between the predicted structures and the reference structure of *Shewanella baltica* OS185 (PDB:6R2Q). The structures that fell off the linear regression trend of sequence similarity vs heme-core distance (fig. S19) were discarded and not used for heme-core structural comparison. Confidence intervals at 95% were used from the mean distance values of the predicted structures being analysed for each heme position. Heme-core RMSD distances were performed using the PyMoL pair_fit function used for the alignment. Pairwise sequence similarity was performed using the Needleman–Wunsch algorithm (Needleman and Wunsch 1970) within the EMBOSS Needle webserver (Rice et al. 2000). Signal peptides were removed using Signal peptide IP 5.0 (Almagro Armenteros et al. 2019). All MtrA homolog predicted structures analysed here are available in supplementary files.

## Acknowledgments

Financial support was provided by MOSTMICRO-ITQB unit base funding with references UIDB/04612/2020 and UIDP/04612/2020, LS4FUTURE Associated Laboratory (LA/P/0087/2020), McGEA project (Grant agreement ID: 101183014)

## Data availability

All relevant data are available in the supplementary materials. The heme-binding motif counting script is available in GitHub together with the respective readme file: https://github.com/ricardosoares90/heme_binding_motif_counter

