## supplementary figures for "The modular evolution of multiheme cytochromes *c* bucks the general trend observed in proteins"

### Cytochrome $c_7$

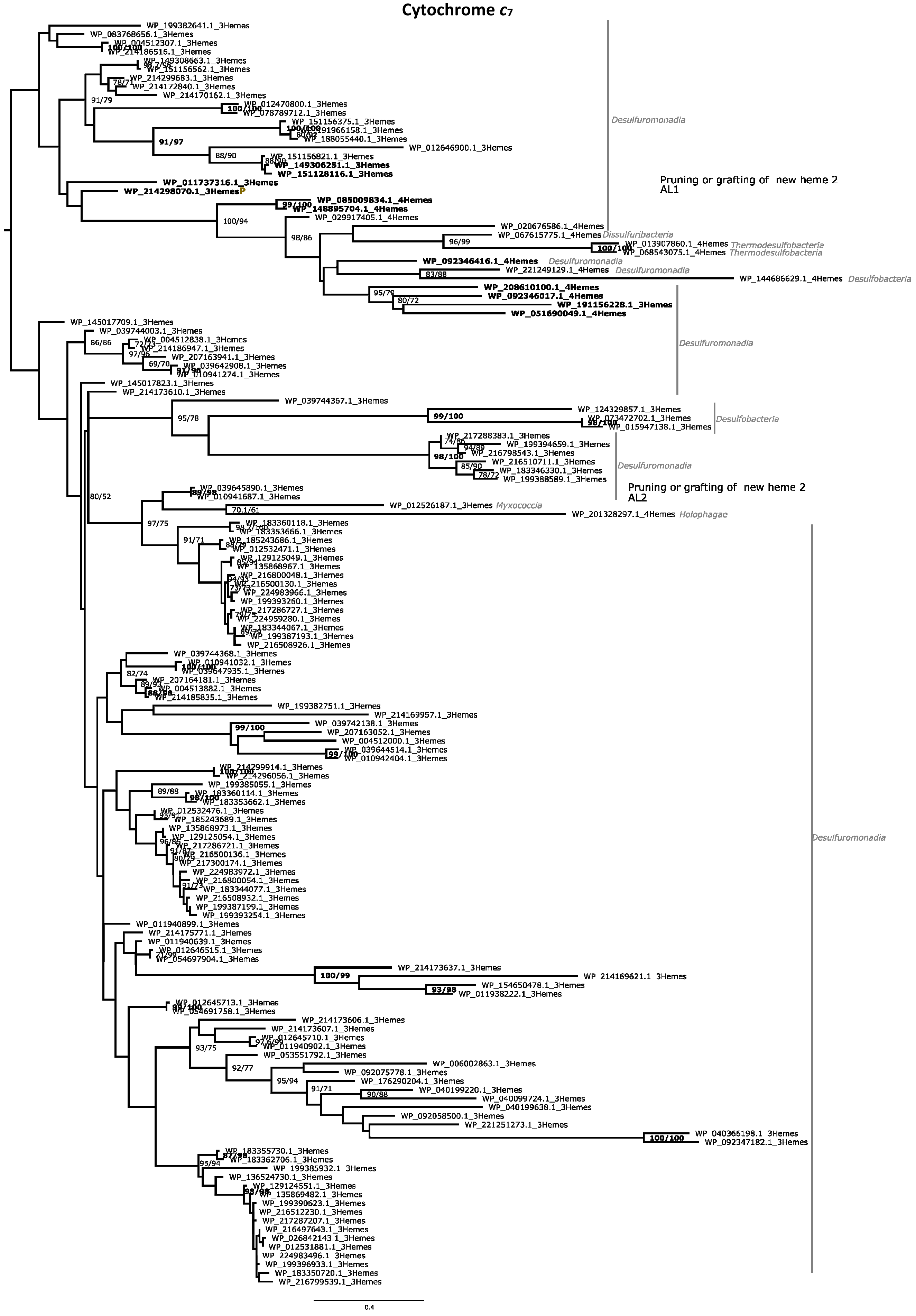

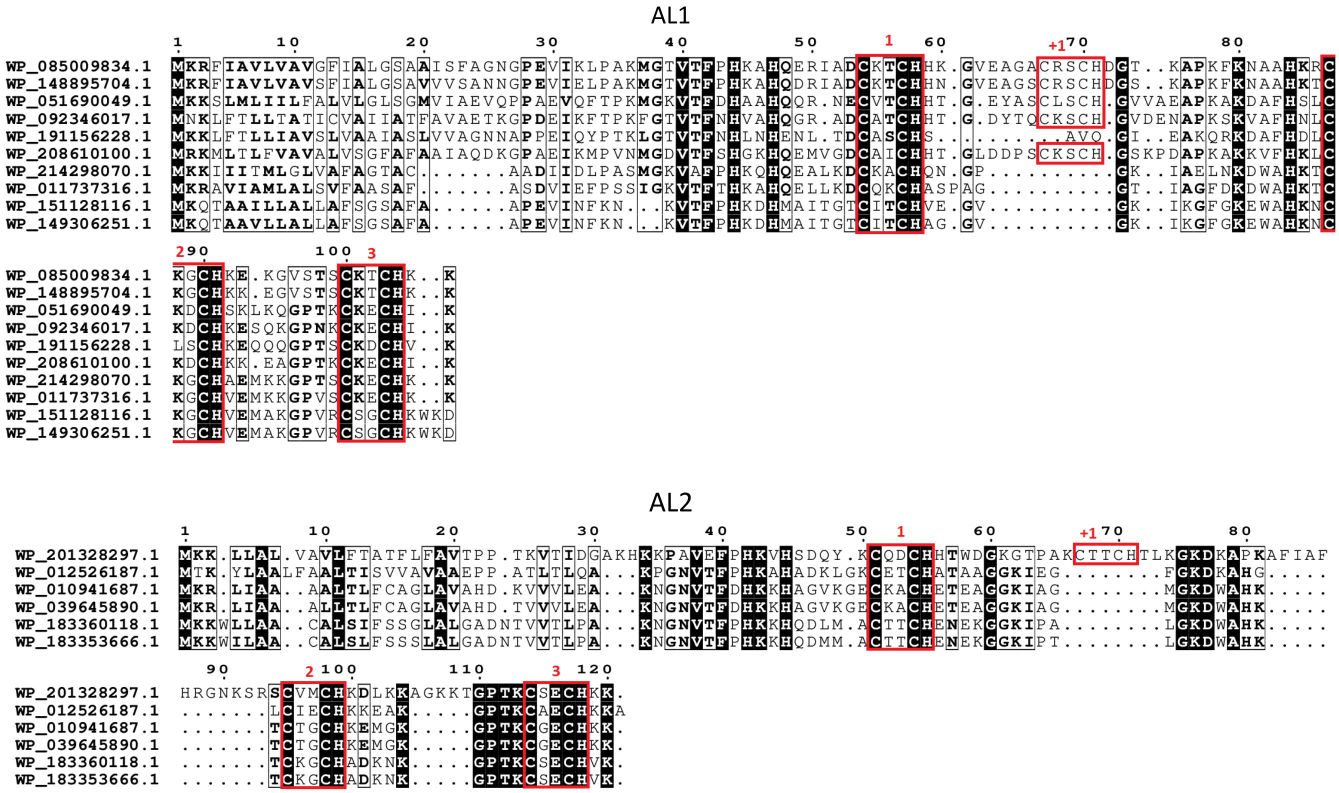


Figure S1. Maximum likelihood phylogenetic tree of cytochrome $c_7$ family. Each presented tip is labeled with the RefSeq accession code, number of heme-binding motifs and if it contains a paralogue within this analysis and taxonomic class. Confidence values (expressed in %) of SH-aLRT/ultrafast bootstrap are presented near each node. Each heme-binding motif gain event was considered when confidence values are ≥ 80%/95%, respectively. For cytochrome $c_7$ only low-confidence events were obtained (highlighted in gray). Confidence values below 70% are not shown. At the bottom of the tree, a subset of the aligned sequences is presented (those that are in bold in the phylogenetic tree) that are related to each event of heme-binding motif gain / loss (AL1 and AL2).

### Cytochrome $c_3$

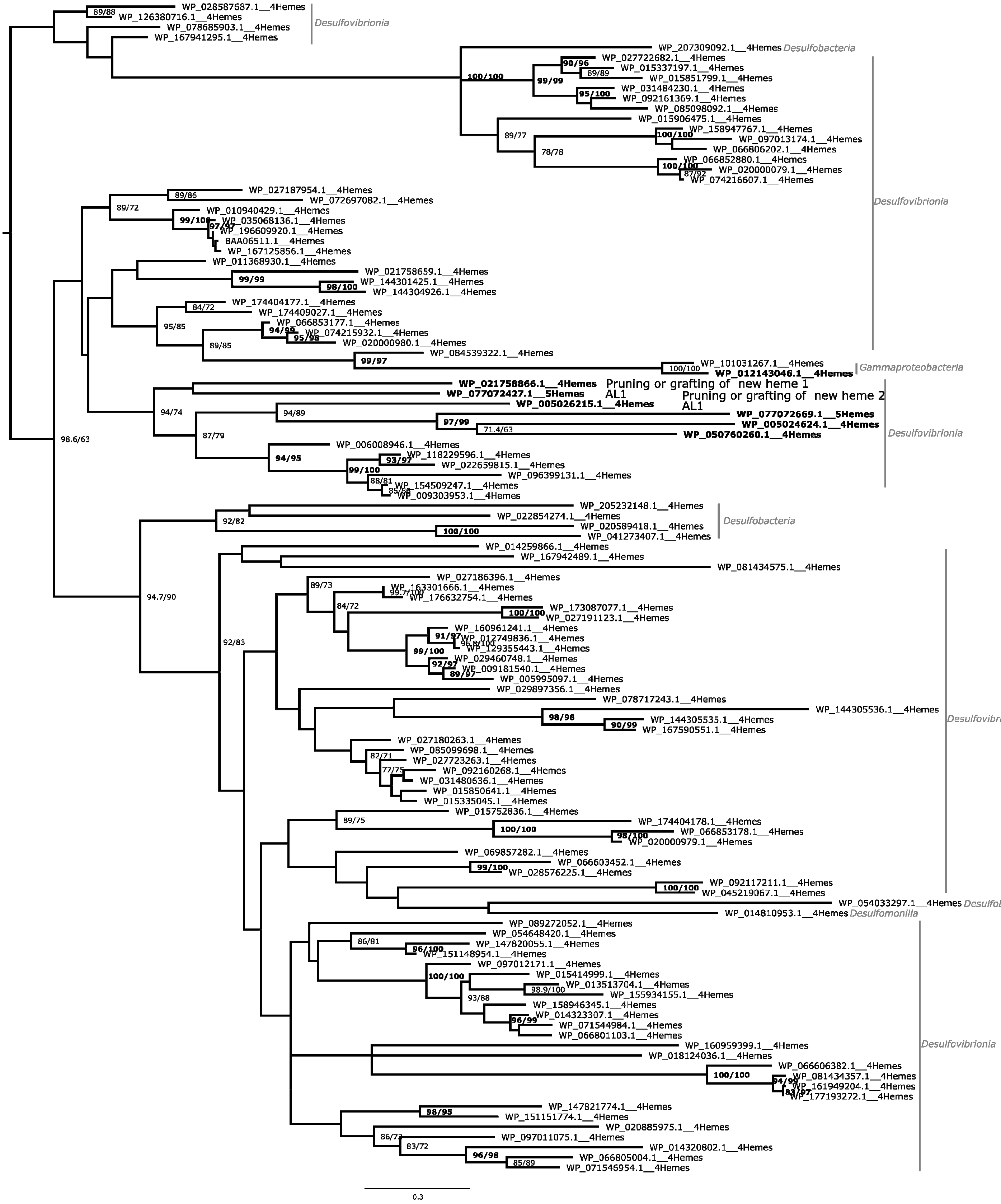

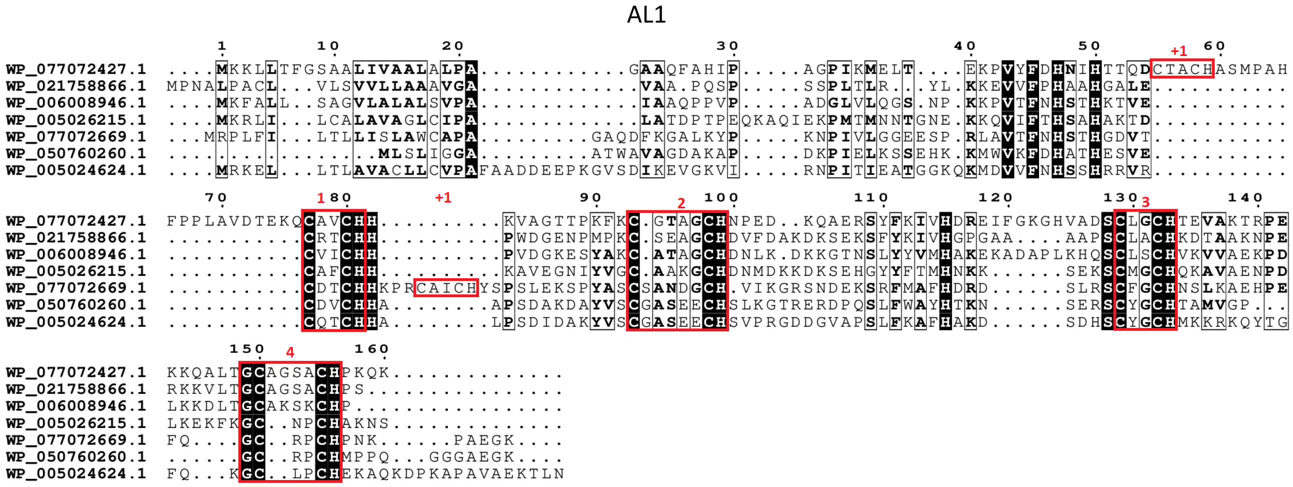


Figure S2. Maximum likelihood phylogenetic tree of cytochrome $c_3$ family. Each presented tip is labeled with the RefSeq accession code, number of heme-binding motifs and if it contains a paralogue within this analysis and taxonomic class. Confidence values (expressed in %) of SH-aLRT/ultrafast bootstrap are presented near each node. Each heme-binding motif gain event was considered when confidence values are ≥ 80%/95%, respectively. For cytochrome $c_3$ only low-confidence events were obtained (highlighted in gray). Confidence values below 70% are not shown. At the bottom of the tree, a subset of the aligned sequences is presented (those that are in bold in the phylogenetic tree) that are related to each event of heme-binding motif gain / loss (AL1).

#### FccA

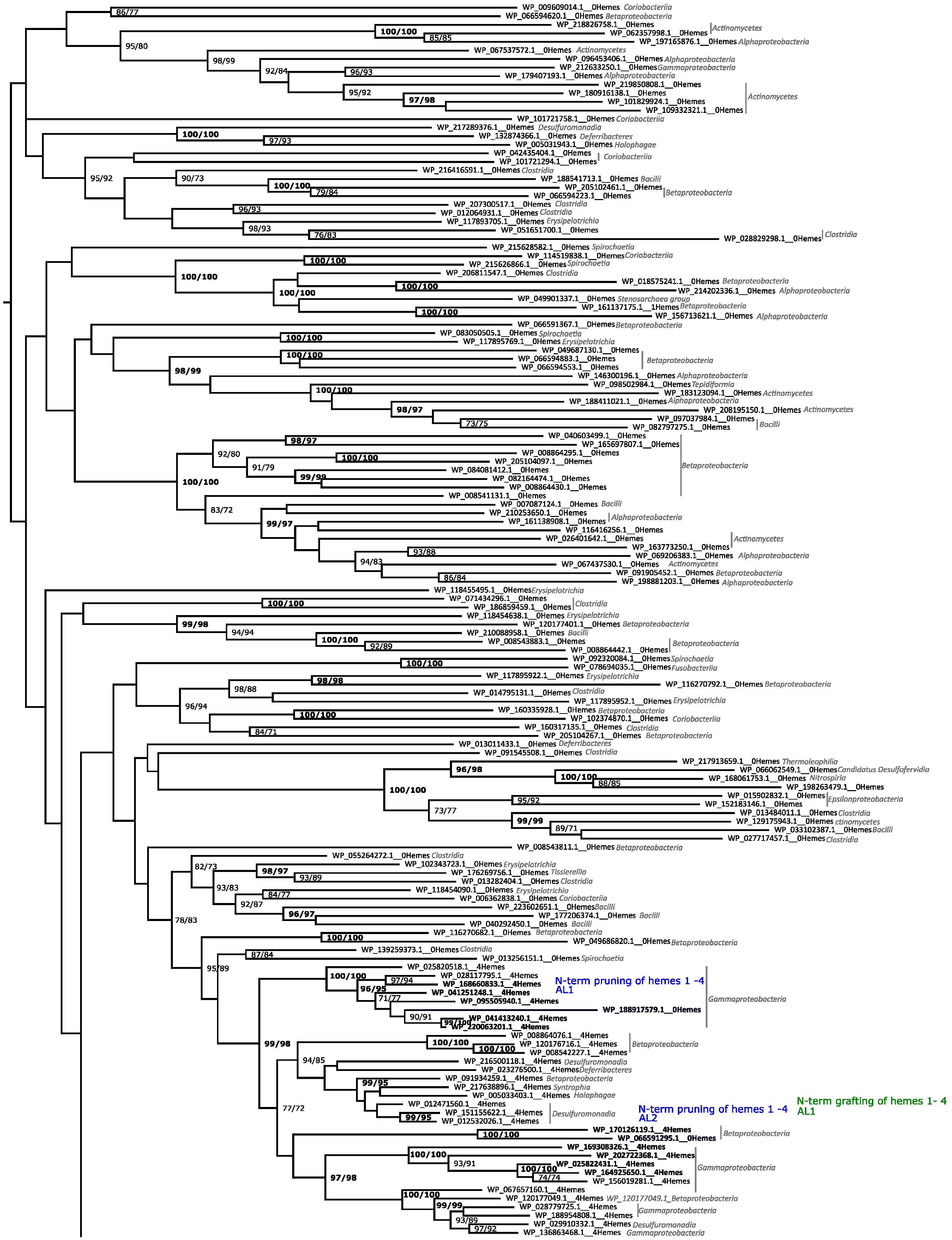


(Continues in next page)

(Continuation)

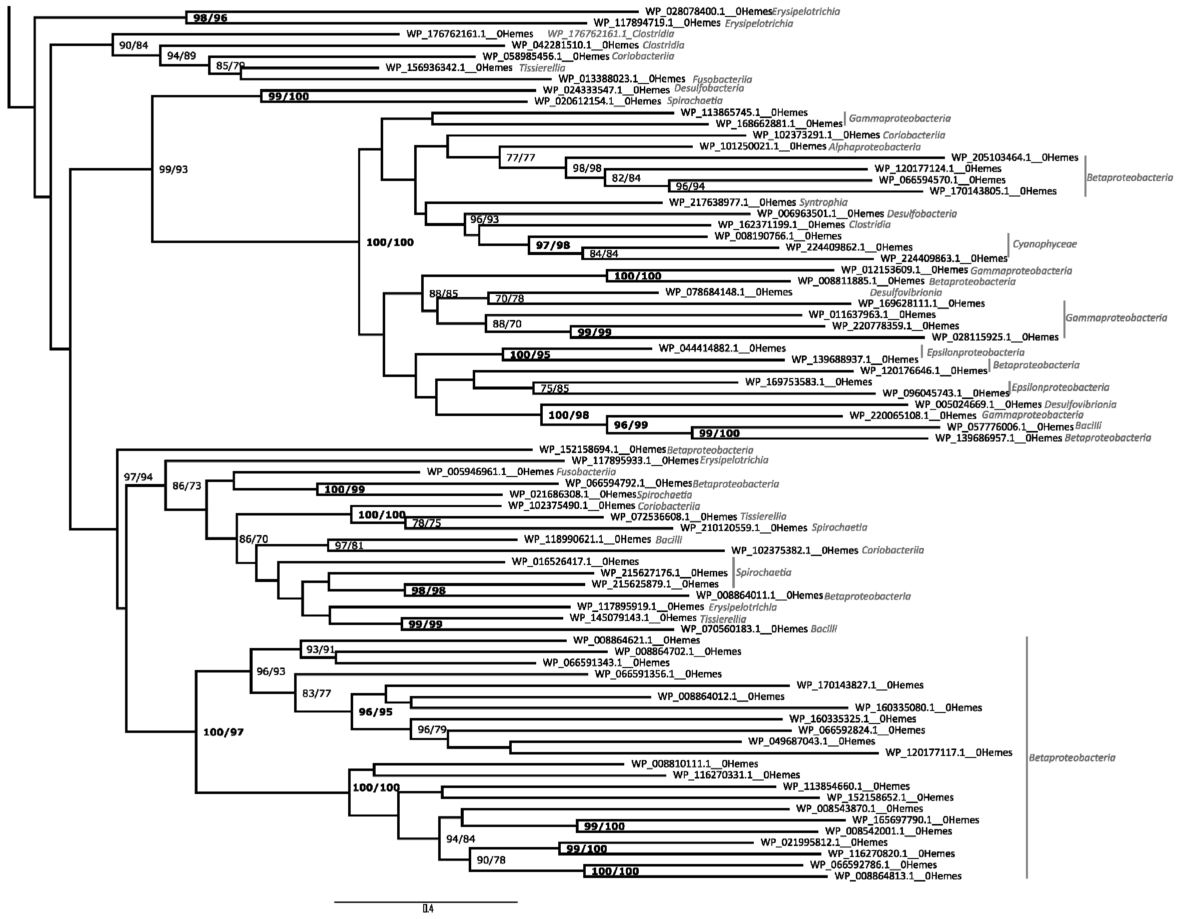

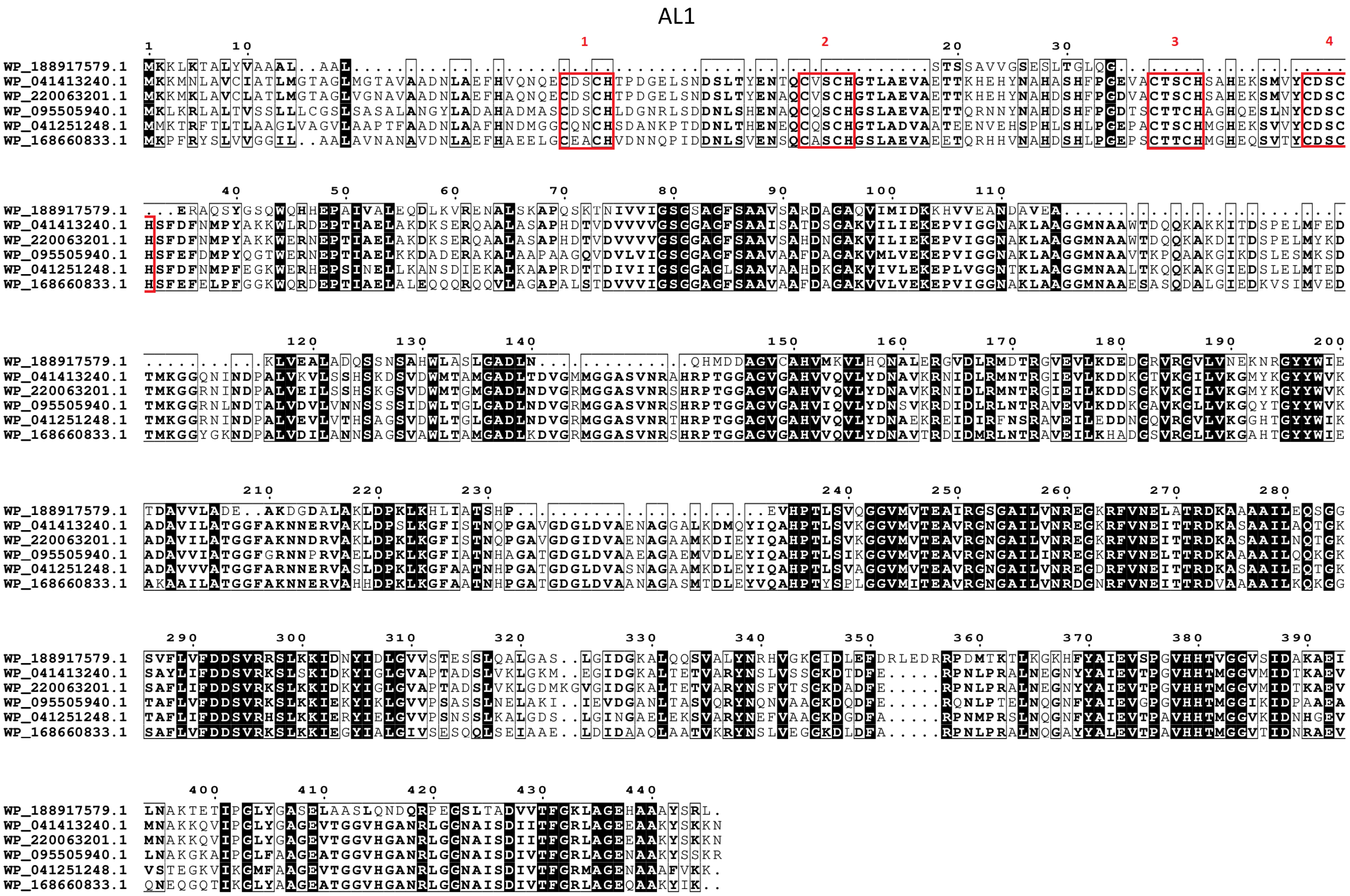


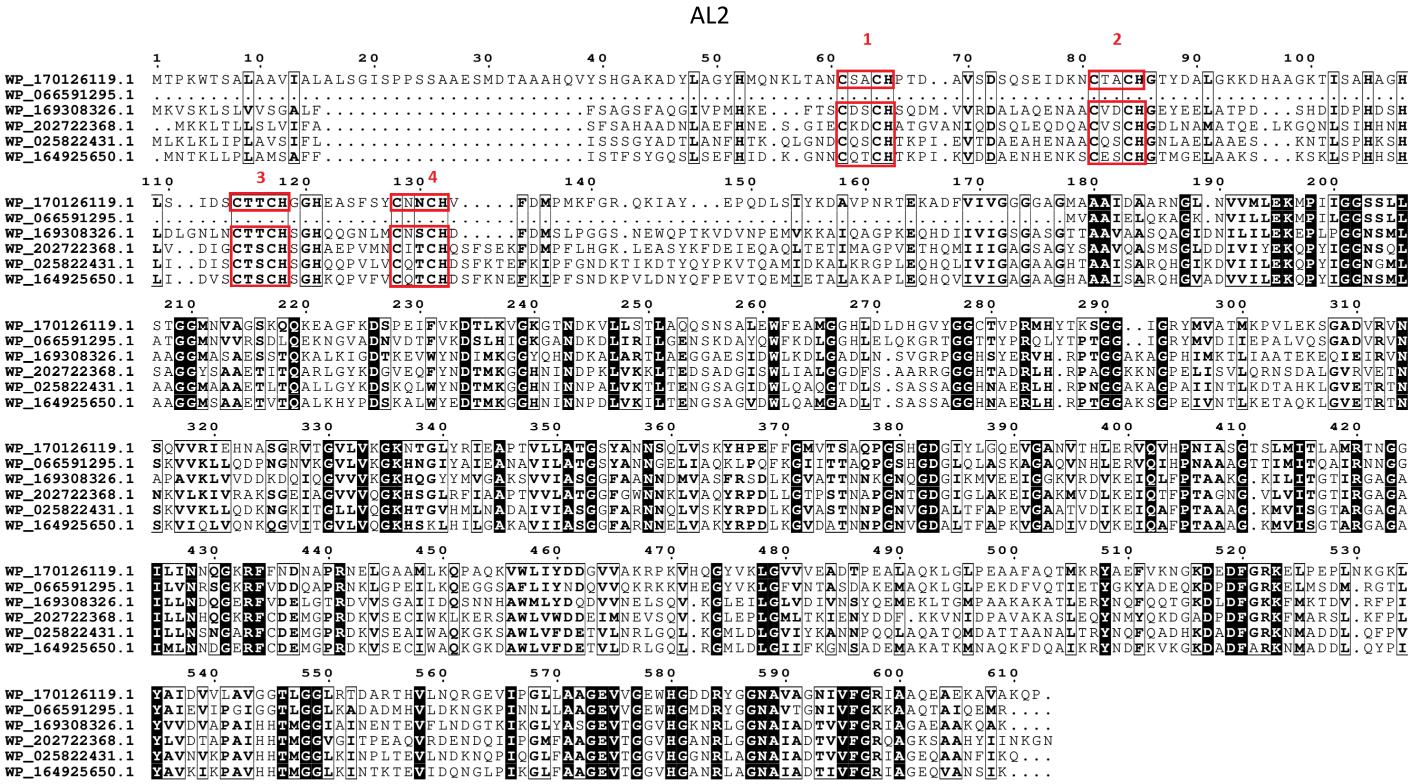


Figure S3. Maximum likelihood phylogenetic tree of cytochrome FccA family. Each presented tip is labeled with the RefSeq accession code, number of heme-binding motifs and if it contains a paralogue within this analysis and taxonomic class. Confidence values (expressed in %) of SH-aLRT/ultrafast bootstrap are presented near each node. Each heme-binding motif gain (green) / loss (blue) event was considered when confidence values are ≥ 80%/95%, respectively, otherwise only highlighted in gray. Confidence values below 70% are not shown. At the bottom of the tree, a subset of the aligned sequences is presented (those that are in bold in the phylogenetic tree) that are related to each event of heme-binding motif gain / loss (AL1).

#### NrfH

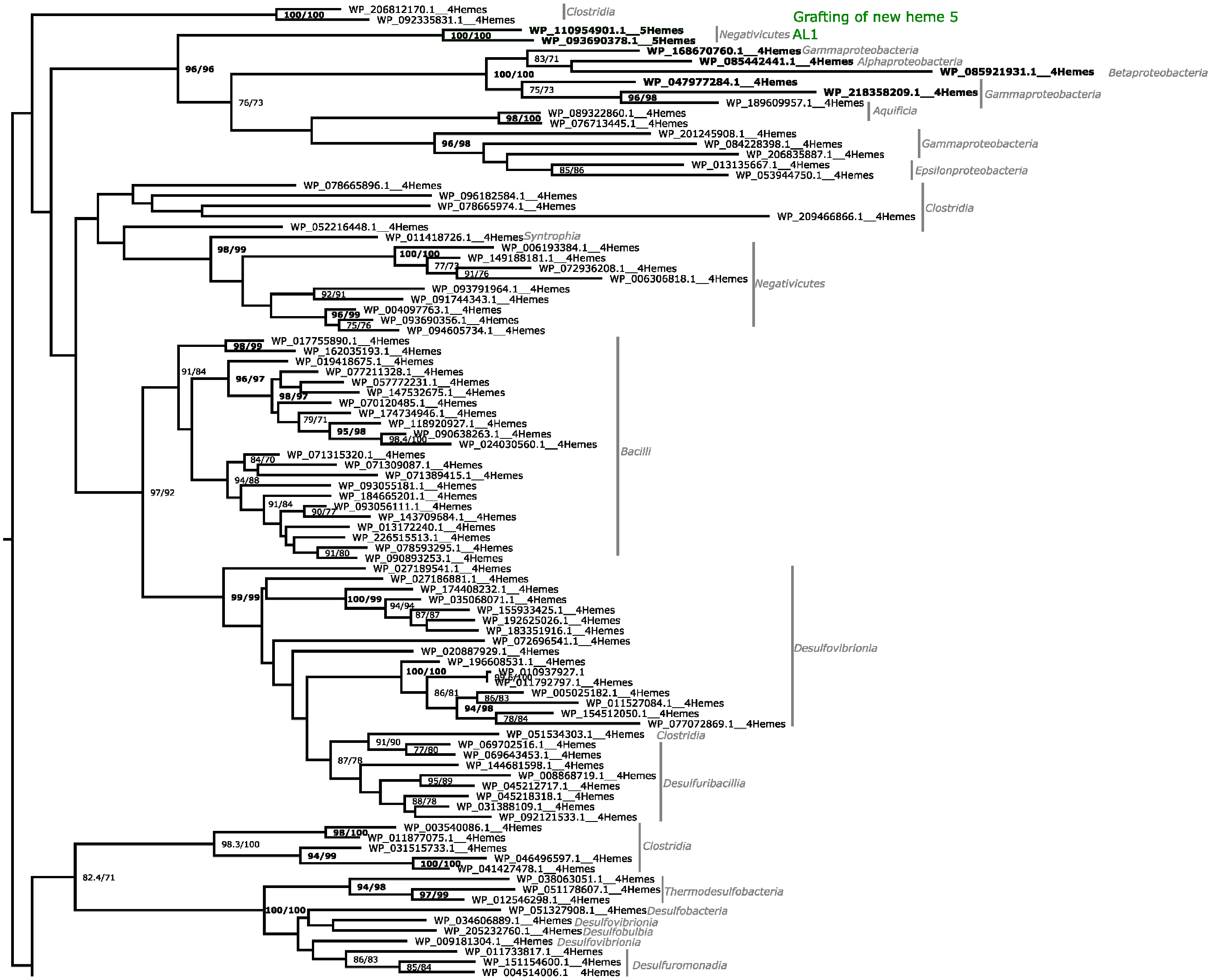


(Continues in next page)

(Continuation)

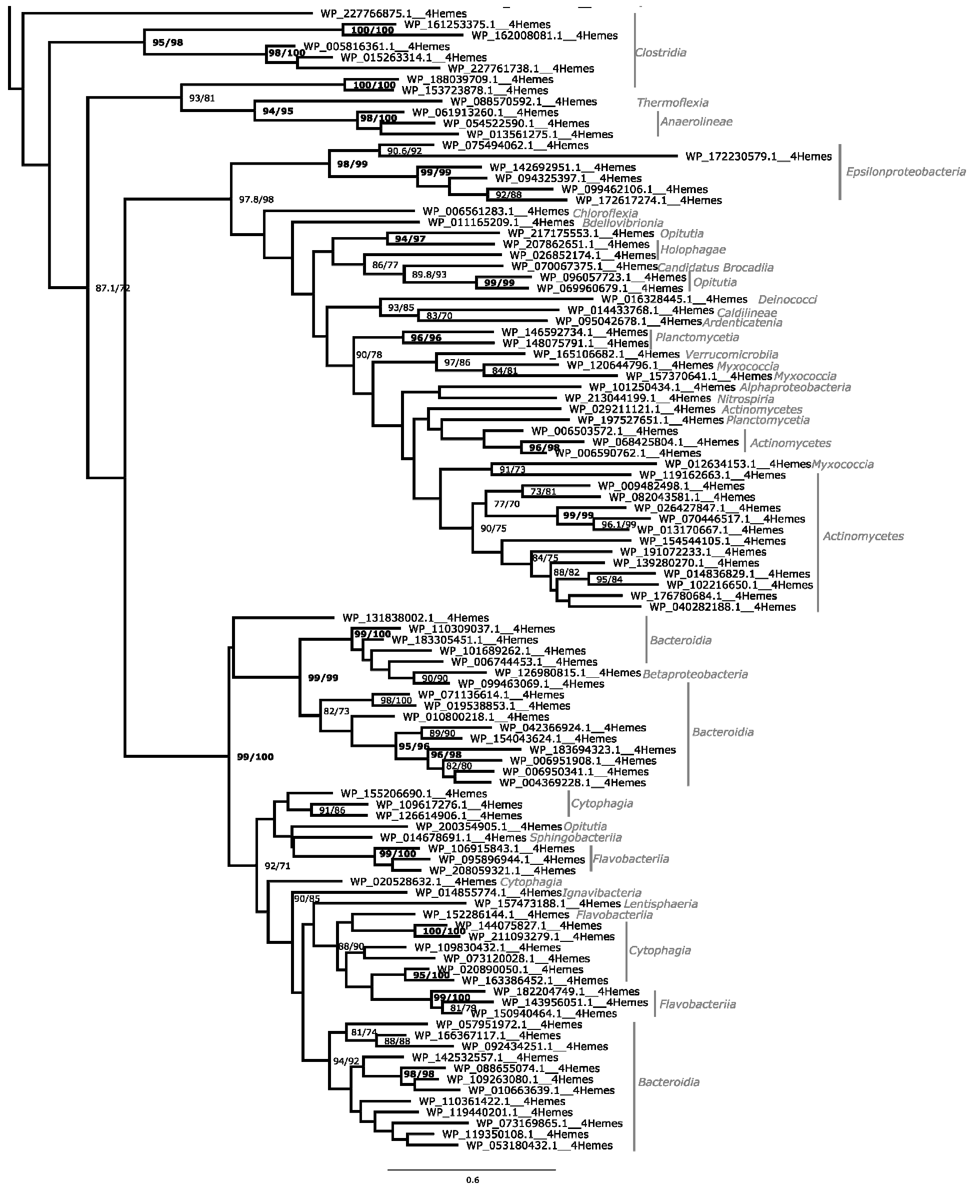

AL1

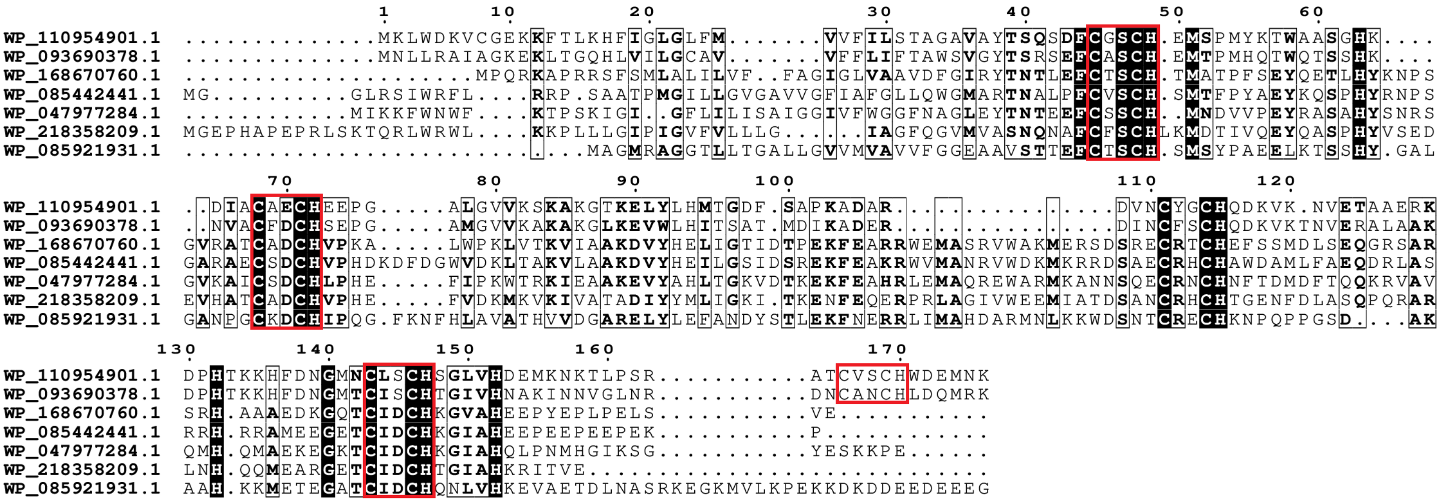


Figure S4. Maximum likelihood phylogenetic tree of cytochrome NrfH family. Each presented tip is labeled with the RefSeq accession code, number of heme-binding motifs and if it contains a paralogue within this analysis and taxonomic class. Confidence values (expressed in %) of SH-aLRT/ultrafast bootstrap are presented near each node. Each heme-binding motif gain (green) / loss (blue) event was considered when confidence values are ≥ 80%/95%, respectively, otherwise only highlighted in gray. Confidence values below 70% are not shown. At the bottom of the tree, a subset of the aligned sequences is presented (those that are in bold in the phylogenetic tree) that are related to each event of heme-binding motif gain / loss (AL1).

**OmcE**

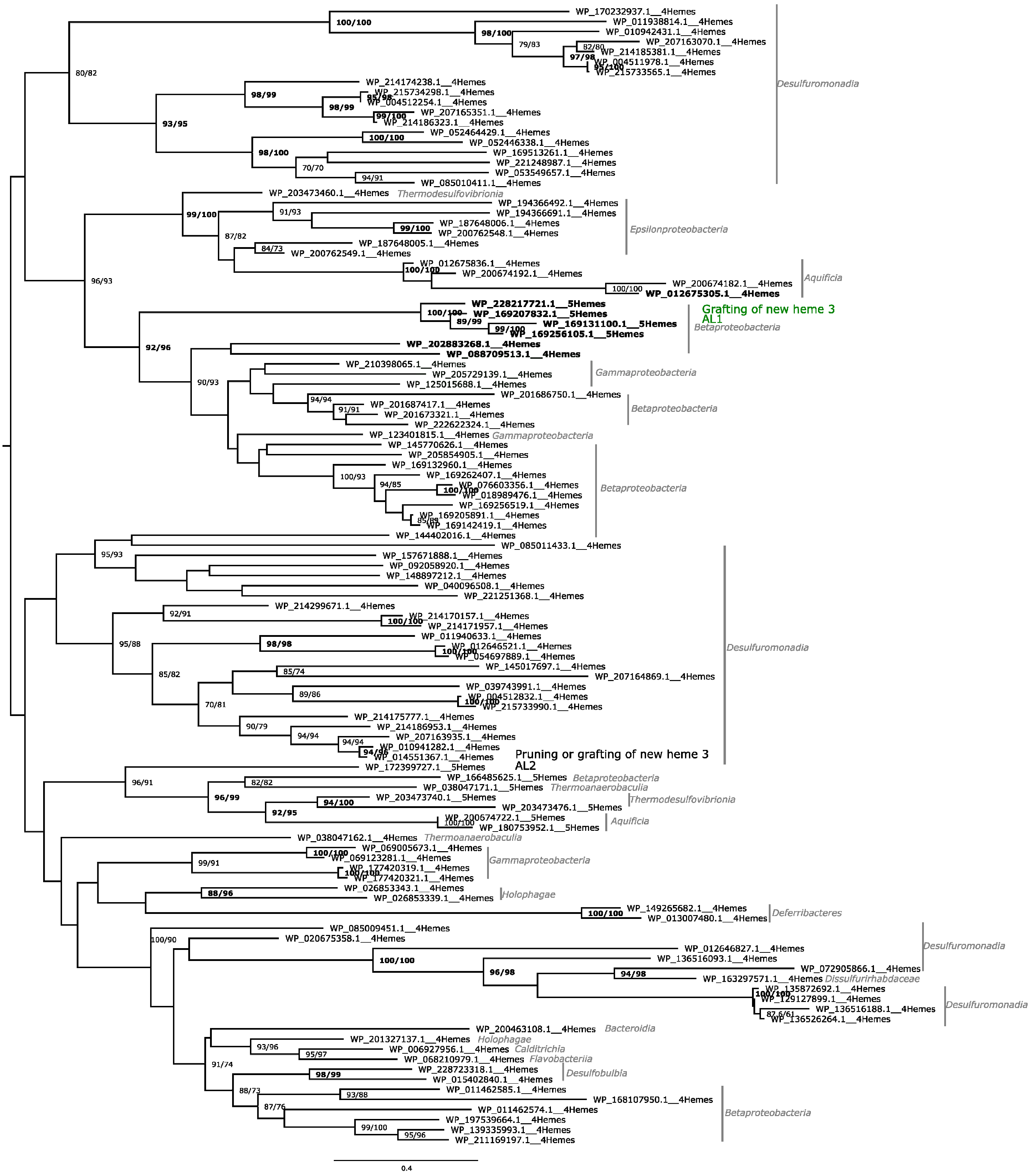
WP_170232937.1__4Hemes
WP_011938814.1__4Hemes
WP_010942431.1__4Hemes
WP_207163070.1__4Hemes
WP_214185381.1__4Hemes
WP_004511978.1__4Hemes
WP_215733565.1__4Hemes
WP_214174238.1__4Hemes
WP_215734298.1__4Hemes
WP_004512254.1__4Hemes
WP_207165351.1__4Hemes
WP_214186323.1__4Hemes
WP_052464429.1__4Hemes
WP_052446338.1__4Hemes
WP_169513261.1__4Hemes
WP_221248987.1__4Hemes
WP_053549657.1__4Hemes
WP_085010411.1__4Hemes
Desulfuromonadia
WP_203473460.1__4Hemes
Thermodesulfovibrionia
WP_194366492.1__4Hemes
WP_194366691.1__4Hemes
WP_187648006.1__4Hemes
WP_200762548.1__4Hemes
WP_187648005.1__4Hemes
WP_200762549.1__4Hemes
Epsilonproteobacteria
WP_012675836.1__4Hemes
WP_200674192.1__4Hemes
WP_200674182.1__4Hemes
WP_012675305.1__4Hemes
Aquificia
WP_228217721.1__5Hemes
WP_169207832.1__5Hemes
WP_169131100.1__5Hemes
WP_169256105.1__5Hemes
Grafting of new heme 3
AL1
Betaproteobacteria
WP_202883268.1__4Hemes
WP_088709513.1__4Hemes
WP_210398065.1__4Hemes
WP_205729139.1__4Hemes
Gammaproteobacteria
WP_125015688.1__4Hemes
WP_201686750.1__4Hemes
WP_201687417.1__4Hemes
WP_201673321.1__4Hemes
WP_222622324.1__4Hemes
Betaproteobacteria
WP_123401815.1__4Hemes
Gammaproteobacteria
WP_145770626.1__4Hemes
WP_205854905.1__4Hemes
WP_169132960.1__4Hemes
WP_169262407.1__4Hemes
WP_076603356.1__4Hemes
WP_018989476.1__4Hemes
WP_169256519.1__4Hemes
WP_169205891.1__4Hemes
WP_169142419.1__4Hemes
Betaproteobacteria
WP_144402016.1__4Hemes
WP_085011433.1__4Hemes
WP_157671888.1__4Hemes
WP_092058920.1__4Hemes
WP_148897212.1__4Hemes
WP_040096508.1__4Hemes
WP_221251368.1__4Hemes
WP_214299671.1__4Hemes
WP_214170157.1__4Hemes
WP_214171957.1__4Hemes
WP_011940633.1__4Hemes
WP_012646521.1__4Hemes
WP_054697889.1__4Hemes
WP_145017697.1__4Hemes
WP_207164869.1__4Hemes
WP_039743991.1__4Hemes
WP_004512832.1__4Hemes
WP_215733990.1__4Hemes
WP_214175777.1__4Hemes
WP_214186953.1__4Hemes
WP_207163935.1__4Hemes
WP_010941282.1__4Hemes
WP_014551367.1__4Hemes
Desulfuromonadia
Pruning or grafting of new heme 3
AL2
WP_172399727.1__5Hemes
WP_166485625.1__5Hemes
Betaproteobacteria
WP_038047171.1__5Hemes
Thermoanaerobaculia
WP_203473740.1__5Hemes
WP_203473476.1__5Hemes
Thermodesulfovibrionia
WP_200674722.1__5Hemes
WP_180753952.1__5Hemes
Aquificia
WP_038047162.1__4Hemes
Thermoanaerobaculia
WP_069005673.1__4Hemes
WP_069123281.1__4Hemes
WP_177420319.1__4Hemes
WP_177420321.1__4Hemes
Gammaproteobacteria
WP_026853343.1__4Hemes
WP_026853339.1__4Hemes
Holophagae
WP_149265682.1__4Hemes
WP_013007480.1__4Hemes
Deferribacteres
WP_085009451.1__4Hemes
WP_020675358.1__4Hemes
WP_012646827.1__4Hemes
WP_136516093.1__4Hemes
WP_072905866.1__4Hemes
Desulfuromonadia
WP_163297571.1__4Hemes
Dissulfurirhabdaceae
WP_135872692.1__4Hemes
WP_129127899.1__4Hemes
WP_136516188.1__4Hemes
WP_136526264.1__4Hemes
Desulfuromonadia
WP_200463108.1__4Hemes
Bacteroidia
WP_201327137.1__4Hemes
Holophagae
WP_006927956.1__4Hemes
Calditrichia
WP_068210979.1__4Hemes
Flavobacteriia
WP_228723318.1__4Hemes
WP_015402840.1__4Hemes
Desulfobulbia
WP_011462585.1__4Hemes
WP_168107950.1__4Hemes
WP_011462574.1__4Hemes
WP_197539664.1__4Hemes
WP_139335993.1__4Hemes
WP_211169197.1__4Hemes
Betaproteobacteria
0.4

## AL1

```
WP_228217721.1  MISRSNKGMHHAMHLTTRALSVLLGSMLSA......GSVQADGDLPSKFSGQGTIGNTRHNMTQRPNS....GSGPAAEMMDFVRNDYNE
WP_169207832.1  ............MKCKISAAFILGLATLAT......GSALAGLDRPTKFSNQGTIGNTRHNLTQRPTD....GSALNAGMMDAYRNDYQQ
WP_169131100.1  ............MLNKIATATIVTIGVLAS......DVAWSGTRSDSEFLQQGTIGNTRHNLTQRQTA....GGGPTGTQMDAVRNNYEQ
WP_169256105.1  ............MTIRIATVAIATLGVLVA......DVAWPGTRSDSEFLNQGTIGNTRHNLSQRQTA....GGGPTGSQMDAVRNNYQQ
WP_202883268.1  MSWSNW...MGRQSLRERGLAVVVAGVVVAVLGTAINWPASAQSTGLAARKGSGISTTKHNLGSSGT.....GNNTIAAGTGAAASPSTE
WP_210398065.1  .........MIKSGKNKLALAIVGAAALAV.................AGLGTAGIKNTRHNLGSSAA.....SNGPDNNRTDA....TGE
WP_125015688.1  .........MFKSMTKNIGLAALGGALLVA.................SAATFAGISGTKHNLGS........GGVVGNNLFDG....TAE
WP_205729139.1  .........MKRALVGMICLTLSGGTL.......................AASIVDSKHNLGNIG......GNTANRNYTDA....TQE
WP_088709513.1  MGTQ.....FGKGLAQVFHRATYAAVLAVG.............ALGLMQGADAQVVDTKHNLSATAPAAGSQGATRVNYYTGAGA..TTQ

WP_228217721.1  ICVYCHTPHAANTT....IAAPLWNRVVRA...TTYTLYNQQ...........TLSGETTQPGPNSLTCLSCHDGQTAIDSIVNMPGTGP
WP_169207832.1  VCVYCHTPHAANTN....ISAPLWNRTVRT...TTYTLYNQQ...........TLSQDATQPGPNSLTCLSCHDGQTAVDSIVNMPGSG.
WP_169131100.1  VCVYCHTPHGANQN....IKAPLWNRTVRT...TTYTLYSQL...........SLSQDATQPGPSSLTCLSCHDGQTAVDSIINMPGSG.
WP_169256105.1  VCVYCHTPHAANQN....ISAPLWNRTVRS...TTYTLYNQQ...........SLSQEATQPGPNSLTCLSCHDGQTAVDSIVNMPGSG.
WP_202883268.1  ICVFCHTPHAAEV.....ARGPLWNRLAGT...ATITPYTSATLTGGTVS...GGMGTALTPGSVSLACLSCHDGTQAMNTLINAPGSN.
WP_210398065.1  ICVFCHTPHGADNA....AAVPLWNRALESDAANTYTTYDQL.....GTS...SLDGGVVAVGSVSIACLSCHDGQQAMDVMINEPGSG.
WP_125015688.1  ICVFCHTPHGSDTS....APVPLWNRNINSS..PTYTTYATL.....QTS...SLQGKEAAIGSVSLACLSCHDGTQAMNSVINAPGSG.
WP_205729139.1  VCVFCHTPHGGSATHNGEAVAPLWNKAVTT...STFQTYDSL.....GTS...SLKGSVANVGSVSIACLSCHDGSQAMDVMINQPDSG.
WP_088709513.1  VCVFCHTPHGAAS.....AAAPLWQRVVKAD..NYYQTYGSV.....NASPTMEADQANVTVGSVSLACLSCHDGTQAINTMANKPGSG.

WP_228217721.1  DRYWTDMATSPS....NTFLNGWPKSARAVVDATSHNSL.....SKTGCLSCHNPGTEIGAPDFSAAAIGTDLSNDHPVGVKLPSGSDWN
WP_169207832.1  .GYNGALALDPSAATAETLLDTWRRNG.GDPDAVTHQGLNSDPNSETSCLSCHSPGKTAAATNFQVFAIGTDLTNDHPVGVKLPAGSDWN
WP_169131100.1  .KYNAALALNPDAATTKTLLDAWSTTD....DAFDHYALK...DNNTECLSCHSPSGQGGAQDFSAYALGTDLRNDHPVGVKLPTGAEWN
WP_169256105.1  .KYNAALALNPDTSTTKALLDAWSTTD....DAFDHFGLK...DNNTECLSCHSPGGQGGAQDFSAFAIGTDLRNDHPVGVKLPSGSDWN
WP_202883268.1  .GYNAGGA........AFGTLVSGAG............................LIT....GVAKIGGDLTNDHPIGMKYAGGRADT
WP_210398065.1  .TDTIS............AGTWSGST...SDGLTG...................GDTMNDANFGITFIGTDLTNDHPIGIQYAGGGESG
WP_125015688.1  .GYNATGS.........PMAGTWDGTT...VGSAN....................GLLLD....GITKIGTDLRDDHPISIQYAGGGFLT
WP_205729139.1  .GYTAAGA........SLGGTWTANG..AVDAAT....................GKMLTAN..PIPGVGTDLRNDHPIGIQYAGGLISG
WP_088709513.1  .NNALGGD........WAAGNNLTNS..GLSTTS....................MANLGGG..TVGTDGKDLQNDHPIGIKYAGGFTYS

WP_228217721.1  VPGG.....IRGNMKYFDRD.................................................GNNRPSKDEIRFF.........
WP_169207832.1  VPTG.....TYQNIRFFDKD.................................................GDNRPDKDELRFY.........
WP_169131100.1  VPSG.....TYQNVRFFDTD.................................................GDNRPDKNELRFY.........
WP_169256105.1  VPSG.....TYKNVRFFDTD.................................................GDSRPDKEELRFY.........
WP_202883268.1  VQEV............GGYVQFTKGTINATDAWWVDTDTIGRTGAPA......................ADSSRQKTDMWLYART......
WP_210398065.1  SRGD....GEGTVSNYADTDFKPAWKGTVNNMPIWWVPTVAATGSSDIGDVDGNGGTKASGASRTAGTAPTSRNKTDMQLYTRLLVSQGG
WP_125015688.1  TSCTGGDGALCSNTMLSTSGFNLPRMAIANSNPVWWVDNAQGAG.........................TDNARNKEDIQLYTRTDLS...
WP_205729139.1  TDHT.......NASNYRNPDFVAAAEVTINGNQIWYVPVPASNATTGLAN.........SVSGSQISPVAGTRNKTDMALYTRSVAT...
WP_088709513.1  VSNG.....VVTNNAFADTAFRSTSSTVVNNVPVWYVEKAAG.........................NVAGRDKNDIQLYTRST.....

WP_228217721.1  ...DSG......QGYEVECASCHDPHGVPGTGGTFLPTFLRVTADGSQICLTCHVK
WP_169207832.1  ...DSG......NGYEVECASCHDPHGVPGTTSTFLPTFLRVTADGSQICMTCHVK
WP_169131100.1  ...DSG......QGFEVECASCHDPHGVPGATAKFLPTFLRVSVDGSQICLTCHVK
WP_169256105.1  ...DSG......QGYEVECASCHDPHGVPGATASFLPTFLRVTADGSQICLTCHVK
WP_202883268.1  ...GTT......DTAYVECASCHDPHTDN........ALFLRIPNTYSSVCLSCHIK
WP_210398065.1  GIFDSGTDGAADAEPYVECASCHDPHTEN........ATFLRVDNFGSGVCLACHTK
WP_125015688.1  ..GDSN......AQPFVECASCHDPHTSN........TTFLRIENTGSNVCLSCHDK
WP_205729139.1  ..IDSGR.....AQPYVECASCHDPHQANT.......ATFLRIDNTGSAVCLACHKK
WP_088709513.1  ..GGGN......YQPYVECASCHDPHANA........ELFMRVPVAGSEICLTCHIK
```

**AL2**

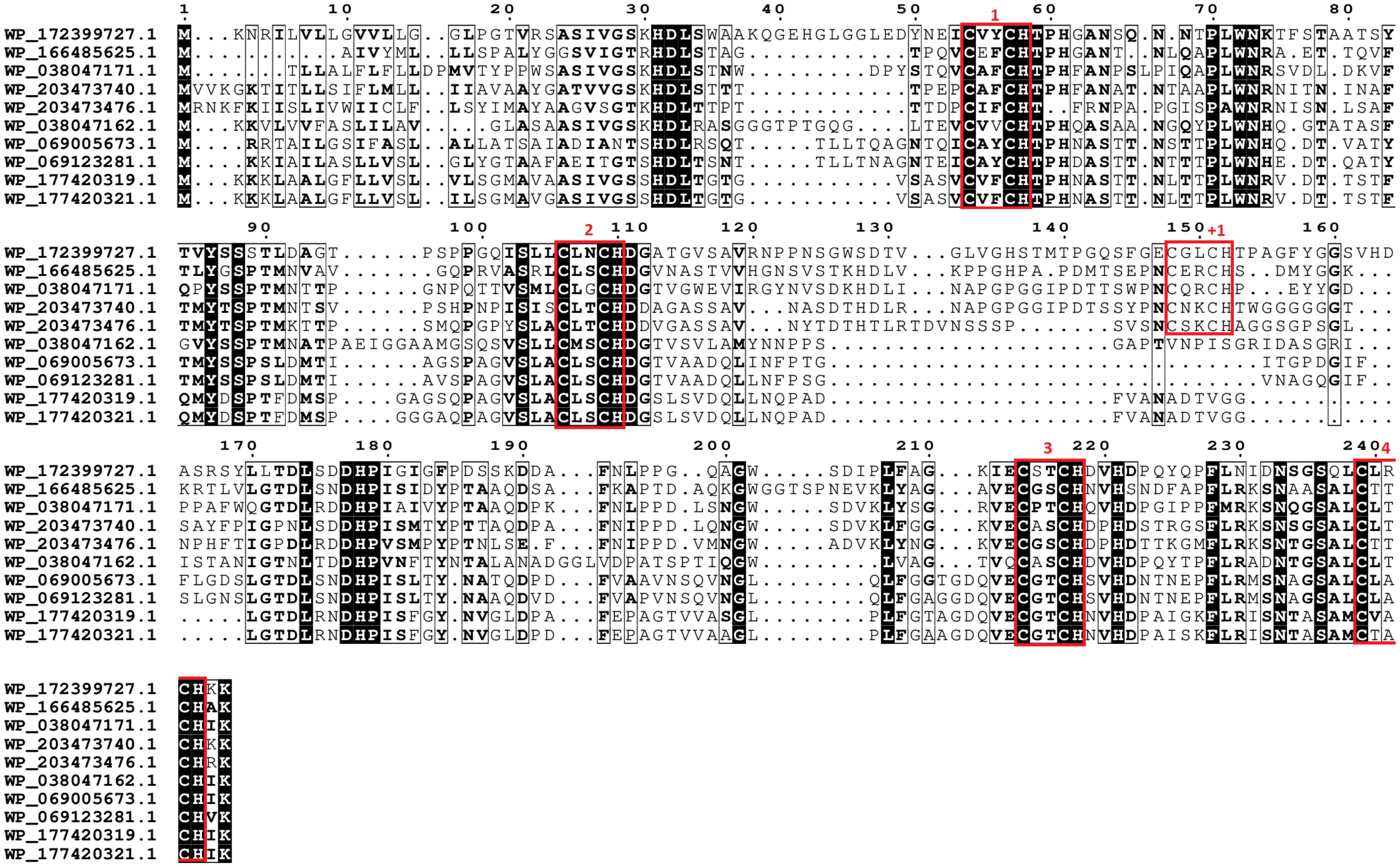


Figure S5. Maximum likelihood phylogenetic tree of cytochrome OmcE family. Each presented tip is labeled with the RefSeq accession code, number of heme-binding motifs and if it contains a paralogue within this analysis and taxonomic class. Confidence values (expressed in %) of SH-aLRT/ultrafast bootstrap are presented near each node. Each heme-binding motif gain (green) / loss (blue) event was considered when confidence values are ≥ 80%/95%, respectively, otherwise only highlighted in gray. Confidence values below 70% are not shown. At the bottom of the tree, a subset of the aligned sequences is presented (those that are in bold in the phylogenetic tree) that are related to each event of heme-binding motif gain / loss (AL1 and AL2).

### PufC

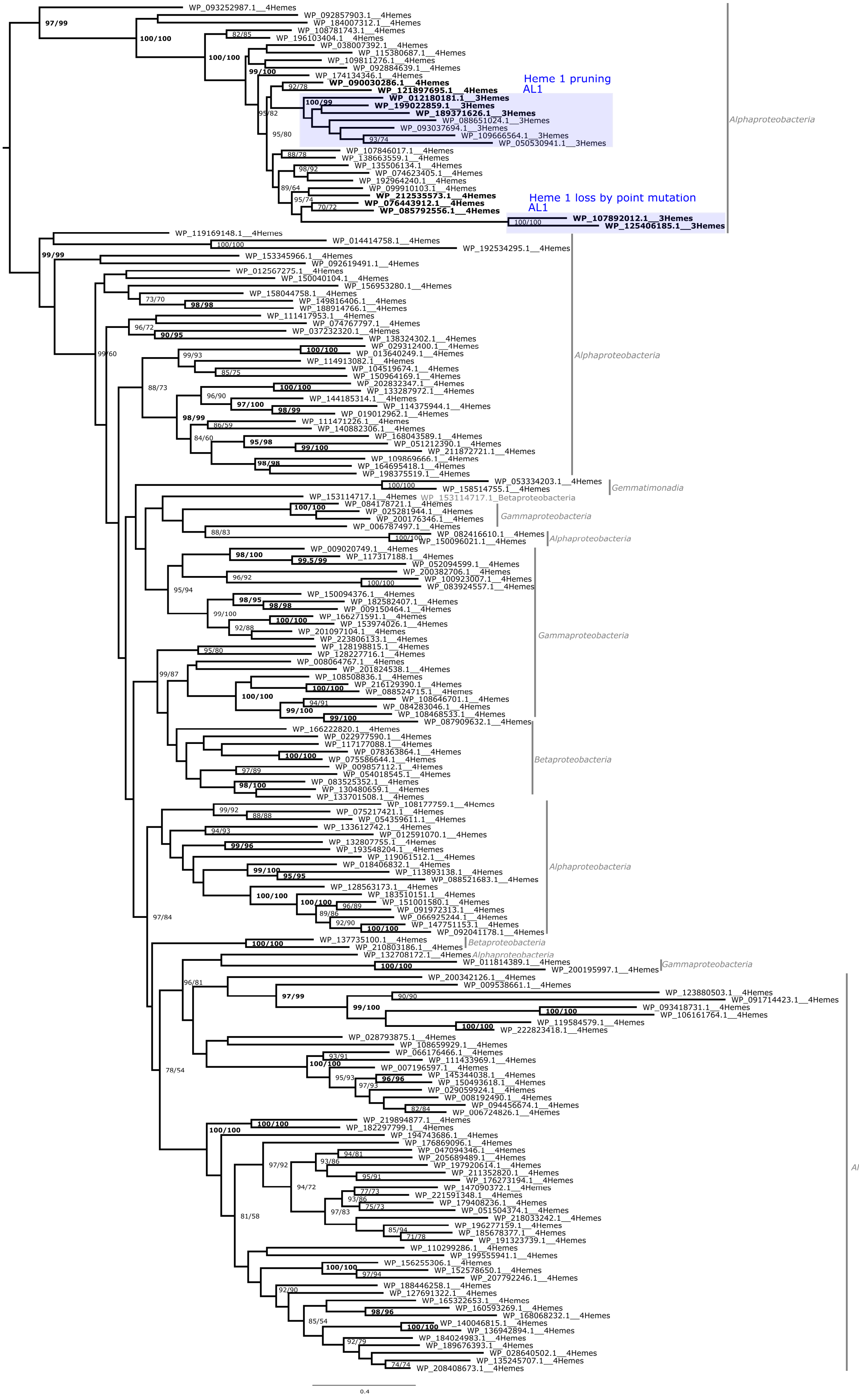

WP_093252987.1__4Hemes
97/99
WP_092857903.1__4Hemes
WP_184007312.1__4Hemes
100/100
WP_108781743.1__4Hemes
82/85
WP_196103404.1__4Hemes
WP_038007392.1__4Hemes
100/100
WP_115380687.1__4Hemes
WP_109811276.1__4Hemes
WP_092884639.1__4Hemes
99/100
WP_174134346.1__4Hemes
92/78
WP_090030286.1__4Hemes
WP_121897695.1__4Hemes
Heme 1 pruning
AL1
WP_012180181.1__3Hemes
100/99
WP_199022859.1__3Hemes
WP_189371626.1__3Hemes
95/82
WP_088651024.1__3Hemes
WP_093037694.1__3Hemes
95/80
WP_109666564.1__3Hemes
93/74
WP_050530941.1__3Hemes
WP_107846017.1__4Hemes
88/78
WP_138663559.1__4Hemes
98/92
WP_135506134.1__4Hemes
WP_074623405.1__4Hemes
WP_192964240.1__4Hemes
89/64
WP_099910103.1__4Hemes
WP_212535573.1__4Hemes
95/74
WP_076443912.1__4Hemes
70/72
WP_085792556.1__4Hemes
Heme 1 loss by point mutation
AL1
WP_107892012.1__3Hemes
100/100
WP_125406185.1__3Hemes
Alphaproteobacteria
WP_119169148.1__4Hemes
WP_014414758.1__4Hemes
100/100
WP_192534295.1__4Hemes
99/99
WP_153345966.1__4Hemes
WP_092619491.1__4Hemes
WP_012567275.1__4Hemes
WP_150040104.1__4Hemes
WP_156953280.1__4Hemes
WP_158044758.1__4Hemes
73/70
WP_149816406.1__4Hemes
98/98
WP_188914766.1__4Hemes
WP_111417953.1__4Hemes
96/72
WP_074767797.1__4Hemes
90/95
WP_037232320.1__4Hemes
WP_138324302.1__4Hemes
99/60
WP_029312400.1__4Hemes
100/100
WP_013640249.1__4Hemes
99/93
WP_114913082.1__4Hemes
WP_104519674.1__4Hemes
85/75
WP_150964169.1__4Hemes
88/73
WP_202832347.1__4Hemes
100/100
WP_133287972.1__4Hemes
96/90
WP_144185314.1__4Hemes
97/100
WP_114375944.1__4Hemes
98/99
WP_019012962.1__4Hemes
98/99
WP_111471226.1__4Hemes
86/59
WP_140882306.1__4Hemes
84/60
WP_168043589.1__4Hemes
95/98
WP_051212390.1__4Hemes
99/100
WP_211872721.1__4Hemes
WP_109869666.1__4Hemes
98/98
WP_164695418.1__4Hemes
WP_198375519.1__4Hemes
Alphaproteobacteria
WP_053334203.1__4Hemes
100/100
WP_158514755.1__4Hemes
Gemmatimonadia
WP_153114717.1__4Hemes
WP_153114717.1_Betaproteobacteria
WP_084178721.1__4Hemes
100/100
WP_025281944.1__4Hemes
WP_200176346.1__4Hemes
WP_006787497.1__4Hemes
Gammaproteobacteria
88/83
WP_082416610.1__4Hemes
100/100
WP_150096021.1__4Hemes
Alphaproteobacteria
WP_009020749.1__4Hemes
98/100
WP_117317188.1__4Hemes
99.5/99
WP_052094599.1__4Hemes
WP_200382706.1__4Hemes
96/92
WP_100923007.1__4Hemes
100/100
WP_083924557.1__4Hemes
95/94
WP_150094376.1__4Hemes
98/95
WP_182582407.1__4Hemes
98/98
WP_009150464.1__4Hemes
99/100
WP_166271591.1__4Hemes
100/100
WP_153974026.1__4Hemes
92/88
WP_201097104.1__4Hemes
WP_223806133.1__4Hemes
WP_128198815.1__4Hemes
95/80
WP_128227716.1__4Hemes
WP_008064767.1__4Hemes
99/87
WP_201824538.1__4Hemes
WP_108508836.1__4Hemes
WP_216129390.1__4Hemes
100/100
WP_088524715.1__4Hemes
100/100
WP_108646701.1__4Hemes
94/91
WP_084283046.1__4Hemes
99/100
WP_108468533.1__4Hemes
99/100
WP_087909632.1__4Hemes
Gammaproteobacteria
WP_166222820.1__4Hemes
WP_022977590.1__4Hemes
WP_117177088.1__4Hemes
WP_078363864.1__4Hemes
100/100
WP_075586644.1__4Hemes
WP_009857112.1__4Hemes
97/89
WP_054018545.1__4Hemes
WP_083525352.1__4Hemes
98/100
WP_130480659.1__4Hemes
WP_133701508.1__4Hemes
Betaproteobacteria
WP_108177759.1__4Hemes
99/92
WP_075217421.1__4Hemes
88/88
WP_054359611.1__4Hemes
WP_133612742.1__4Hemes
94/93
WP_012591070.1__4Hemes
WP_132807755.1__4Hemes
99/96
WP_193548204.1__4Hemes
WP_119061512.1__4Hemes
WP_018406832.1__4Hemes
99/100
WP_113893138.1__4Hemes
95/95
WP_088521683.1__4Hemes
WP_128563173.1__4Hemes
100/100
WP_183510151.1__4Hemes
100/100
WP_151001580.1__4Hemes
96/89
WP_091972313.1__4Hemes
89/86
WP_066925244.1__4Hemes
92/90
WP_147751153.1__4Hemes
100/100
WP_092041178.1__4Hemes
Alphaproteobacteria
97/84
WP_137735100.1__4Hemes
100/100
WP_210803186.1__4Hemes
Betaproteobacteria
WP_132708172.1__4Hemes
Alphaproteobacteria
WP_011814389.1__4Hemes
100/100
WP_200195997.1__4Hemes
Gammaproteobacteria
WP_200342126.1__4Hemes
96/81
WP_009538661.1__4Hemes
WP_123880503.1__4Hemes
90/90
WP_091714423.1__4Hemes
97/99
WP_093418731.1__4Hemes
99/100
100/100
WP_106161764.1__4Hemes
WP_119584579.1__4Hemes
100/100
WP_222823418.1__4Hemes
WP_028793875.1__4Hemes
78/54
WP_108659929.1__4Hemes
WP_066176466.1__4Hemes
93/91
WP_111433969.1__4Hemes
100/100
WP_007196597.1__4Hemes
95/93
WP_145344038.1__4Hemes
96/96
WP_150493618.1__4Hemes
97/93
WP_029059924.1__4Hemes
WP_008192490.1__4Hemes
WP_094456674.1__4Hemes
82/84
WP_006724826.1__4Hemes
WP_219894877.1__4Hemes
100/100
WP_182297799.1__4Hemes
100/100
WP_194743686.1__4Hemes
WP_176869096.1__4Hemes
94/81
WP_047094346.1__4Hemes
93/86
WP_205689489.1__4Hemes
97/92
WP_197920614.1__4Hemes
WP_211352820.1__4Hemes
95/91
WP_176273194.1__4Hemes
94/72
WP_147090372.1__4Hemes
77/73
WP_221591348.1__4Hemes
93/86
WP_179408236.1__4Hemes
75/73
WP_051504374.1__4Hemes
97/83
WP_218033242.1__4Hemes
81/58
WP_196277159.1__4Hemes
85/94
WP_185678377.1__4Hemes
71/78
WP_191323739.1__4Hemes
WP_110299286.1__4Hemes
WP_199555941.1__4Hemes
WP_156255306.1__4Hemes
100/100
WP_152578650.1__4Hemes
97/94
WP_207792246.1__4Hemes
WP_188446258.1__4Hemes
92/90
WP_127691322.1__4Hemes
WP_165322653.1__4Hemes
WP_160593269.1__4Hemes
98/96
WP_168068232.1__4Hemes
85/54
WP_140046815.1__4Hemes
100/100
WP_136942894.1__4Hemes
WP_184024983.1__4Hemes
92/79
WP_189676393.1__4Hemes
WP_028640502.1__4Hemes
WP_135245707.1__4Hemes
74/74
WP_208408673.1__4Hemes
Alphaproteobacteria
0.4

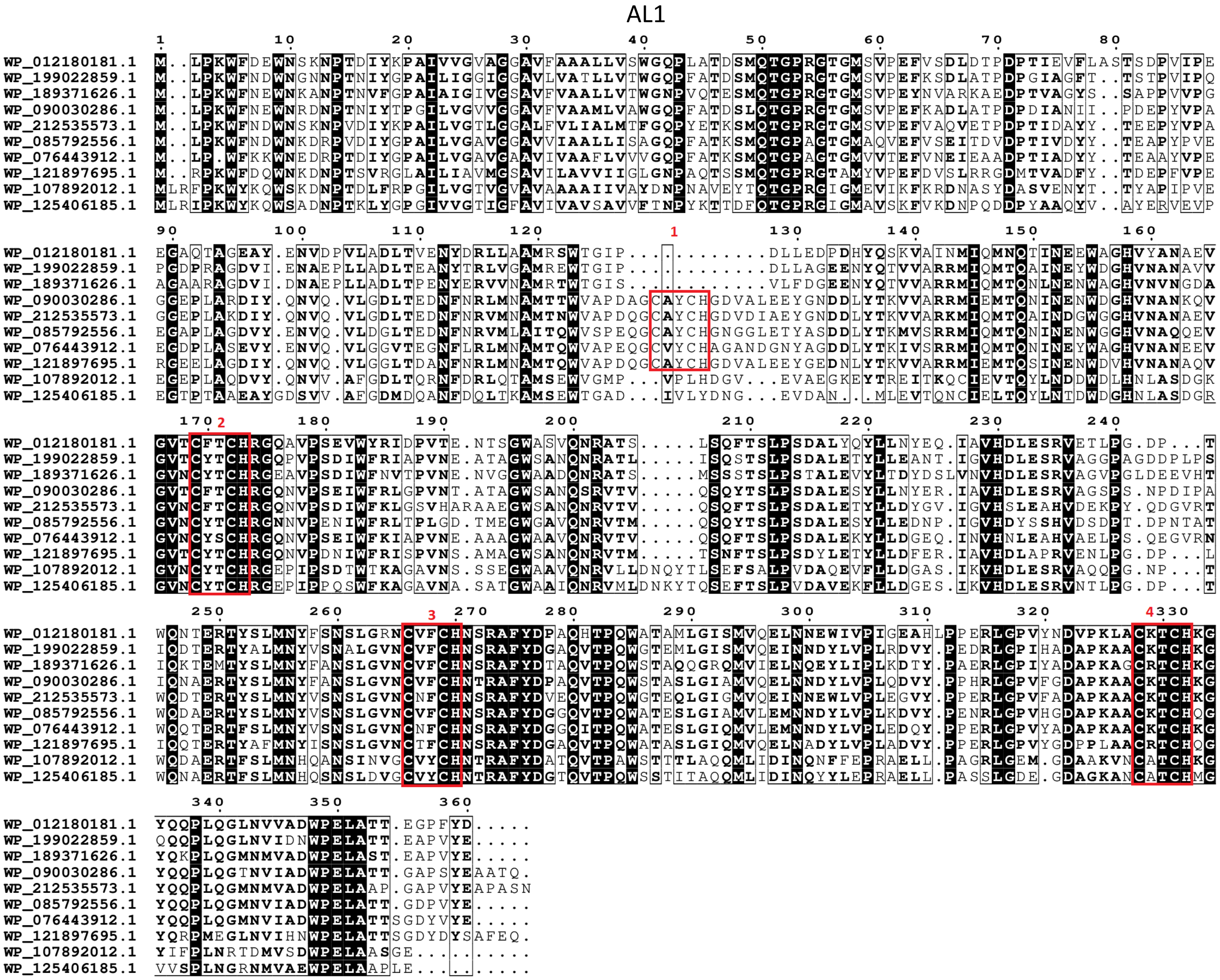


Figure S6. Maximum likelihood phylogenetic tree of cytochrome PufC family. Each presented tip is labeled with the RefSeq accession code, number of heme-binding motifs and if it contains a paralogue within this analysis and taxonomic class. Confidence values (expressed in %) of SH-aLRT/ultrafast bootstrap are presented near each node. Each heme-binding motif gain (green) / loss (blue) event was considered when confidence values are ≥ 80%/95%, respectively, otherwise only highlighted in gray. Confidence values below 70% are not shown. At the bottom of the tree, a subset of the aligned sequences is presented (those that are in bold in the phylogenetic tree) that are related to each event of heme-binding motif gain / loss (AL1).

#### ActA (5 hemes)

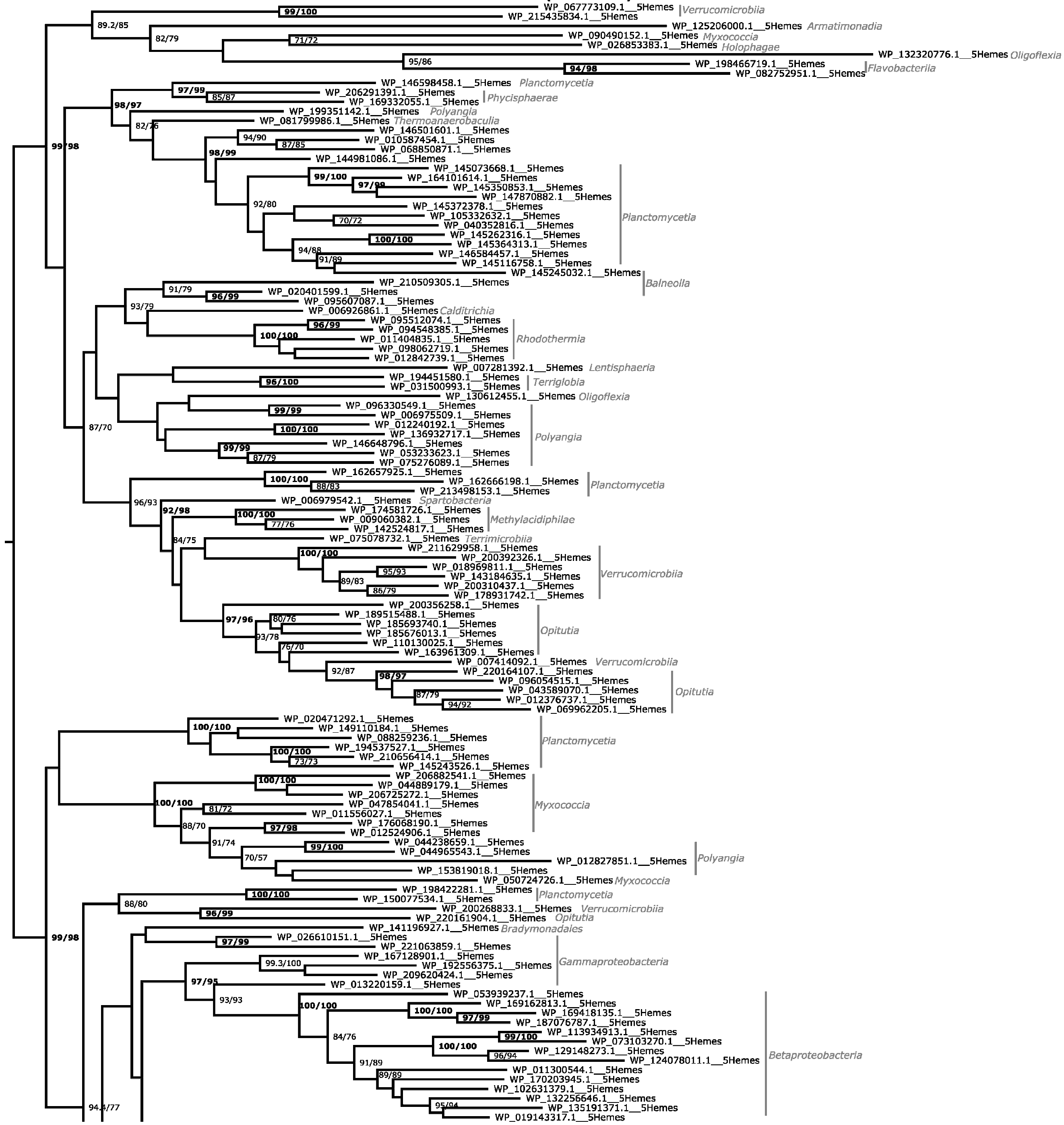

(Continues in next page)

(Continuation)

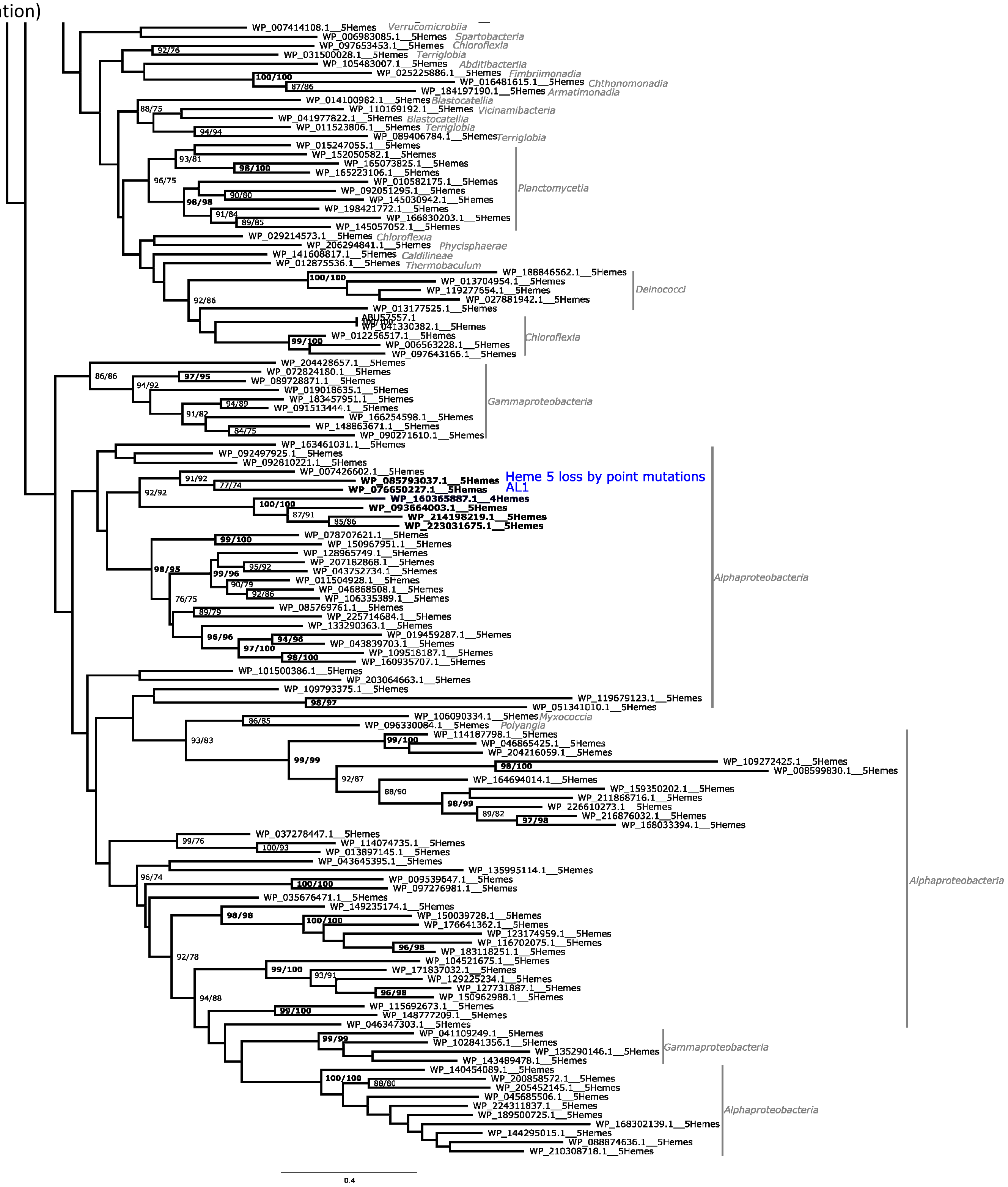

AL1

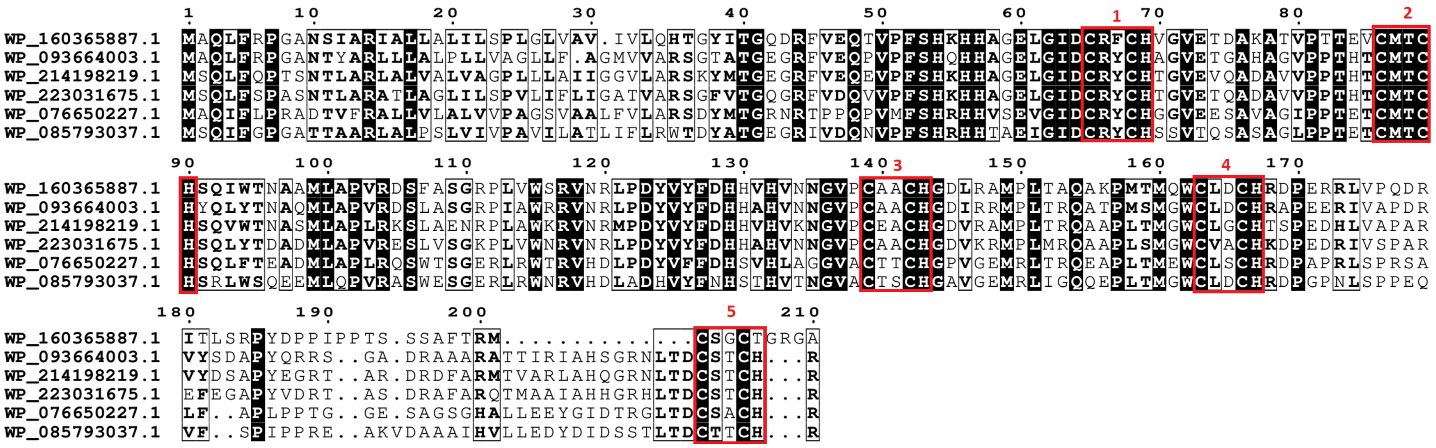


Figure S7. Maximum likelihood phylogenetic tree of cytochrome ActA (5 hemes) family. Each presented tip is labeled with the RefSeq accession code, number of heme-binding motifs and if it contains a paralogue within this analysis and taxonomic class. Confidence values (expressed in %) of SH-aLRT/ultrafast bootstrap are presented near each node. Each heme-binding motif gain (green) / loss (blue) event was considered when confidence values are ≥ 80%/95%, respectively, otherwise only highlighted in gray. Confidence values below 70% are not shown. At the bottom of the tree, a subset of the aligned sequences is presented (those that are in bold in the phylogenetic tree) that are related to each event of heme-binding motif gain / loss (AL1).

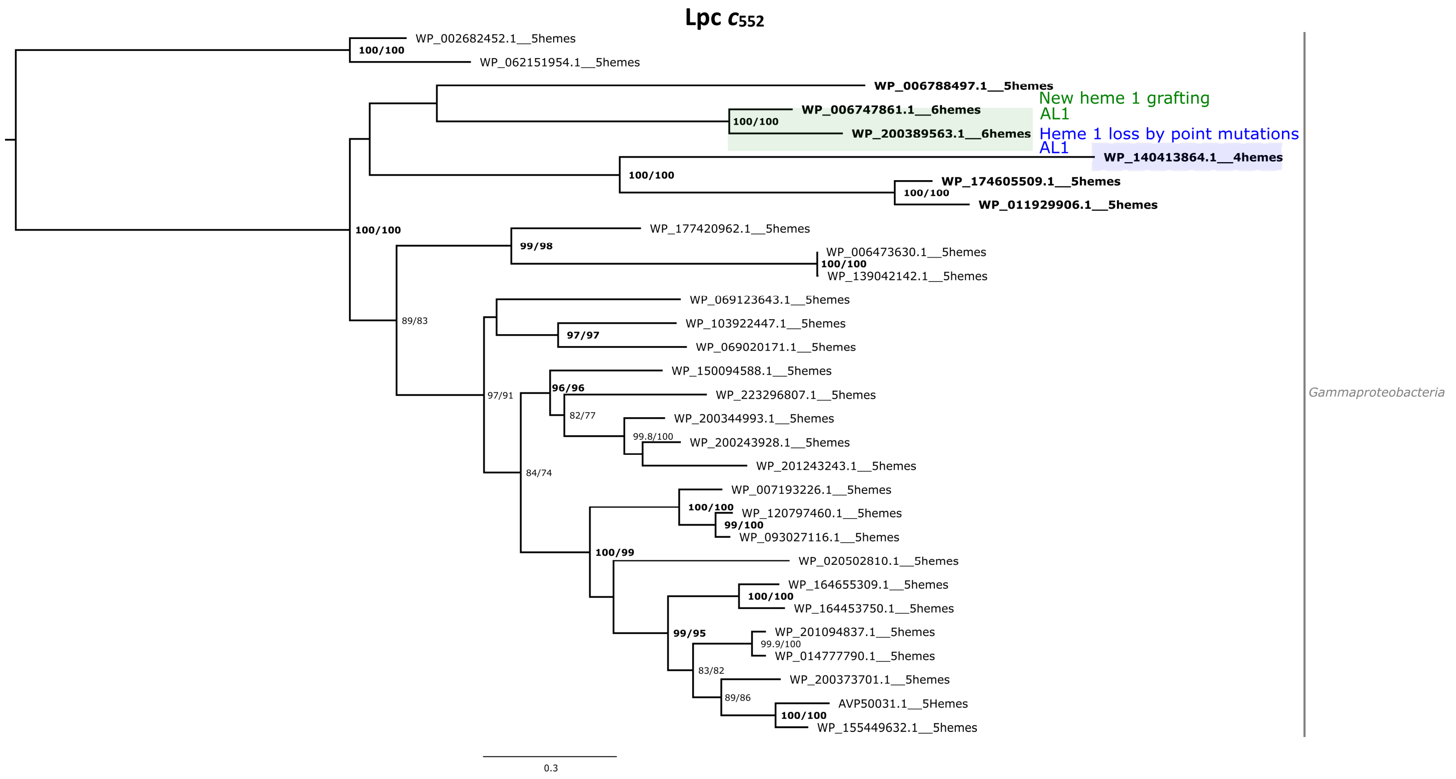


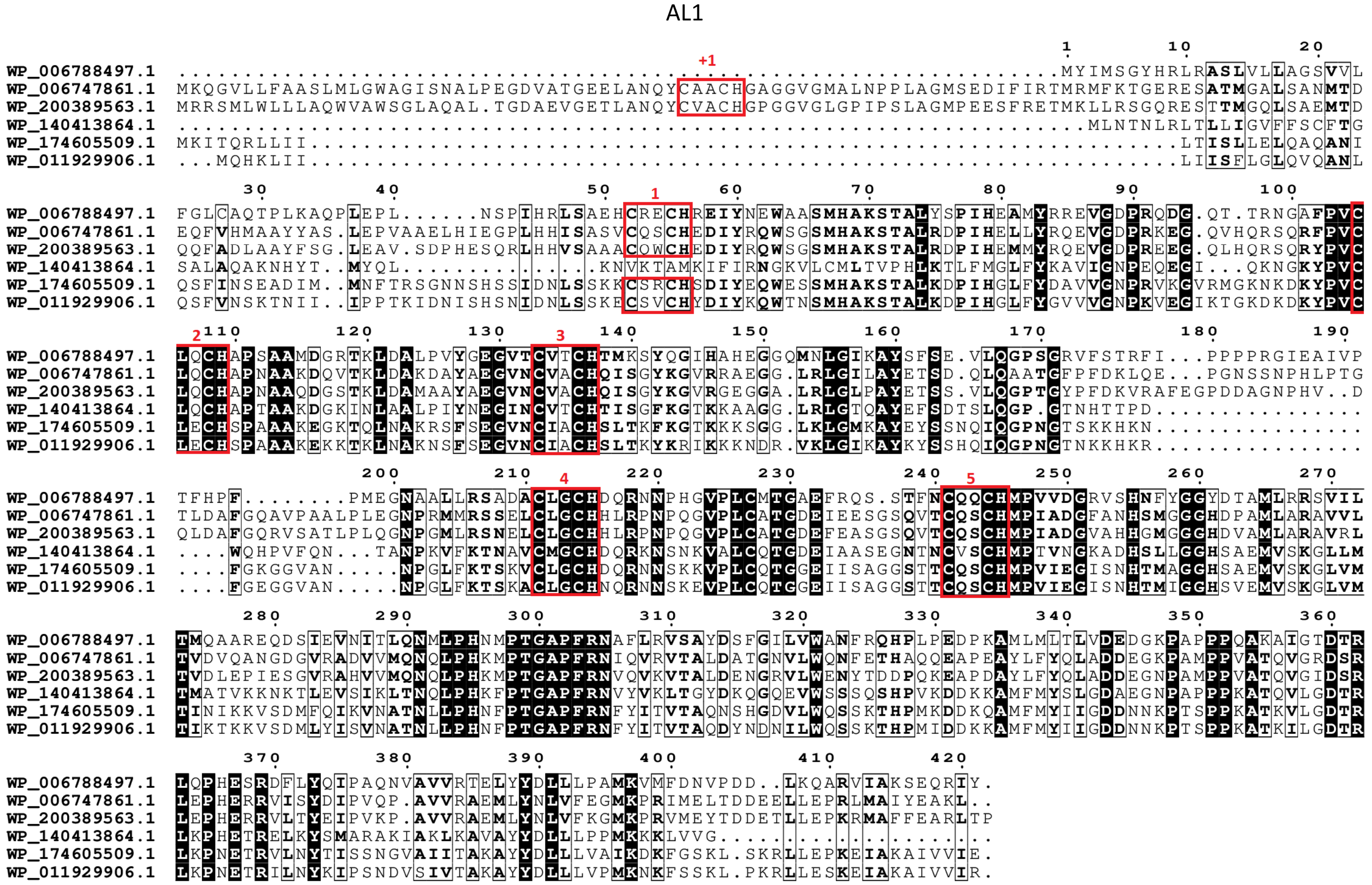


Figure S8. Maximum likelihood phylogenetic tree of cytochrome $c_{552}$ family. Each presented tip is labeled with the RefSeq accession code, number of heme-binding motifs and if it contains a paralogue within this analysis and taxonomic class. Confidence values (expressed in %) of SH-aLRT/ultrafast bootstrap are presented near each node. Each heme-binding motif gain (green) / loss (blue) event was considered when confidence values are ≥ 80%/95%, respectively, otherwise only highlighted in gray. Confidence values below 70% are not shown. At the bottom of the tree, a subset of the aligned sequences is presented (those that are in bold in the phylogenetic tree) that are related to each event of heme-binding motif gain / loss (AL1).

#### ActA (6 Hemes)

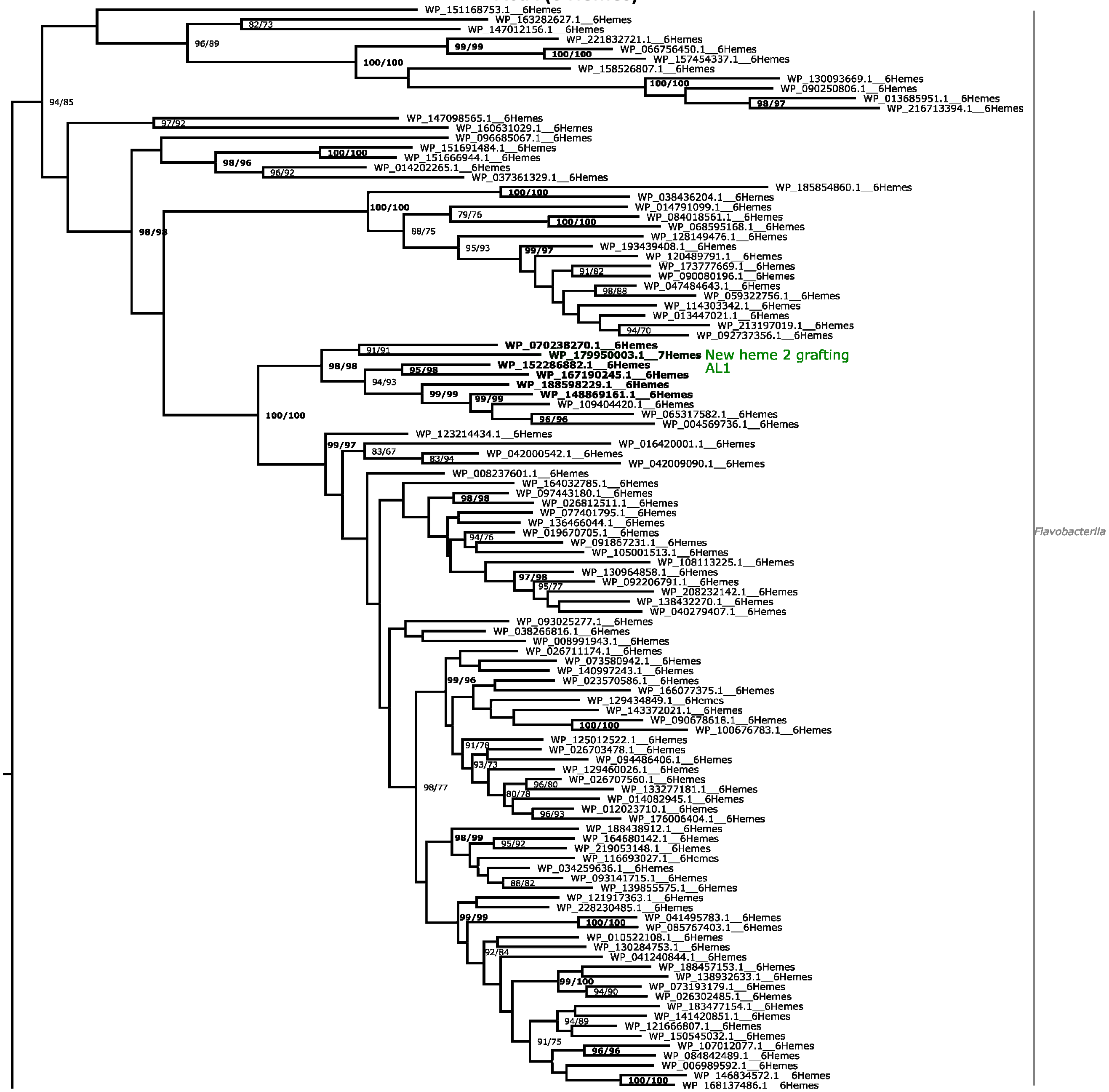


(Continues in next page)

(Continuation)

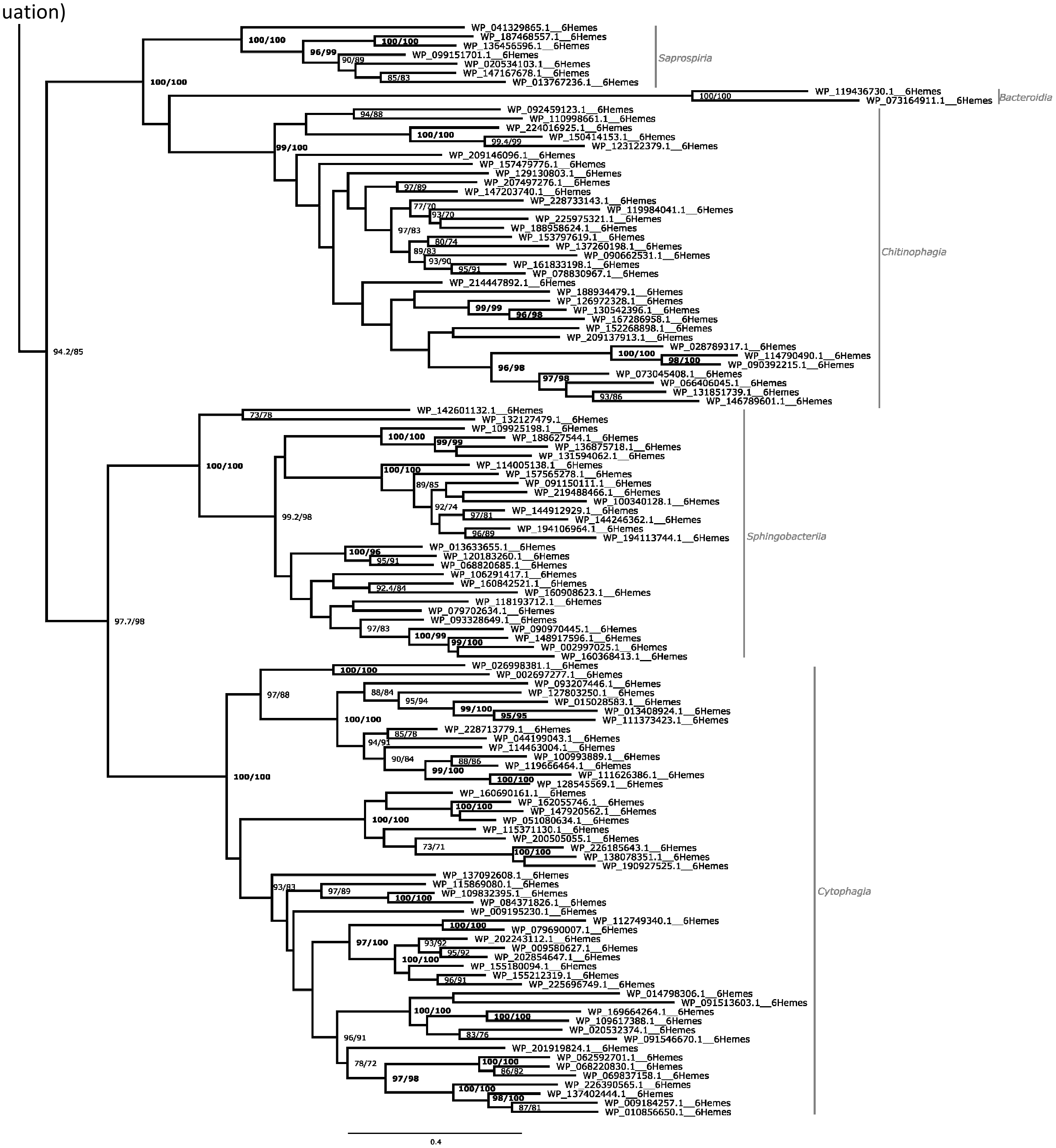

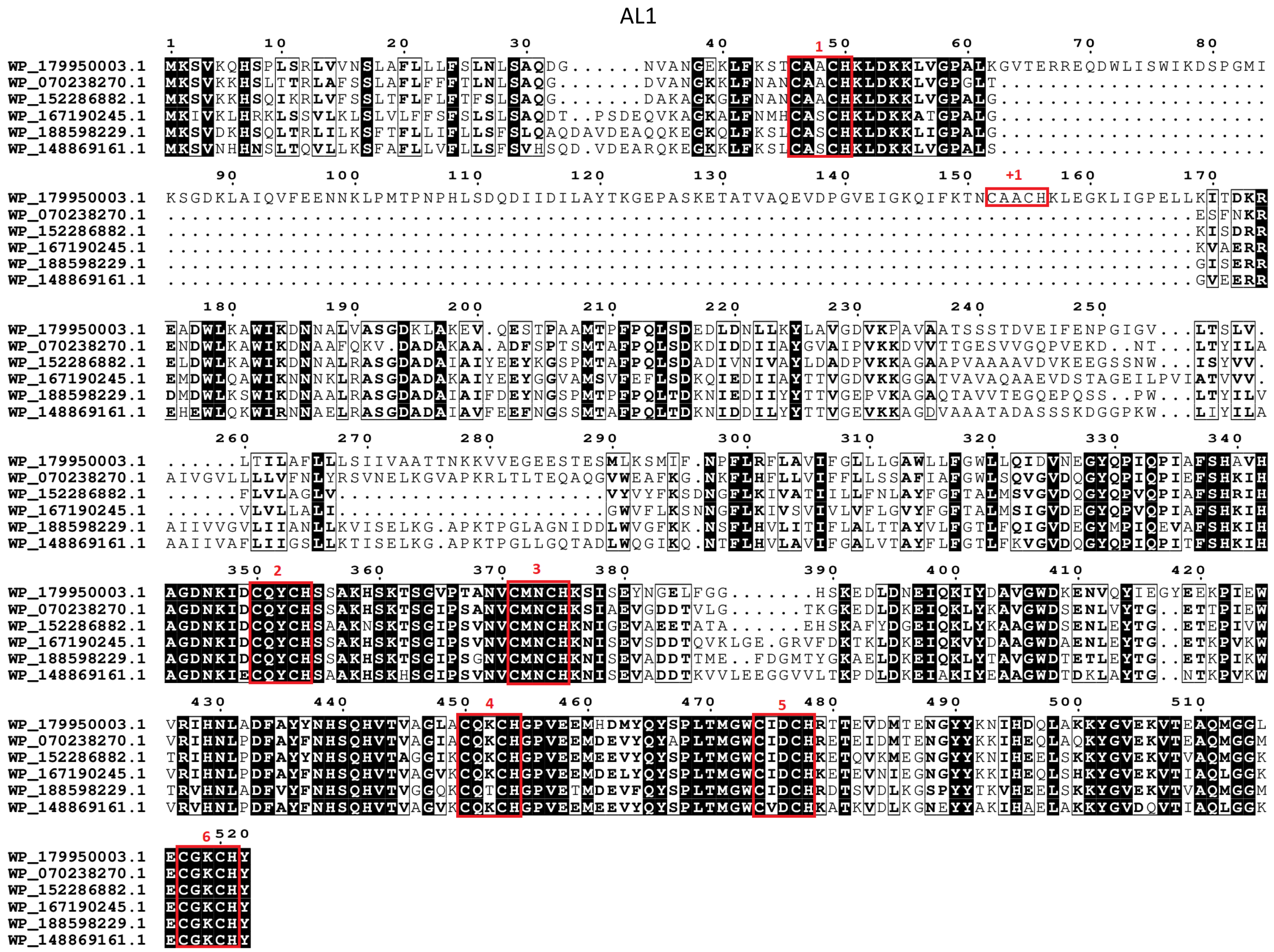


Figure S9. Maximum likelihood phylogenetic tree of ActA (6 hemes) family. Each presented tip is labeled with the RefSeq accession code, number of heme-binding motifs and if it contains a paralogue within this analysis and taxonomic class. Confidence values (expressed in %) of SH-aLRT/ultrafast bootstrap are presented near each node. Each heme-binding motif gain (green) / loss (blue) event was considered when confidence values are ≥ 80%/95%, respectively, otherwise only highlighted in gray. Confidence values below 70% are not shown At the bottom of the tree, a subset of the aligned sequences is presented (those that are in bold in the phylogenetic tree) that are related to each event of heme-binding motif gain / loss (AL1).

#### IhOCC

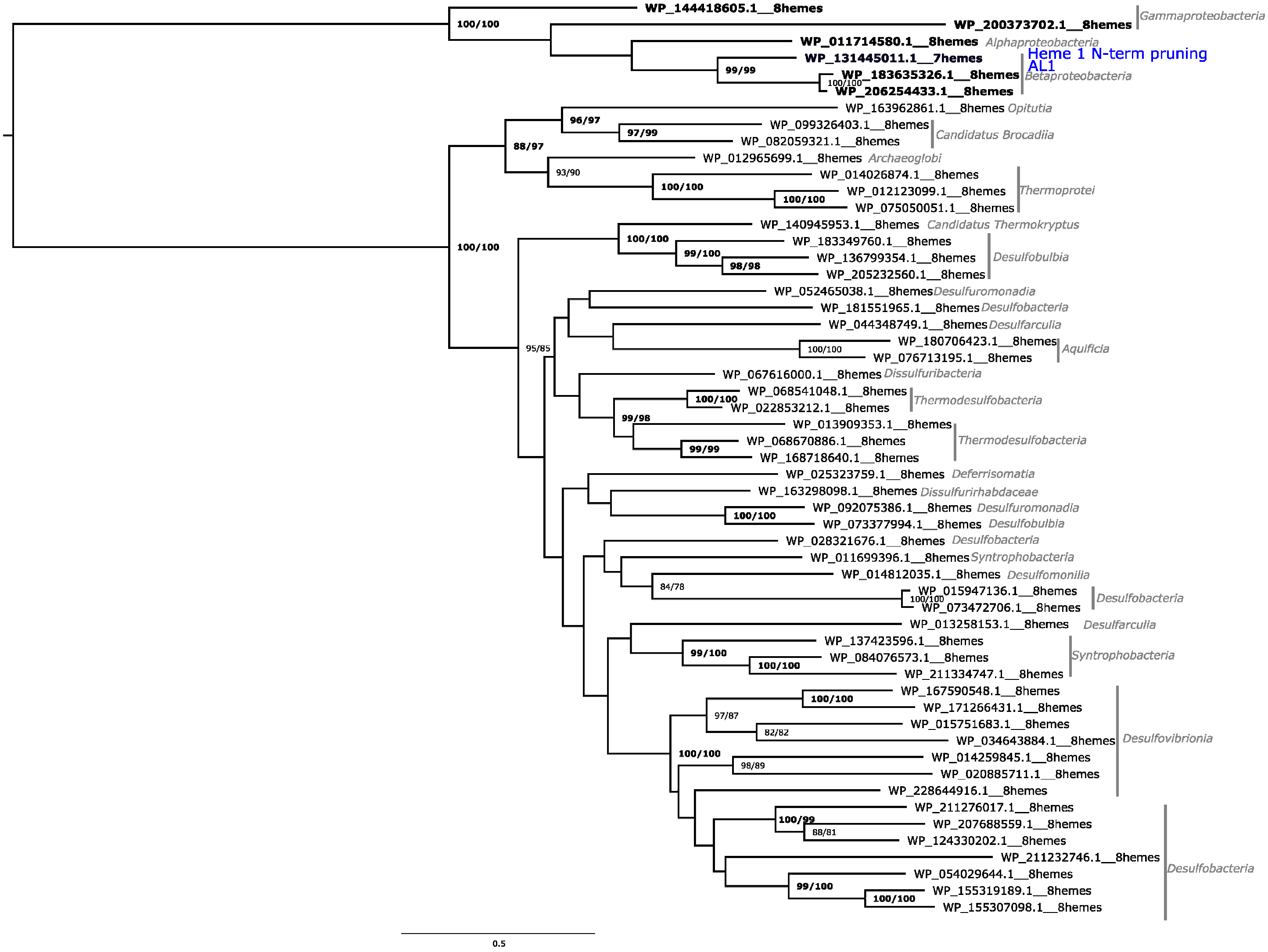
WP_144418605.1__8hemes
WP_200373702.1__8hemes
Gammaproteobacteria
WP_011714580.1__8hemes
Alphaproteobacteria
WP_131445011.1__7hemes
Heme 1 N-term pruning
AL1
WP_183635326.1__8hemes
WP_206254433.1__8hemes
Betaproteobacteria
WP_163962861.1__8hemes Opitutia
WP_099326403.1__8hemes
WP_082059321.1__8hemes
Candidatus Brocadiia
WP_012965699.1__8hemes Archaeoglobi
WP_014026874.1__8hemes
WP_012123099.1__8hemes
WP_075050051.1__8hemes
Thermoprotei
WP_140945953.1__8hemes Candidatus Thermokryptus
WP_183349760.1__8hemes
WP_136799354.1__8hemes
WP_205232560.1__8hemes
Desulfobulbia
WP_052465038.1__8hemes Desulfuromonadia
WP_181551965.1__8hemes Desulfobacteria
WP_044348749.1__8hemes Desulfarculia
WP_180706423.1__8hemes
WP_076713195.1__8hemes
Aquificia
WP_067616000.1__8hemes Dissulfuribacteria
WP_068541048.1__8hemes
WP_022853212.1__8hemes
Thermodesulfobacteria
WP_013909353.1__8hemes
WP_068670886.1__8hemes
WP_168718640.1__8hemes
Thermodesulfobacteria
WP_025323759.1__8hemes Deferrisomatia
WP_163298098.1__8hemes Dissulfurirhabdaceae
WP_092075386.1__8hemes Desulfuromonadia
WP_073377994.1__8hemes Desulfobulbia
WP_028321676.1__8hemes Desulfobacteria
WP_011699396.1__8hemes Syntrophobacteria
WP_014812035.1__8hemes Desulfomonilia
WP_015947136.1__8hemes
WP_073472706.1__8hemes
Desulfobacteria
WP_013258153.1__8hemes Desulfarculia
WP_137423596.1__8hemes
WP_084076573.1__8hemes
WP_211334747.1__8hemes
Syntrophobacteria
WP_167590548.1__8hemes
WP_171266431.1__8hemes
WP_015751683.1__8hemes
WP_034643884.1__8hemes
WP_014259845.1__8hemes
WP_020885711.1__8hemes
Desulfovibrionia
WP_228644916.1__8hemes
WP_211276017.1__8hemes
WP_207688559.1__8hemes
WP_124330202.1__8hemes
WP_211232746.1__8hemes
WP_054029644.1__8hemes
WP_155319189.1__8hemes
WP_155307098.1__8hemes
Desulfobacteria
0.5

## AL1

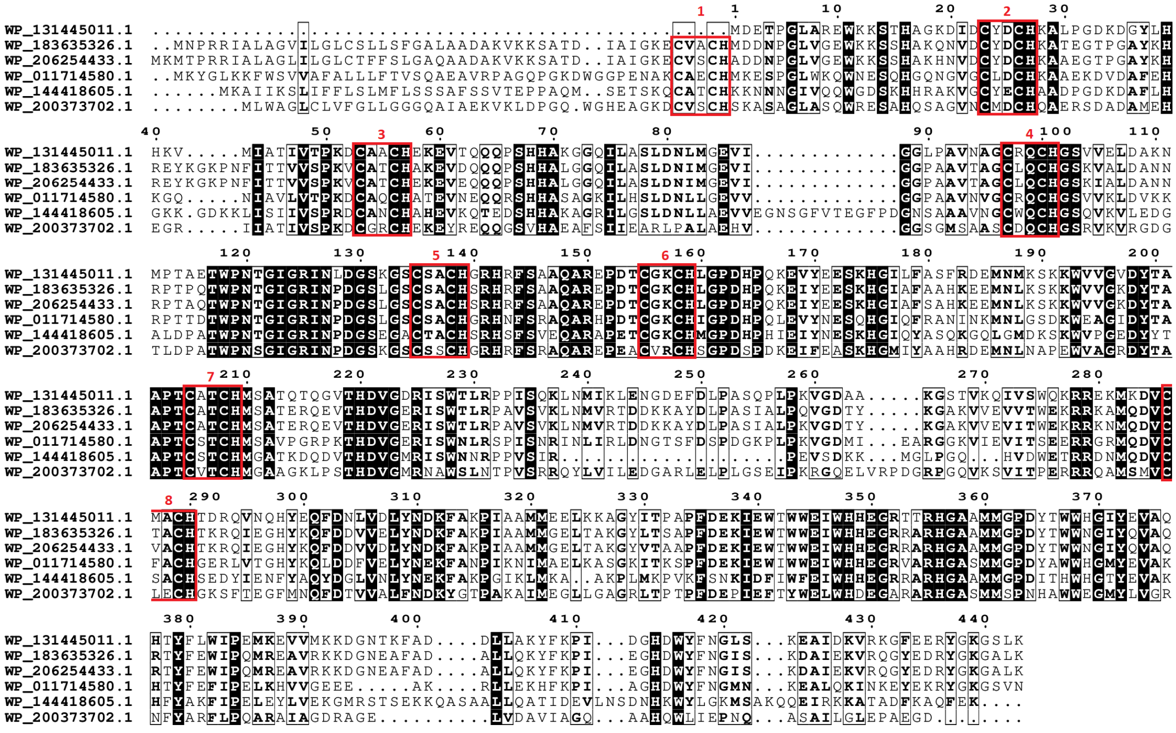
WP_131445011.1
WP_183635326.1
WP_206254433.1
WP_011714580.1
WP_144418605.1
WP_200373702.1

Figure S10. Maximum likelihood phylogenetic tree of IhOCC family. Each presented tip is labeled with the RefSeq accession code, number of heme-binding motifs and if it contains a paralogue within this analysis and taxonomic class. Confidence values (expressed in %) of SH-aLRT/ultrafast bootstrap are presented near each node. Each heme-binding motif gain (green) / loss (blue) event was considered when confidence values are ≥ 80%/95%, respectively, otherwise only highlighted in gray. Confidence values below 70% are not shown. At the bottom of the tree, a subset of the aligned sequences is presented (those that are in bold in the phylogenetic tree) that are related to each event of heme-binding motif gain / loss (AL1).

#### ONR

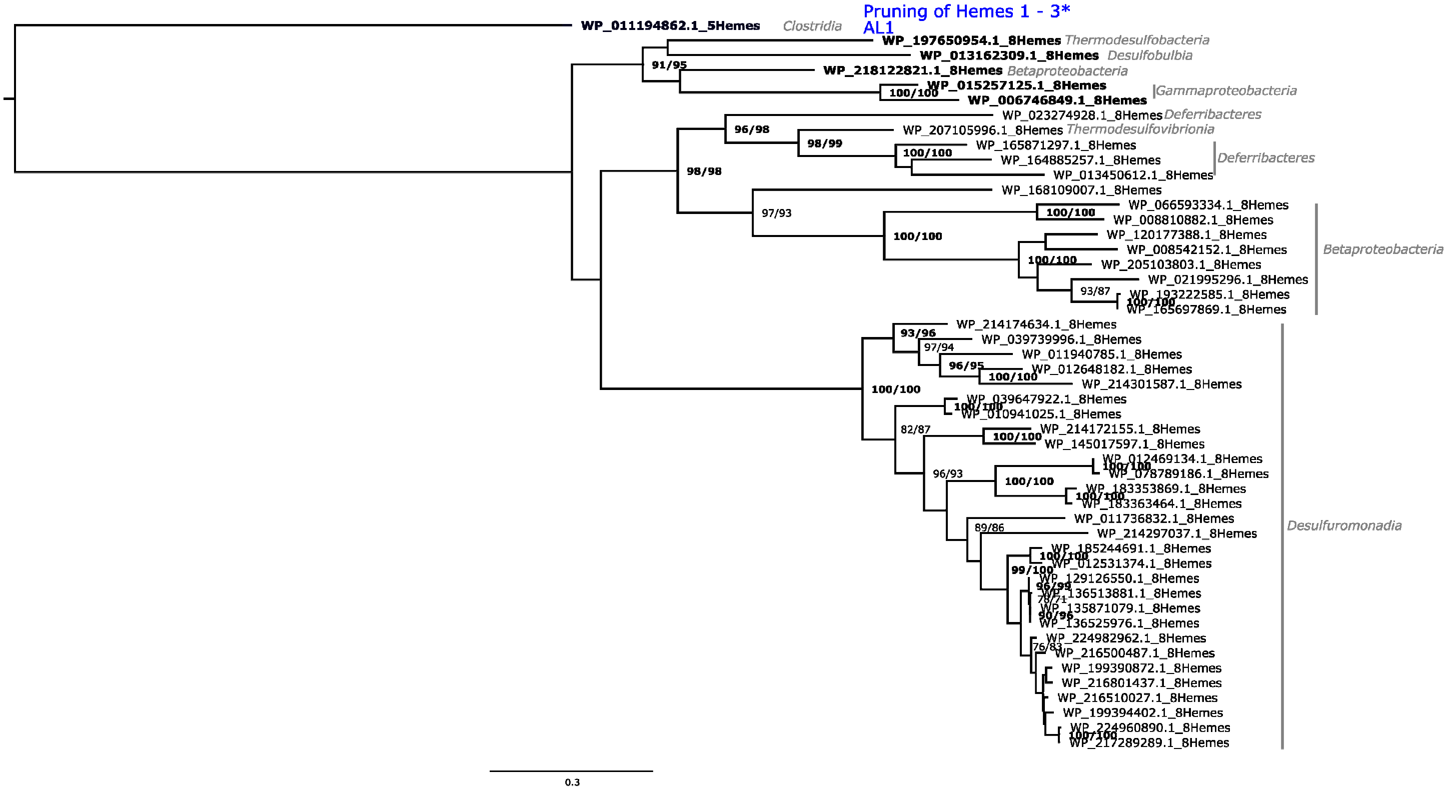

Pruning of Hemes 1 - 3*
AL1
WP_011194862.1_5Hemes
Clostridia
WP_197650954.1_8Hemes Thermodesulfobacteria
WP_013162309.1_8Hemes Desulfobulbia
WP_218122821.1_8Hemes Betaproteobacteria
WP_015257125.1_8Hemes
WP_006746849.1_8Hemes
Gammaproteobacteria
WP_023274928.1_8Hemes Deferribacteres
WP_207105996.1_8Hemes Thermodesulfovibrionia
WP_165871297.1_8Hemes
WP_164885257.1_8Hemes
WP_013450612.1_8Hemes
Deferribacteres
WP_168109007.1_8Hemes
WP_066593334.1_8Hemes
WP_008810882.1_8Hemes
WP_120177388.1_8Hemes
WP_008542152.1_8Hemes
WP_205103803.1_8Hemes
WP_021995296.1_8Hemes
WP_193222585.1_8Hemes
WP_165697869.1_8Hemes
Betaproteobacteria
WP_214174634.1_8Hemes
WP_039739996.1_8Hemes
WP_011940785.1_8Hemes
WP_012648182.1_8Hemes
WP_214301587.1_8Hemes
WP_039647922.1_8Hemes
WP_010941025.1_8Hemes
WP_214172155.1_8Hemes
WP_145017597.1_8Hemes
WP_012469134.1_8Hemes
WP_078789186.1_8Hemes
WP_183353869.1_8Hemes
WP_183363464.1_8Hemes
WP_011736832.1_8Hemes
WP_214297037.1_8Hemes
WP_185244691.1_8Hemes
WP_012531374.1_8Hemes
WP_129126550.1_8Hemes
WP_136513881.1_8Hemes
WP_135871079.1_8Hemes
WP_136525976.1_8Hemes
WP_224982962.1_8Hemes
WP_216500487.1_8Hemes
WP_199390872.1_8Hemes
WP_216801437.1_8Hemes
WP_216510027.1_8Hemes
WP_199394402.1_8Hemes
WP_224960890.1_8Hemes
WP_217289289.1_8Hemes
Desulfuromonadia
91/95
100/100
96/98
98/99
98/98
97/93
93/96
97/94
96/95
82/87
96/93
89/86
99/100
93/87
76/83
0.3


## AL1

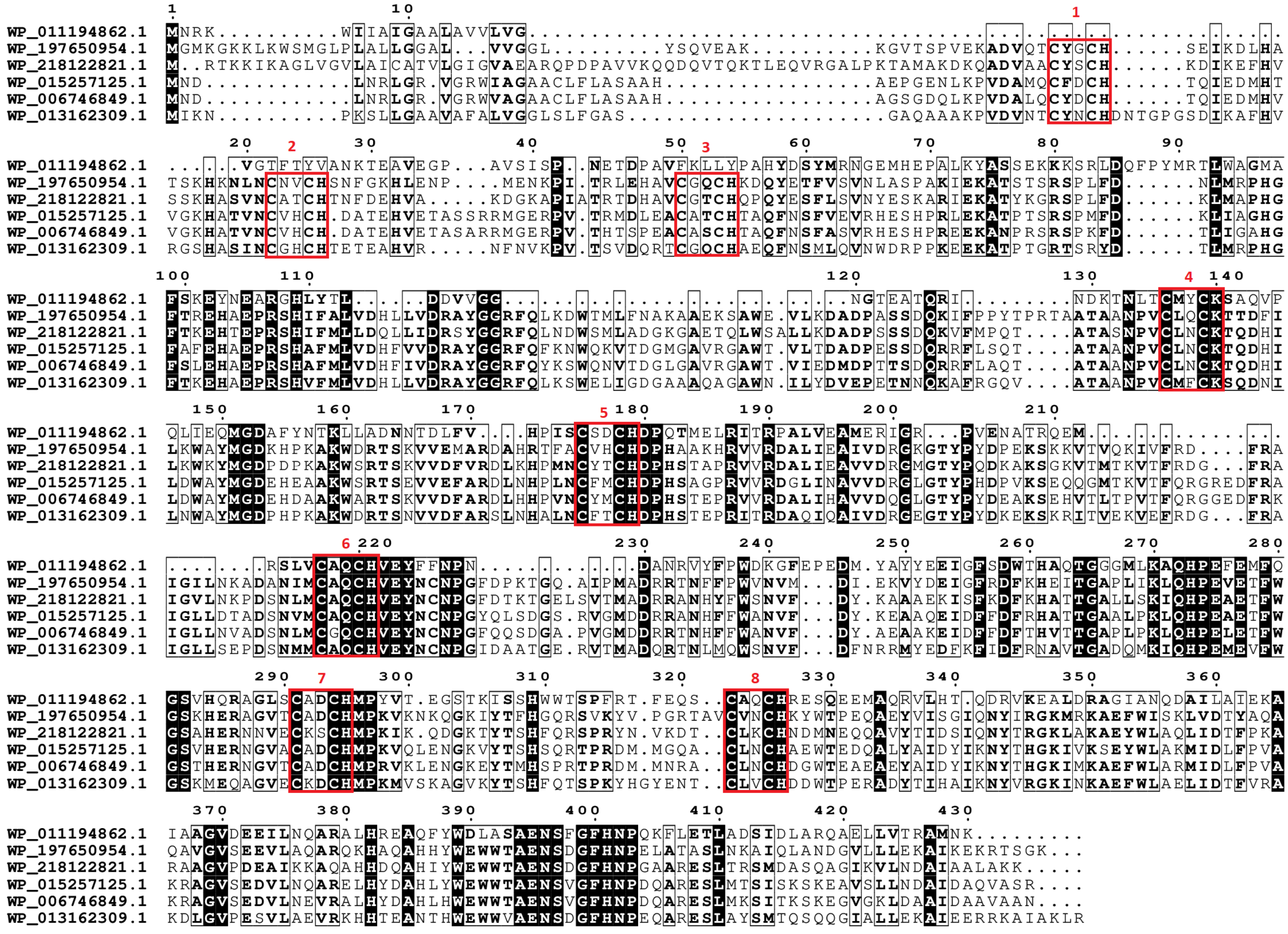

WP_011194862.1
WP_197650954.1
WP_218122821.1
WP_015257125.1
WP_006746849.1
WP_013162309.1
1
2
3
4
5
6
7
8

Figure S11. Maximum likelihood phylogenetic tree of ONR family. Each presented tip is labeled with the RefSeq accession code, number of heme-binding motifs and if it contains a paralogue within this analysis and taxonomic class. Confidence values (expressed in %) of SH-aLRT/ultrafast bootstrap are presented near each node. Each heme-binding motif gain (green) / loss (blue) event was considered when confidence values are ≥ 80%/95%, respectively, otherwise only highlighted in gray. Confidence values below 70% are not shown. At the bottom of the tree, a subset of the aligned sequences is presented (those that are in bold in the phylogenetic tree) that are related to each event of heme-binding motif gain / loss (AL1). Prunning event was considered by the previous work (Soares et. al 2022).

### OTR

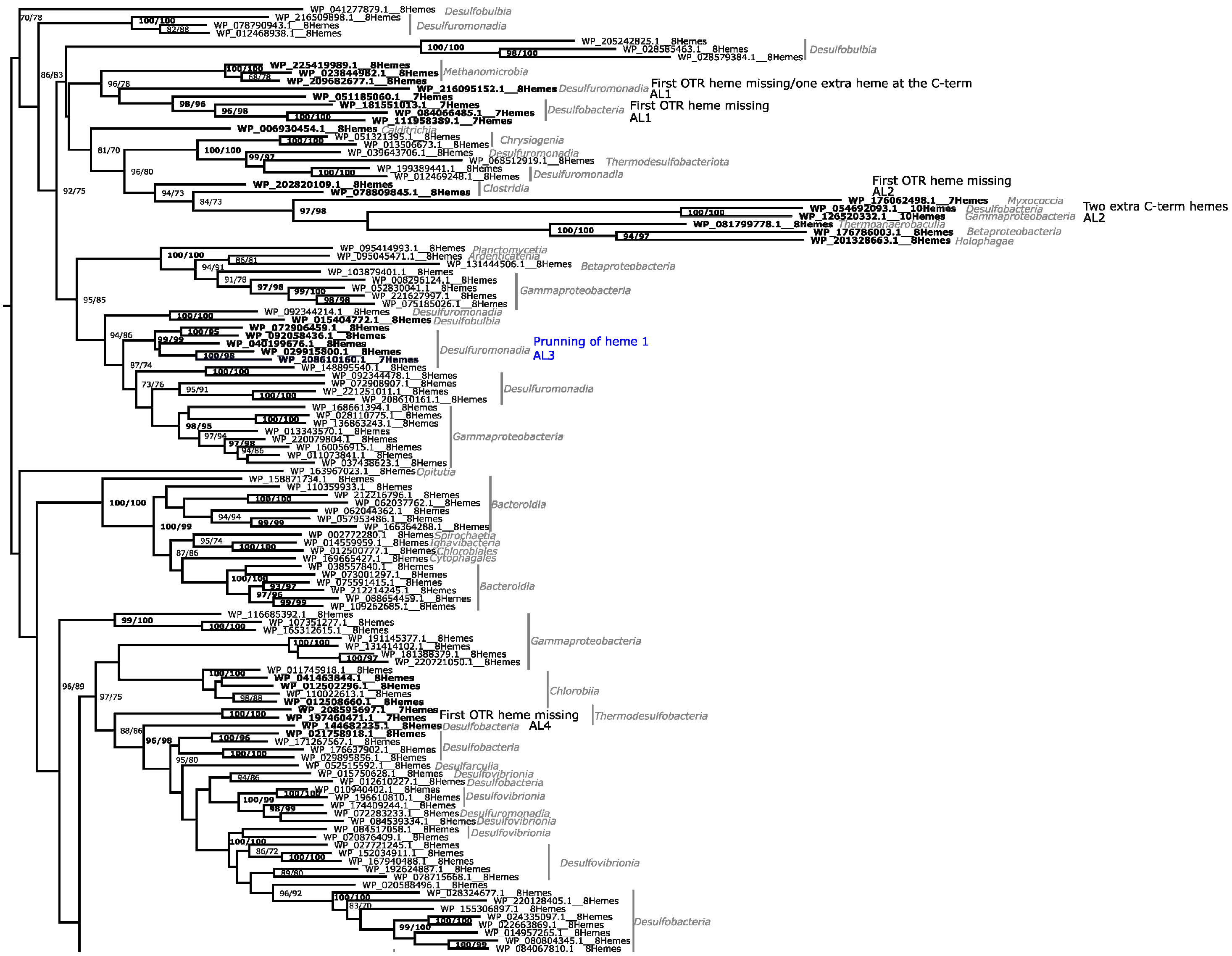


(Continues in next page)

(Continued)

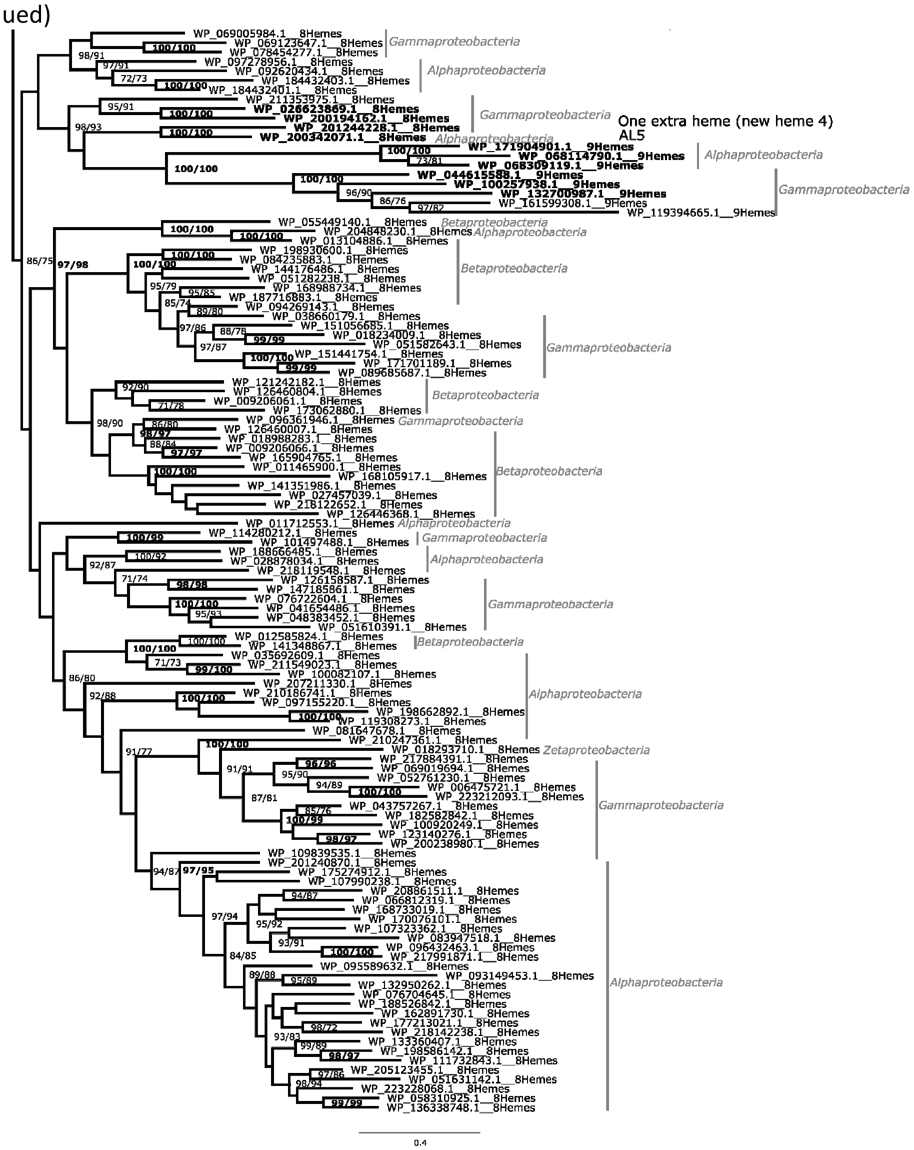
Gammaproteobacteria
Alphaproteobacteria
Gammaproteobacteria
One extra heme (new heme 4)
AL5
Alphaproteobacteria
Alphaproteobacteria
Gammaproteobacteria
Betaproteobacteria
Alphaproteobacteria
Betaproteobacteria
Gammaproteobacteria
Betaproteobacteria
Gammaproteobacteria
Betaproteobacteria
Alphaproteobacteria
Gammaproteobacteria
Alphaproteobacteria
Gammaproteobacteria
Betaproteobacteria
Alphaproteobacteria
Zetaproteobacteria
Gammaproteobacteria
Alphaproteobacteria
0.4

AL1

```
                                                                  1
WP_051185060.1  ..........................................................................................
WP_181551013.1  ..........................................................................................
WP_084066485.1  ..........................................................................................
WP_111958389.1  ..........................................................................................
WP_209682677.1  MKICLATGIFLILLIVAVFPASSATMTHSFLEGPYDSGPEVTEECIECHVNQANDLLNSTHWLWTSCAACGCEE.EHLYDLGKRTVINNF
WP_023844982.1  .........MLACVIVLSFSSSASYLNHSFLTGPYENGPDVTADCISCHSQQAADMLNSTHWLWTTCSEC.CGD.EAYGGLGKRTVINNF
WP_225419989.1  .........................MNHSFLEGPYDSGPEVTEECIGCHEPQAKDMLNSTHWLWTSCGDCECGEIEAYKDMGKRTVINNF
WP_006930454.1  ......MRYILLIFFLLIVGHLFAEDHSSYIKGPFKNPQEVTKQCLTCHEESAQQIMKTSHWTWSTPPVKVPGH.EGLHRIGKKDIFNNY
WP_216095152.1  ..........................................................................................

                1        10 2      20        30   3    40        50        60        70    4   80
WP_051185060.1  .MSPIANWCSCTSCHAGYGWKENTFDFKDPSRVDCLVCHDRTGSYEKAPAGCGDEKKGLDLVKIARGVGRPTRANC.GSCHFCGGGGDAI
WP_181551013.1  ...MESNWCRCTKCHAGYGKEDDTFDFMDDAAVDCLVCHDQTGKYAGSKGECGYTDGSVDLLKVAQSVAKPTRSNC.GSCHWYGGGGNNV
WP_084066485.1  .MSIESNWCSCTKCHAGYGWKDASFDFSDMEKMDCLICHDTTGTYSKKKNSCGMPNKDLDLTDIAKQVGLPGRGNC.GECHWYGGGGDNA
WP_111958389.1  .MSIESNWCGCTKCHAGYGWKDENFDFTDMANVDCLVCHDTTGRYSKKKNKCGYPDKGIDLAQIAGSVGPPGRGNC.GACHWYGGGGDNF
WP_209682677.1  CVAVASNEPRCTSCHAGYGWEDDTFDFTNASNLDCLVCHETTGTYTKIPTGAGAVDPSVDLVAVAQSVGSPSRETCGGNCHFFGGGGDNV
WP_023844982.1  CVAIASNEPRCTSCHTGYGWEDDTFDFTNESNIDCVVCHETTGTYVKIPTGAGAVDPSVDLVAVAQSVGSPSRDTCGGSCHFFGGGGDNV
WP_225419989.1  CVAIASNEPRCTSCHAGYGWEDDTFDFNNASNIDCLVCHDNTGTYKKIPTGAGAVDTSVDLAEVAQSVGSPTRDTCGGNCHFYGGGGDNV
WP_006930454.1  CIHVQSNWPRCTSCHTGYGWKDDSFDFDKEENVDCLICHADPKIYKKNPAGAGFPKKDVDLLKAARSVRLPDRENC.GSCHFYGGGGENV
WP_216095152.1  .MSILSNESRCTSCHAGYGWTDASFDFADLSRIDCLVCHDRSGRYKKEPTNAGWPVKDLDLKPIAEQVGHSSRASC.GSCHFNGGGGDAI

                90        100       110     5 120       130       140  6      150       160       170 7
WP_051185060.1  KHGNLSEALIDPPRELDVHMGGLDFACQDCHTTNGHRIAGASMTTCV.SEGRVSCTDCHDERPHATDRHAVQKTLNDHCDAIACETCHIP
WP_181551013.1  KHGDMDAALADPSREHDVHMGGMDFQCQDCHTTENHKIAGRSTTSSV.SEGPVACTDCHDSRPHDKSS.AMLSKLNDHTDAVSCEACHIP
WP_084066485.1  KHGDLGSDLLTPDASLDVHMGKLDFTCQKCHVTDAHKIAGSSTTSSV.SEGRVSCTDCHDENPHDSSF.PLLRKLNEHGMAMACQTCHIP
WP_111958389.1  KHGDLSSEFLNPSRSFDVHMGGEDFSCQECHATDQHRIAGSSTTSAV.SEGSVSCLDCHDESPHDDAS.PLLKQLNGHCQSIACQTCHIP
WP_209682677.1  KHGDMSSALLEPSADLDVHMGGLDFSCQNCHETTAHNVAGRFAGTPG.SECRVECSDCHTETPHAG...EYKERLDAHSDSIACQTCHIP
WP_023844982.1  KHGDMSSALLDPSRELDVHMGGMDFDCQNCHKTTAHNIAGRFAGVPG.SECRVECTDCHGDTPHQG...VFKDRLDGHSDSIACQTCHIP
WP_225419989.1  KHGDMSSALSEPSPELDVHMGLLDFKCQNCHETSDHNVAGRFAGLPD.SECRVQCSDCHGQTPHAG...EYKERLDGHVDAIACQTCHIP
WP_006930454.1  KHGDLEQILANPTRDQDVHMGS.GMVCQDCHVHENHDFKGRGSSVSVLDENHLYCADCHGLDAHEN......ERLNLHVRSIACQTCHIP
WP_216095152.1  KHADMGNNLLDPDPRCDVHMGDLDFGCVDCHRTYQHRIAGRSSSVAP.AEGVVRCEDCHSAAPHYRNG.LLAAHLNRHSASLACNVCHSP

                180       190       200       210       220       230       240       250       260
WP_051185060.1  TFAKAKPTLLFWNWALAGRDTKELENSPYC....ITTQHRKKGLFIKKANVAPAYAWYNGRHERYRMGERVDVTGT.TYLNPPAGGIQDP
WP_181551013.1  EYAKDAPTTTLWDWSQSGREDKVLENNGKK....KRILSKKKGLLIREKNLTPEYEWYNGTHKRYLKGDPINREAG.TQINPPQGDIHDP
WP_084066485.1  QYAVNKFTLTYRDGSRISDKTEVVSQS.PD....KRELKTPMGQKILEKNLKPSYAWYNGKHHRYMKGDPVDPKGI.TPLNAPDGTIKDP
WP_111958389.1  TFARKKFTLTSRDFSKGAKENKILEQS.EN....KIVVQGKLGITVKEKALRPTYAWYNGKHHRYLKGDTIKTSGV.TALNSPVGSIKDP
WP_209682677.1  TYAREIPTKMFWDWSKAGQDIDSIPVDEYG....KATYDKKKGEFVWAMDVRPDYAWFNGSFDLYRLGDELDPDEV.DVIKEPLGSREDS
WP_023844982.1  EFAREVPTKMYWDWSQAGQDIDPIPTDEYG....KATYDKKKGSFVWGMNVTPTYLWFNGSFDKYYLGDEVDPDGV.TVLTLPLGDRYDR
WP_225419989.1  QYAREVPTKMYWDWSQAGQDIDPVPTDEYG....KATYHKKKGSFVWEKNVTPSYAWYNGSSTLYKLGDTINTDGV.TVMNAPLGSRDDA
WP_006930454.1  TFAKGMATKIWWDWSKAGMDIEPKEE..YG....RETFNKKKGEFHWAKNVRPEYAWYNGTMDRYLKGDKLDPANI.LYLNRPRGSVYDN
WP_216095152.1  VYAKCTPTKNWWDWSKAGDTGRQPQMTRLGDSDPLPDYHVQKGEFAWQRAATPDYVWFDGTMERVLVGDAVPAGTTPVQLTAPLGQRHDP

                    270       280       290            300       310       320       330       340 8
WP_051185060.1  EAKITPFKIHTGVQPADATFQYLLVPELW.....G..GYWEHFDWERAIRSGMKEAELPYSGKSTFVTTVMHWRLNHGVAPKEQALGCMD
WP_181551013.1  EARITPYKIMKGVEPADPVNGHLIVPKLF.....GDGGYFETYDWVDAAEKGMAAAGLDFSGEVVFVETIMHWRLNHQVAPKEEAFSCLD
WP_084066485.1  SAKITPYKIIKTRQAIDARHGYLIVPHLA.....GKDGLISTHDWQVTAEKGMKAAGLKFSGEIAFADTMMYWRLNHGVVPKEKALSCLD
WP_111958389.1  LARITPYKLVKSRQPMDSENRILIVPHLT.....GKDGFMATGDWKLAAEKGMKAAKLPFSGKVDFVETVMYWRLNHGVVPKEESLSCLD
WP_209682677.1  DSKIYPFKIHVAKQISDAQYNYLIIPDLF....GGEKSYWATYDWNKAAEAGMDYVGMPYSGEYEFVETALYESINHEVAPASQALQCDD
WP_023844982.1  DSKIYPFKIHIAKQISDAQYNYLIIPDLF....GGENSYWASYDWDKAARAGMEYVDVPYSGQYEFVETSLYESINHEVAPKEDALQCKE
WP_225419989.1  DSQIYPFKIHRAKQISDAEYNYLIVPDLF....GGENSYWATYDWDKAASGMEYVDMPYSGEYEFVETSLYESINHEVPPAEHSLDCSD
WP_006930454.1  KAKIYPFKVMRGKQIYDAEYNYLIIPNLW.....G..GYWKHFNWDKAAKDGMAAAGLPYSGHYAFVETKMYWRINHEVVPKEKALQCVD
WP_216095152.1  QARITPFKVMKGVQAFDSEHGTLLIPHLFPRGAADRTAYWKNFDWHQAFSDGMAVAGLPYSGRWHWRETWTWWRVEHEVMPARLALTCVS

                     +1                                          350       360       370       380
WP_051185060.1  CHGA..............................................KALMDFEALGYPGDPAVAGGRLVSGPEGPKERKRP.....
WP_181551013.1  CHGC..............................................DKVMDFRALGYEGDPADVGGRSAGRQ..............
WP_084066485.1  CHGP..............................................EGVMDFKVLGYKGDPMATGWRFAPDK..............
WP_111958389.1  CHSP..............................................SGVMDFKALGYKQDPAGGGARFEK................
WP_209682677.1  CHLE..............................................GG.MDFVTLGYSGDPMIVGDQRPASSTTED...PADVPE
WP_023844982.1  CHLD..............................................DGVMDFAALGYEGDPMIVGERFMSEEEIEEI.ESMDTETTE
WP_225419989.1  CHLE..............................................TSMMDFETLGYEGDPMLVGERF..SEEVEDVRQSVEQEAPE
WP_006930454.1  CHGK..............................................NSILDWEKLGYEGDPMFMGDRKKEELIREYDLERK......
WP_216095152.1  CHDSLRGEQTCDRCHQDSRHVNFRELAHKPTDFSFLAGKRDDLDQLRQNGNYLDFTALGYAGDPILHGGRFSRLPLGRRPADSPSPHPKE

WP_051185060.1  ................................
WP_181551013.1  ................................
WP_084066485.1  ................................
WP_111958389.1  ................................
WP_209682677.1  AEATPTEATPGFSILLTSGVMLLLTLLLRRKD
WP_023844982.1  SGTESSESSPGFEFFVGVFGLIAAVLILKCK.
WP_225419989.1  AGEESSEGVPGFGIFLGIGMLLVVSILWGRR.
WP_006930454.1  ................................
WP_216095152.1  EP..............................
```

AL2

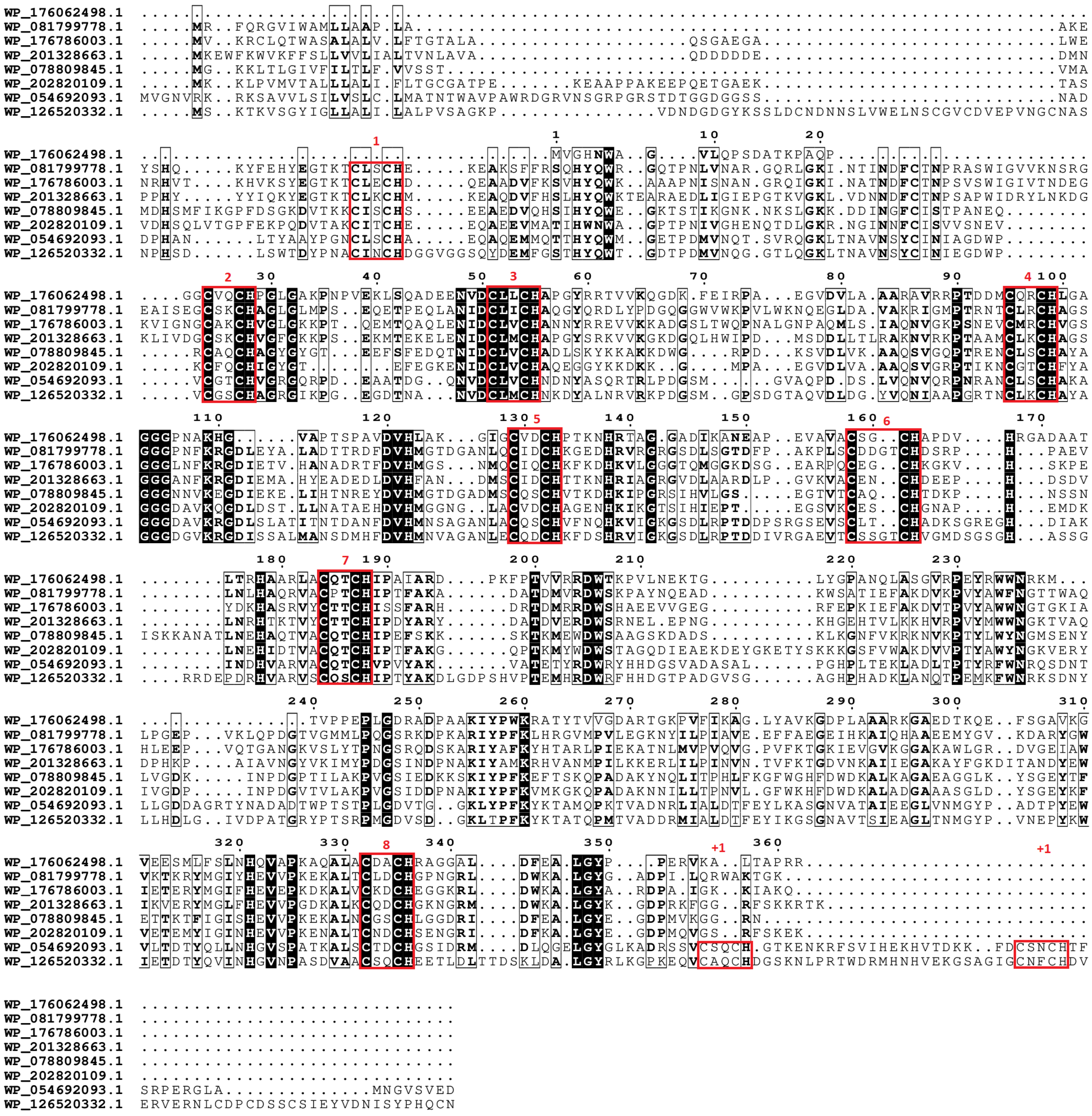
WP_176062498.1
WP_081799778.1
WP_176786003.1
WP_201328663.1
WP_078809845.1
WP_202820109.1
WP_054692093.1
WP_126520332.1
1
2
3
4
5
6
7
8
+1
+1

AL3

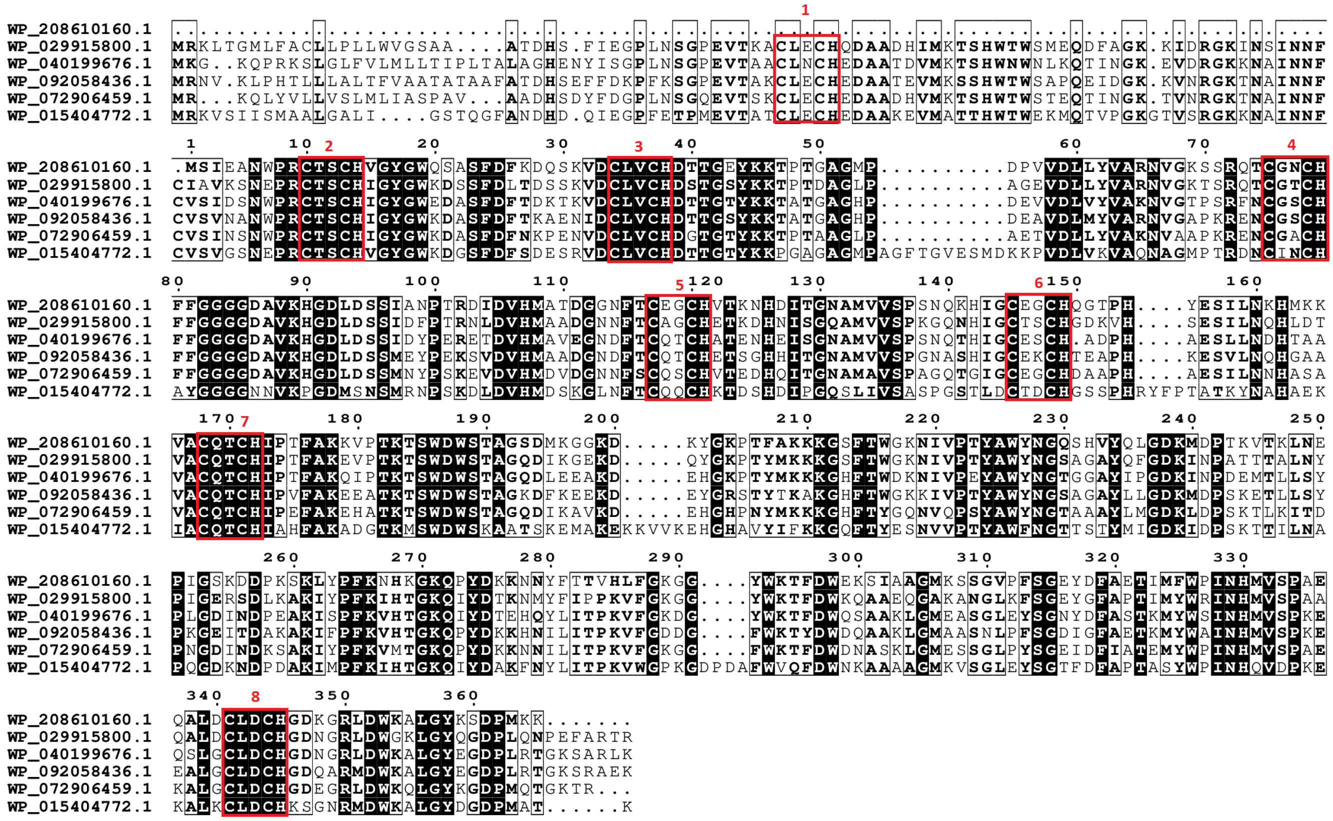

1
WP_208610160.1
WP_029915800.1 MRKLTGMLFACLLPLLWVGSAA....ATDHS.FIEGPLNSGPEVTKACLECHQDAADHIMKTSHWTWSMEQDFAGK.KIDRGKINSINNF
WP_040199676.1 MKG..KQPRKSLGLFVLMLLTIPLTALAGHENYISGPLNSGPEVTAACLNCHEDAATDVMKTSHWNWNLKQTINGK.EVDRGKKNAINNF
WP_092058436.1 MRNV.KLPHTLLLALTFVAATATAAFATDHSEFFDKPFKSGPEVTAKCLECHEDAATEVMKSSHWTWSAPQEIDGK.KVNRGKTNAINNF
WP_072906459.1 MR...KQLYVLLVSLMLIASPAV...AADHSDYFDGPLNSGQEVTSKCLECHEDAADHVMKTSHWTWSTEQTINGK.TVNRGKTNAINNF
WP_015404772.1 MRKVSIISMAALGALI....GSTQGFANDHD.QIEGPFETPMEVTATCLECHEDAAKEVMATTHWTWEKMQTVPGKGTVSRGKKNAINNF
1 10 2 20 30 3 40 50 60 70 4
WP_208610160.1 .MSIEANWPRCTSCHVGYGWQSASFDFKDQSKVDCLVCHDTTGEYKKTPTGAGMP..........DPVVDLLYVARNVGKSSRQTCGNCH
WP_029915800.1 CIAVKSNEPRCTSCHIGYGWKDSSFDLTDSSKVDCLVCHDSTGSYKKTPTDAGLP..........AGEVDLLYVARNVGKTSRQTCGTCH
WP_040199676.1 CVSIDSNWPRCTSCHIGYGWEDASFDFTDKTKVDCLVCHDTTGTYKKTATGAGHP..........DEVVDLVYVAKNVGTPSRFNCGSCH
WP_092058436.1 CVSVNANWPRCTSCHIGYGWKDASFDFTKAENIDCLVCHDTTGSYKKTATGAGHP..........DEAVDLMYVARNVGAPKRENCGSCH
WP_072906459.1 CVSINSNWPRCTSCHIGYGWKDASFDFNKPENVDCLVCHDGTGTYKKTPTAAGLP..........AETVDLLYVAKNVAAPKRENCGACH
WP_015404772.1 CVSVGSNEPRCTSCHVGYGWKDGSFDFSDESRVDCLVCHDTTGTYKKPGAGAGMPAGFTGVESMDKKPVDLVKVAQNAGMPTRDNCINCH
80 90 100 110 5 120 130 140 6 150 160
WP_208610160.1 FFGGGGDAVKHGDLDSSIANPTRDIDVHMATDGGNFTCEGCHVTKNHDITGNAMVVSPSNQKHIGCEGCHQGTPH....YESILNKHMKK
WP_029915800.1 FFGGGGDAVKHGDLDSSIDFPTRNLDVHMAADGNNFTCAGCHETKDHNISGQAMVVSPKGQNHIGCTSCHGDKVH....SESILNQHLDT
WP_040199676.1 FFGGGGDAVKHGDLDSSIDYPERETDVHMAVEGNDFTCQTCHATENHEISGNAMVVSPSNQTHIGCESCH.ADPH....AESLLNDHTAA
WP_092058436.1 FFGGGGDAVKHGDLDSSMEYPEKSVDVHMAADGNDFTCQTCHETSGHHITGNAMVVSPGNASHIGCEKCHTEAPH....KESVLNQHGAA
WP_072906459.1 FFGGGGDAVKHGDLDSSMNYPSKEVDVHMDVDGNNFSCQSCHVTEDHQITGNAMAVSPAGQTGIGCEGCHDAAPH....AESILNNHASA
WP_015404772.1 AYGGGGNNVKPGDMSNSMRNPSKDLDVHMDSKGLNFTCQQCHKTDSHDIPGQSLIVSASPGSTLDCTDCHGSSPHRYFPTATKYNAHAEK
170 7 180 190 200 210 220 230 240 250
WP_208610160.1 VACQTCHIPTFAKKVPTKTSWDWSTAGSDMKGGKD.....KYGKPTFAKKKGSFTWGKNIVPTYAWYNGQSHVYQLGDKMDPTKVTKLNE
WP_029915800.1 VACQTCHIPTFAKEVPTKTSWDWSTAGQDIKGEKD.....QYGKPTYMKKKGSFTWGKNIVPTYAWYNGSAGAYQFGDKINPATTTALNY
WP_040199676.1 VACQTCHIPTFAKQIPTKTSWDWSTAGQDLEEAKD.....EHGKPTYMKKKGHFTWDKNIVPEYAWYNGTGGAYIPGDKINPDEMTLLSY
WP_092058436.1 VACQTCHIPVFAKEEATKTSWDWSTAGKDFKEEKD.....EYGRSTYTKAKGHFTWGKKIVPTYAWYNGSAGAYLLGDKMDPSKETLLSY
WP_072906459.1 VACQTCHIPEFAKEHATKTSWDWSTAGQDIKAVKD.....EHGHPNYMKKKGHFTYGQNVQPSYAWYNGTAAAYLMGDKLDPSKTLKITD
WP_015404772.1 IACQTCHIAHFAKADGTKMSWDWSKAATSKEMAKEKKVVKEHGHAVYIFKKGQFTYESNVVPTYAWFNGTTSTYMIGDKIDPSKTTILNA
260 270 280 290 300 310 320 330
WP_208610160.1 PIGSKDDPKSKLYPFKNHKGKQPYDKKNNYFTTVHLFGKGG....YWKTFDWEKSIAAGMKSSGVPFSGEYDFAETIMFWPINHMVSPAE
WP_029915800.1 PIGERSDLKAKIYPFKIHTGKQIYDTKNMYFIPPKVFGKGG....YWKTFDWKQAAEQGAKANGLKFSGEYGFAPTIMYWRINHMVSPAA
WP_040199676.1 PLGDINDPEAKISPFKVHTGKQIYDTEHQYLITPKVFGKDG....YWKTFDWQSAAKLGMEASGLEYSGNYDFASTKMYWSINHMVSPKE
WP_092058436.1 PKGEITDAKAKIFPFKVHTGKQPYDKKHNILITPKVFGDDG....FWKTYDWDQAAKLGMAASNLPFSGDIGFAETKMYWAINHMVSPKE
WP_072906459.1 PNGDINDKSAKIYPFKVMTGKQPYDKKNNILITPKVFGKGG....FWKTFDWDNASKLGMESSGLPYSGEIDFIATEMYWPINHMVSPAE
WP_015404772.1 PQGDKNDPDAKIMPFKIHTGKQIYDAKFNYLITPKVWGPKGDPDAFWVQFDWNKAAAAGMKVSGLEYSGTFDFAPTASYWPINHQVDPKE
340 8 350 360
WP_208610160.1 QALDCLDCHGDKGRLDWKALGYKSDPMKK.......
WP_029915800.1 QALDCLDCHGDNGRLDWGKLGYQGDPLQNPEFARTR
WP_040199676.1 QSLGCLDCHGDNGRLDWKALGYEGDPLRTGKSARLK
WP_092058436.1 EALGCLDCHGDQARMDWKALGYEGDPLRTGKSRAEK
WP_072906459.1 KALGCLDCHGDEGRLDWKQLGYKGDPMQTGKTR...
WP_015404772.1 KALKCLDCHKSGNRMDWKALGYDGDPMAT......K

**AL4**

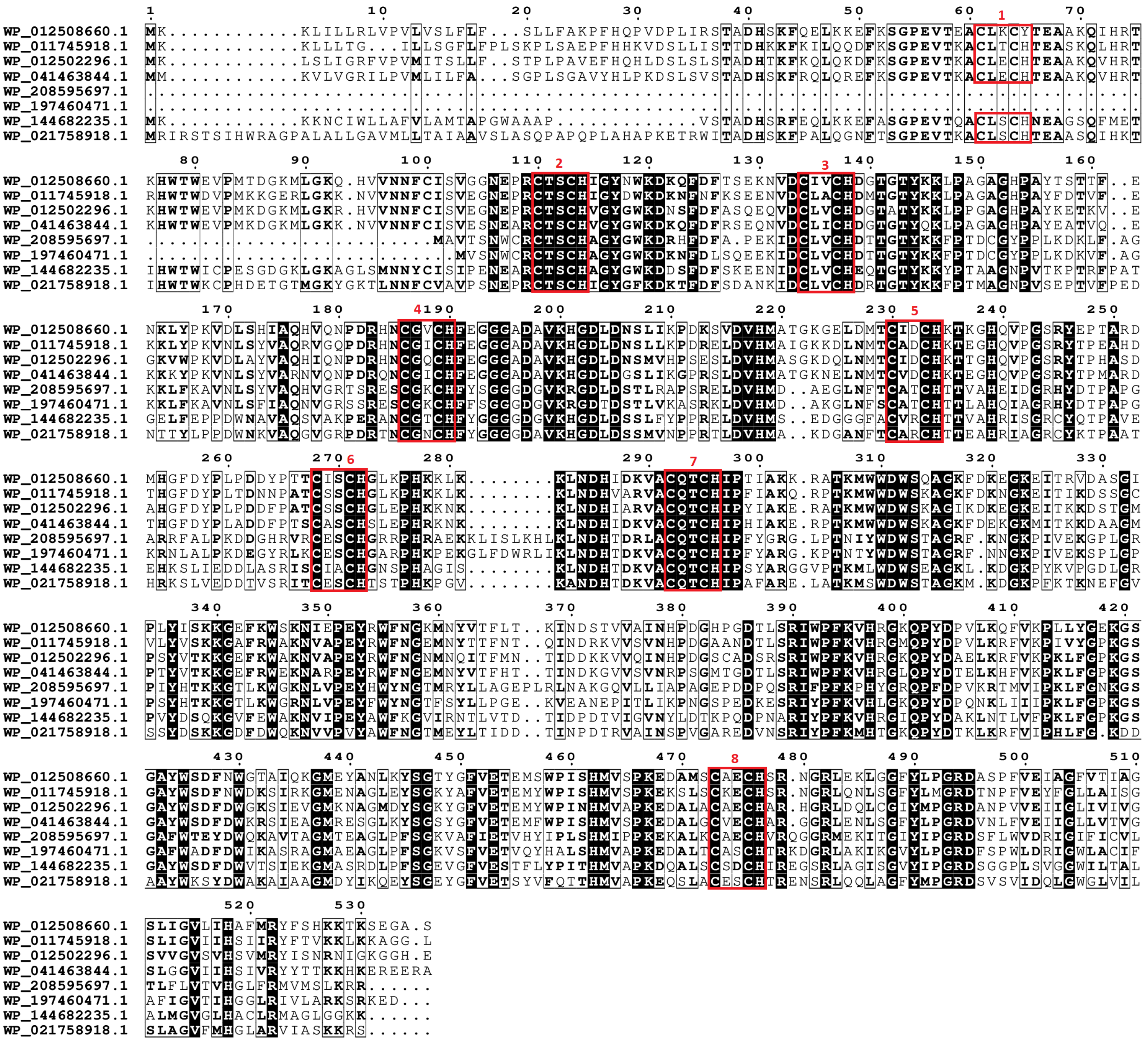
WP_012508660.1
WP_011745918.1
WP_012502296.1
WP_041463844.1
WP_208595697.1
WP_197460471.1
WP_144682235.1
WP_021758918.1

AL5

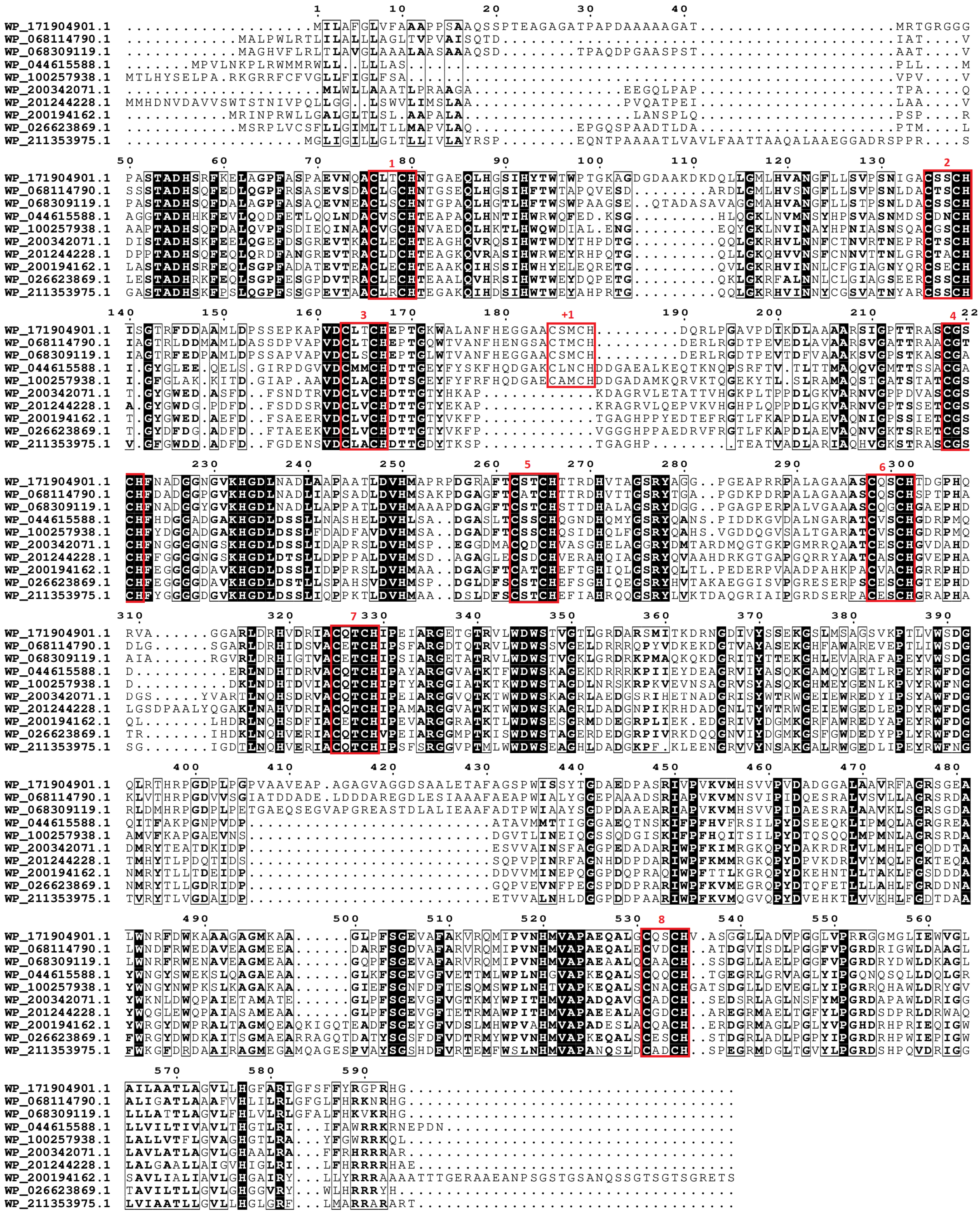

Figure S12. Maximum likelihood phylogenetic tree of OTR family. Each presented tip is labeled with the RefSeq accession code, number of heme-binding motifs and if it contains a paralogue within this analysis and taxonomic class. Confidence values (expressed in %) of SH-aLRT/ultrafast bootstrap are presented near each node. Each heme-binding motif gain (green) / loss (blue) event was considered when confidence values are ≥ 80%/95%, respectively, otherwise only highlighted in gray. Confidence values below 70% are not shown. At the bottom of the tree, a subset of the aligned sequences is presented (those that are in bold in the phylogenetic tree) that are related to each event of heme-binding motif gain / loss (AL1 to AL5).

#### MtrA

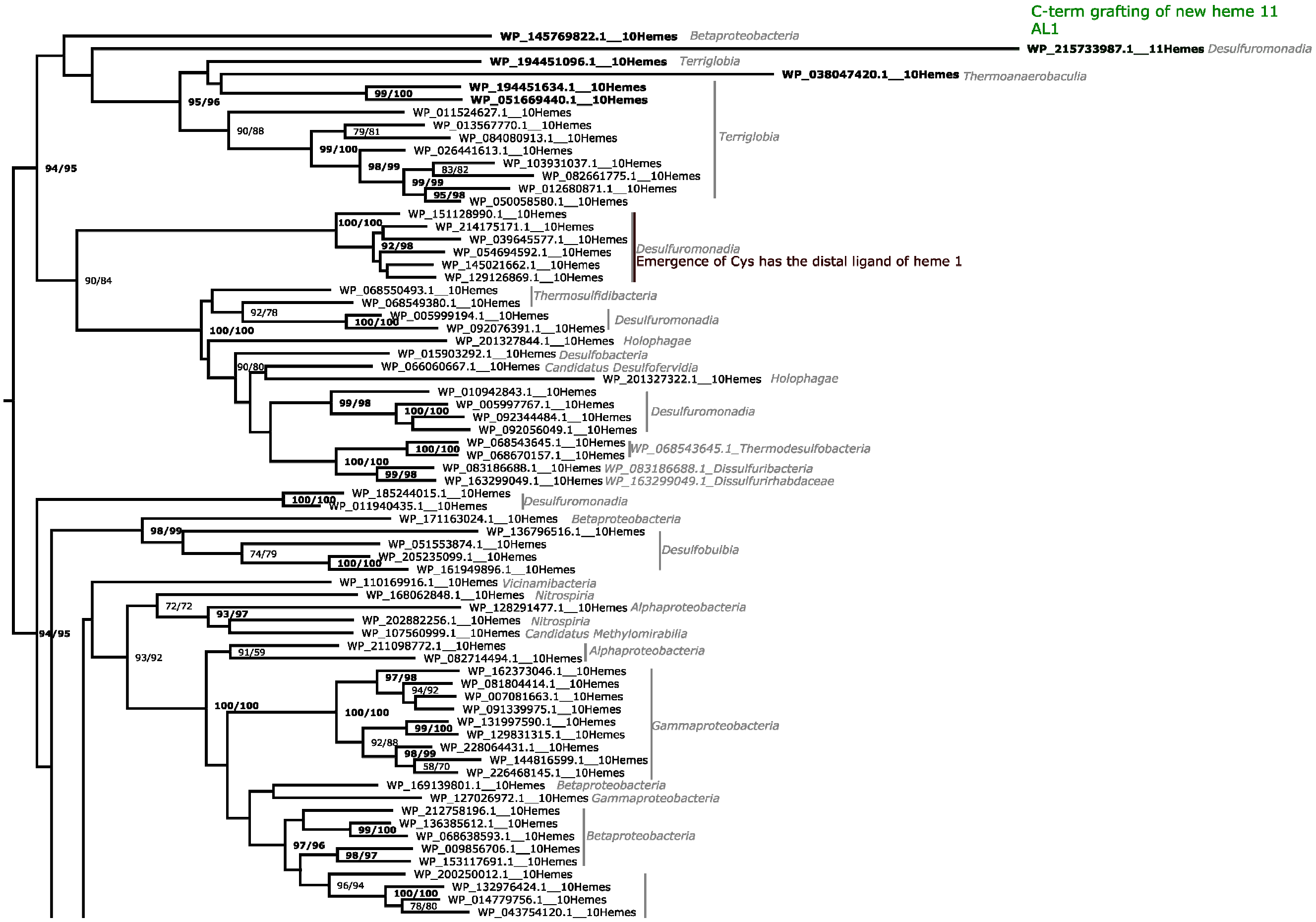


(Continues in next page)

(Continued)

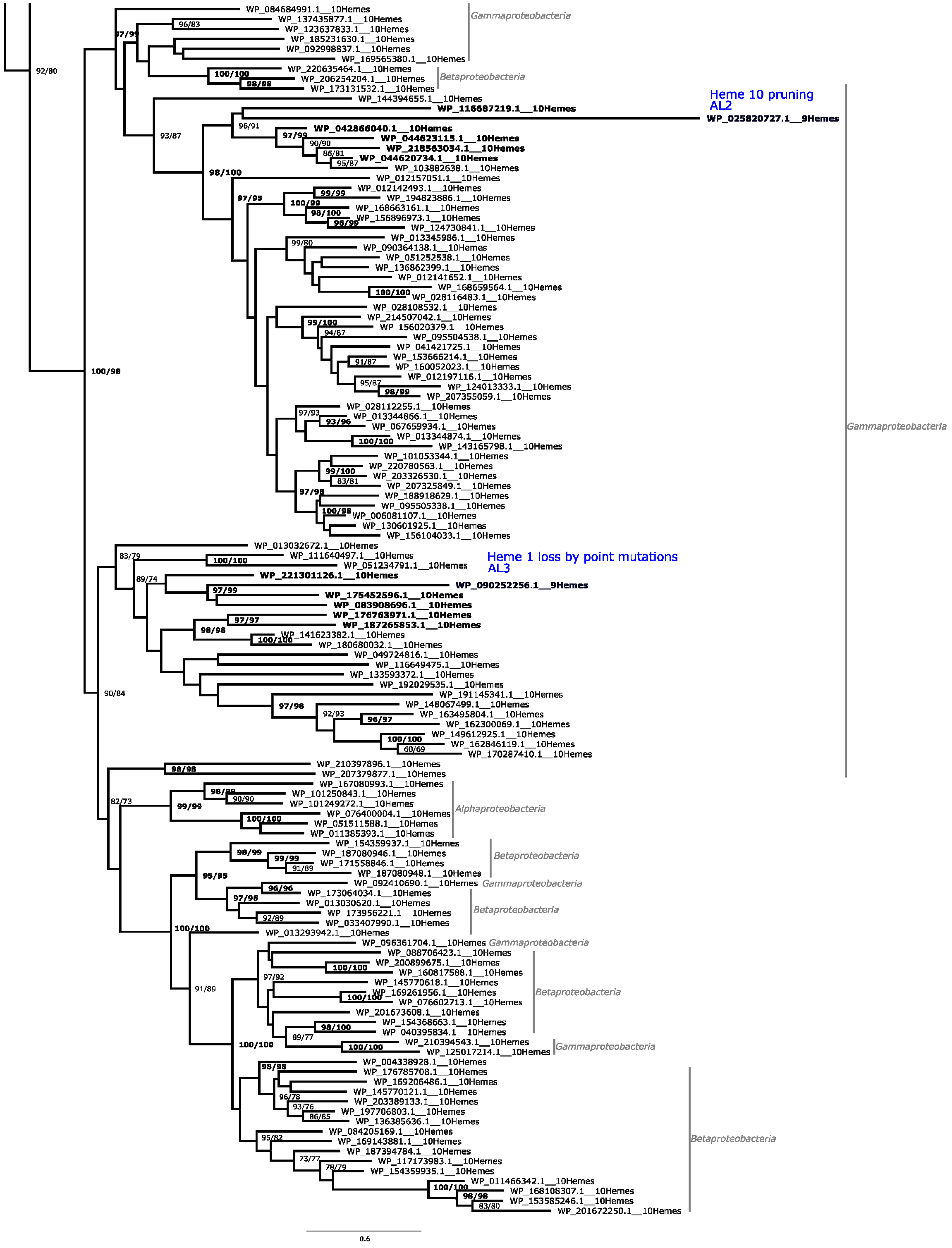

Heme 10 pruning
AL2
Heme 1 loss by point mutations
AL3
Gammaproteobacteria
Betaproteobacteria
Alphaproteobacteria
WP_116687219.1__10Hemes
WP_025820727.1__9Hemes
WP_090252256.1__9Hemes
0.6

AL1

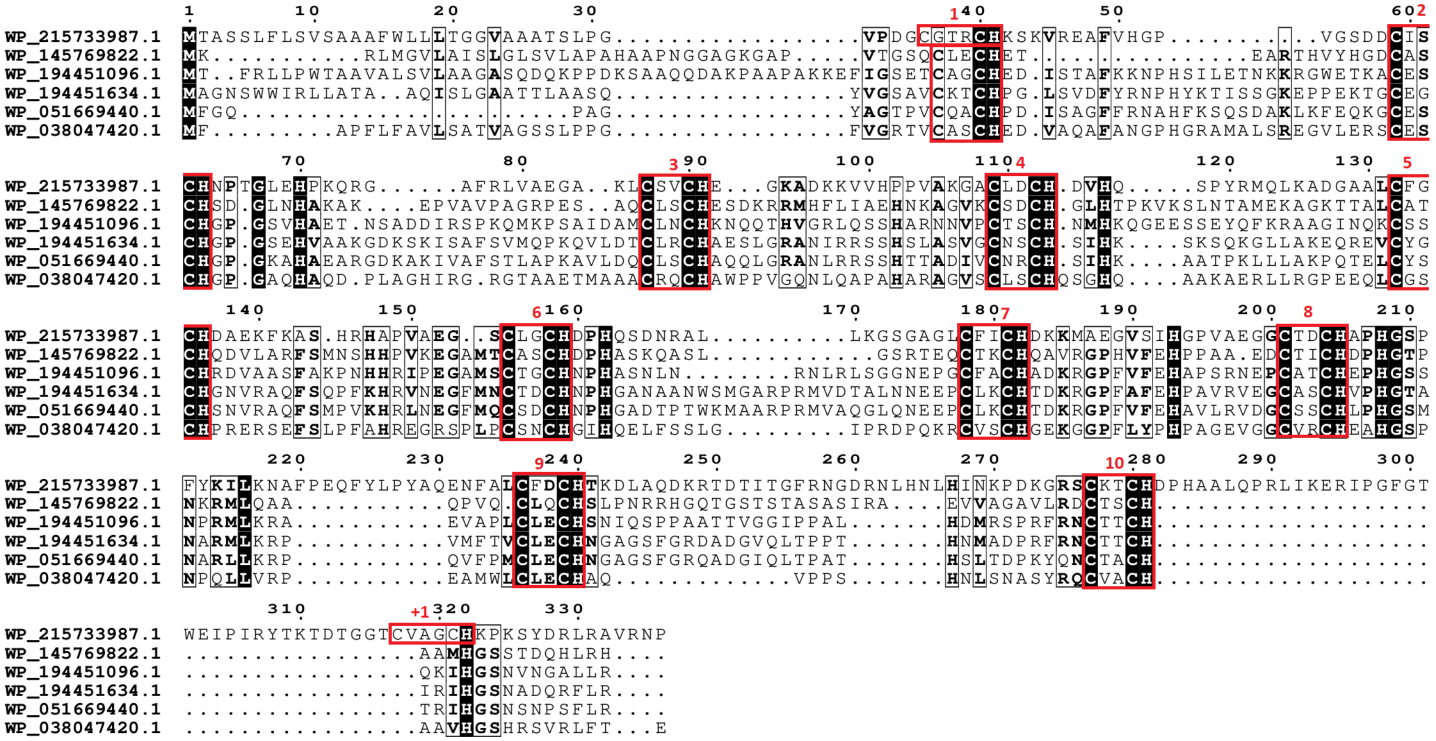
WP_215733987.1
WP_145769822.1
WP_194451096.1
WP_194451634.1
WP_051669440.1
WP_038047420.1

AL2

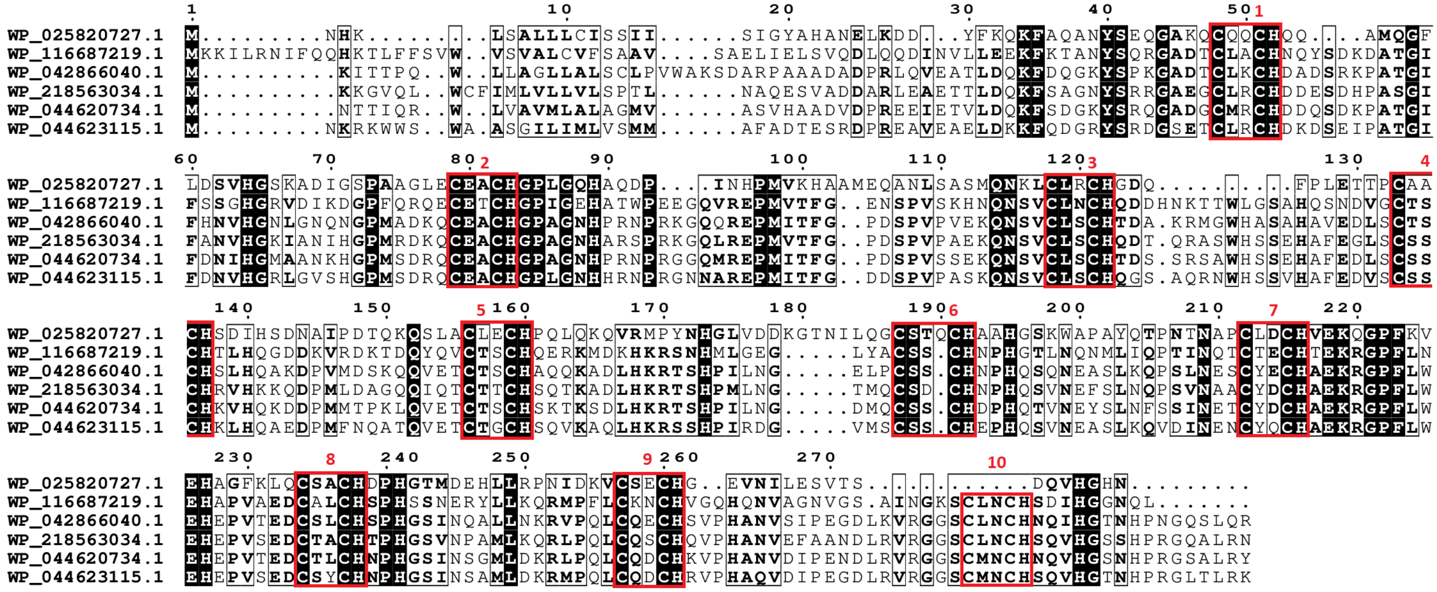
WP_025820727.1
WP_116687219.1
WP_042866040.1
WP_218563034.1
WP_044620734.1
WP_044623115.1

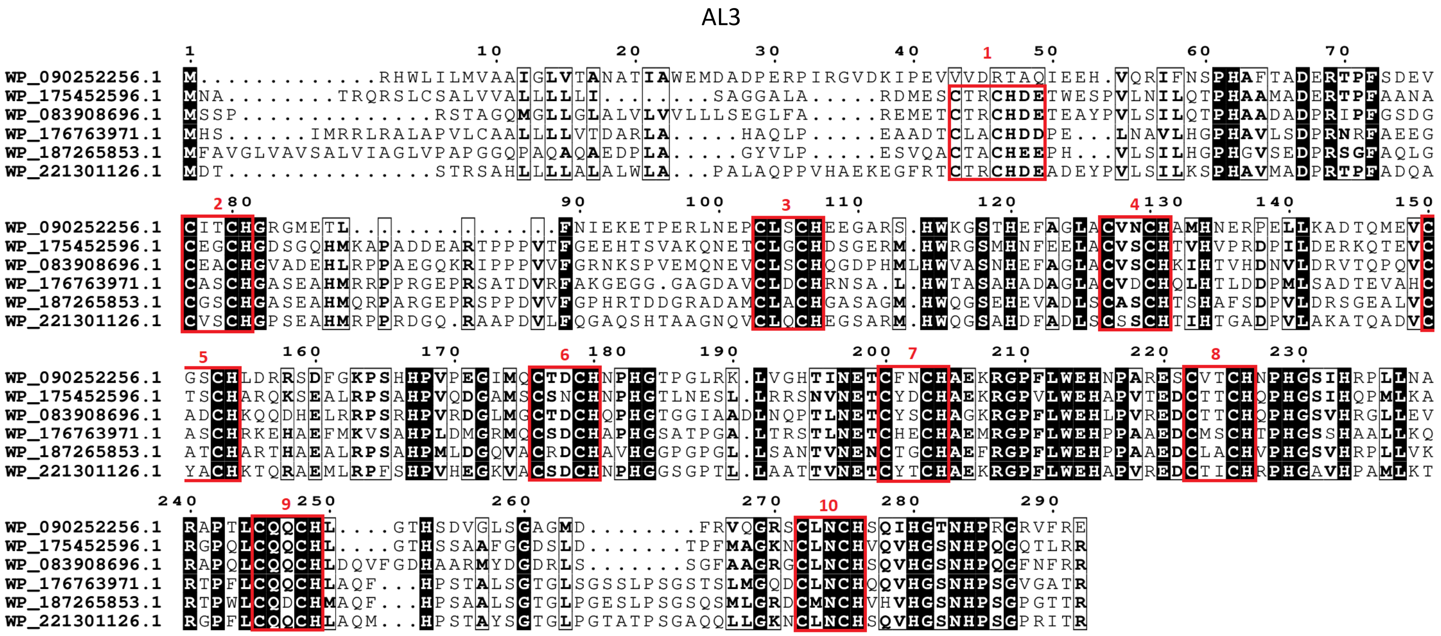


Figure S13. Maximum likelihood phylogenetic tree of MtrA family. Each presented tip is labeled with the RefSeq accession code, number of heme-binding motifs and if it contains a paralogue within this analysis and taxonomic class. Confidence values (expressed in %) of SH-aLRT/ultrafast bootstrap are presented near each node. Each heme-binding motif gain (green) / loss (blue) event was considered when confidence values are ≥ 80%/95%, respectively, otherwise only highlighted in gray. Confidence values below 70% are not shown. At the bottom of the tree, a subset of the aligned sequences is presented (those that are in bold in the phylogenetic tree) that are related to each event of heme-binding motif gain / loss (AL1 to AL3).

**UndA**

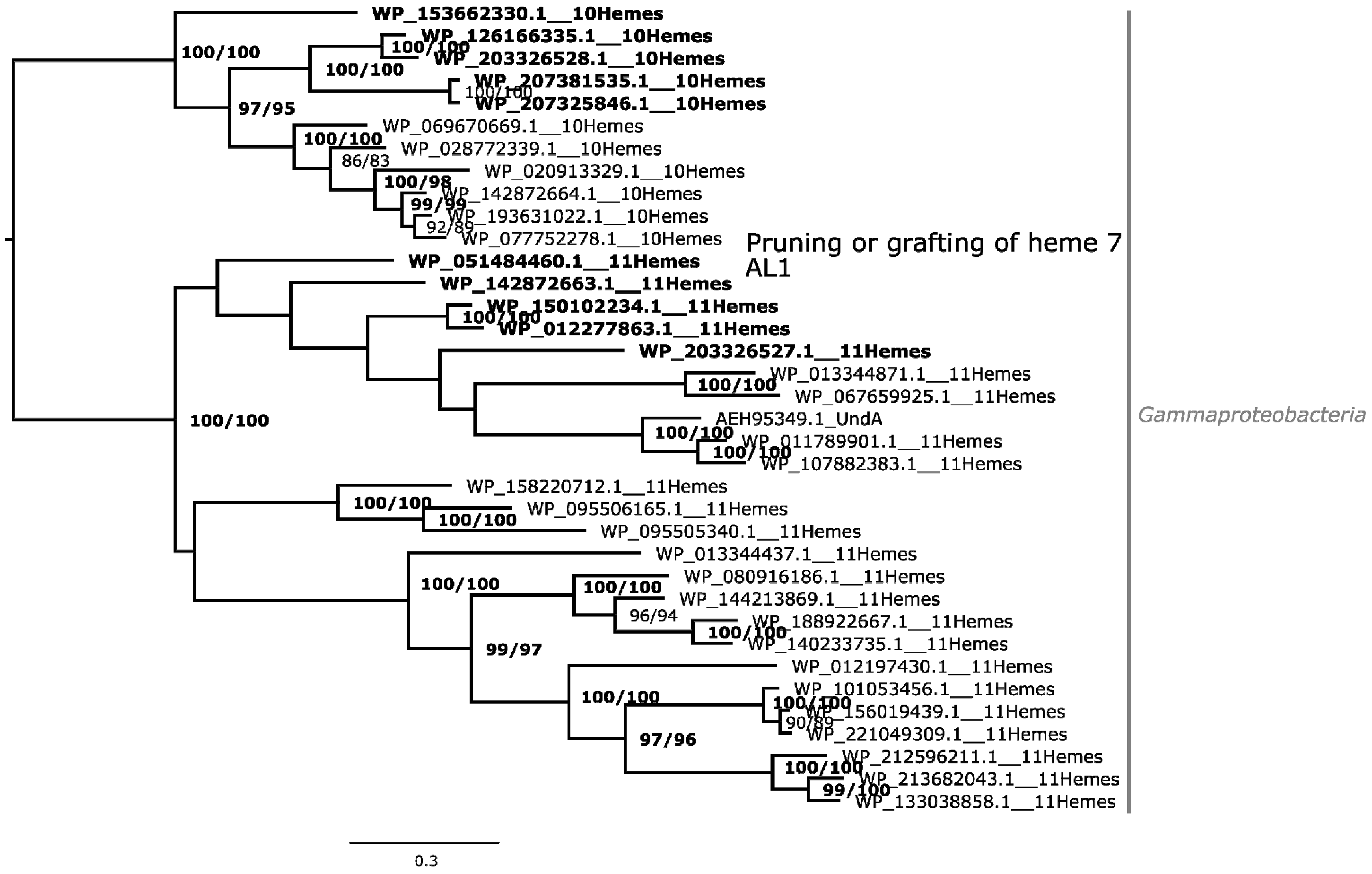
WP_153662330.1__10Hemes
WP_126166335.1__10Hemes
WP_203326528.1__10Hemes
WP_207381535.1__10Hemes
WP_207325846.1__10Hemes
WP_069670669.1__10Hemes
WP_028772339.1__10Hemes
WP_020913329.1__10Hemes
WP_142872664.1__10Hemes
WP_193631022.1__10Hemes
WP_077752278.1__10Hemes
Pruning or grafting of heme 7
AL1
WP_051484460.1__11Hemes
WP_142872663.1__11Hemes
WP_150102234.1__11Hemes
WP_012277863.1__11Hemes
WP_203326527.1__11Hemes
WP_013344871.1__11Hemes
WP_067659925.1__11Hemes
AEH95349.1_UndA
WP_011789901.1__11Hemes
WP_107882383.1__11Hemes
WP_158220712.1__11Hemes
WP_095506165.1__11Hemes
WP_095505340.1__11Hemes
WP_013344437.1__11Hemes
WP_080916186.1__11Hemes
WP_144213869.1__11Hemes
WP_188922667.1__11Hemes
WP_140233735.1__11Hemes
WP_012197430.1__11Hemes
WP_101053456.1__11Hemes
WP_156019439.1__11Hemes
WP_221049309.1__11Hemes
WP_212596211.1__11Hemes
WP_213682043.1__11Hemes
WP_133038858.1__11Hemes
Gammaproteobacteria
0.3

AL1

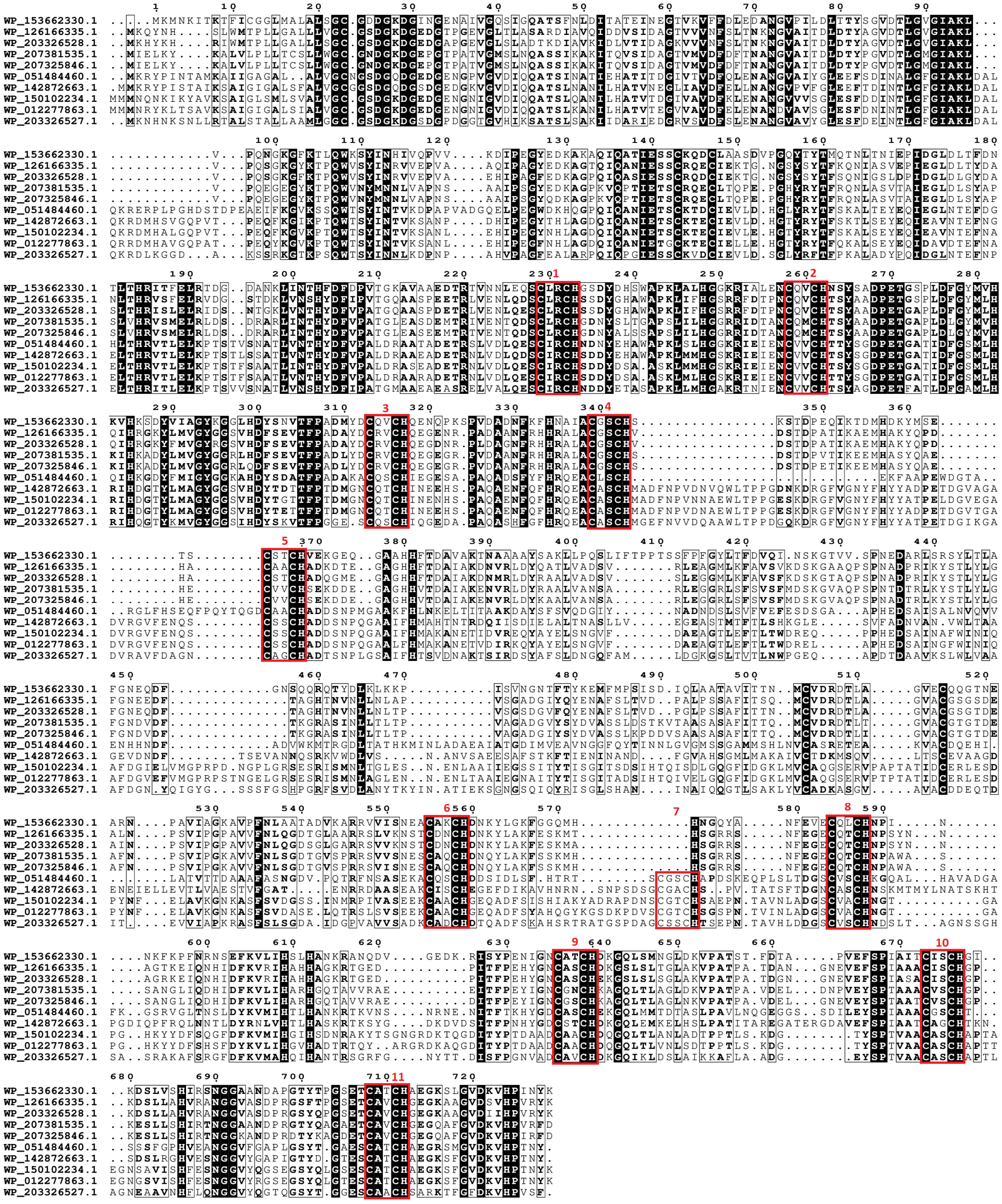


Figure S14. Maximum likelihood phylogenetic tree of UndA family. Each presented tip is labeled with the RefSeq accession code, number of heme-binding motifs and if it contains a paralogue within this analysis and taxonomic class. Confidence values (expressed in %) of SH-aLRT/ultrafast bootstrap are presented near each node. Each heme-binding motif gain (green) / loss (blue) event was considered when confidence values are ≥ 80%/95%, respectively, otherwise only highlighted in gray. Confidence values below 70% are not shown. At the bottom of the tree, a subset of the aligned sequences is presented (those that are in bold in the phylogenetic tree) that are related to each event of heme-binding motif gain / loss (AL1).

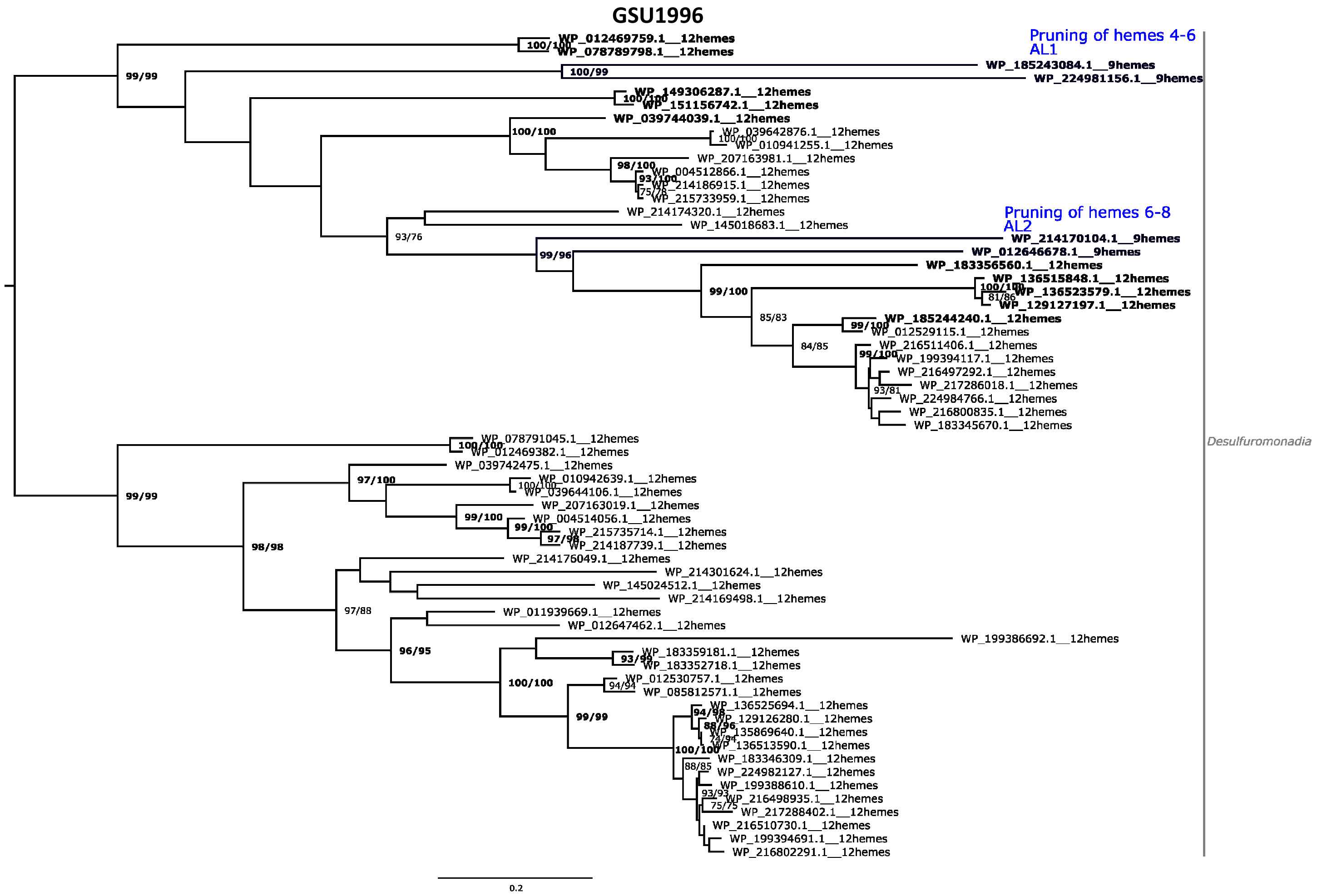
GSU1996
Pruning of hemes 4-6
AL1
Pruning of hemes 6-8
AL2
Desulfuromonadia
0.2

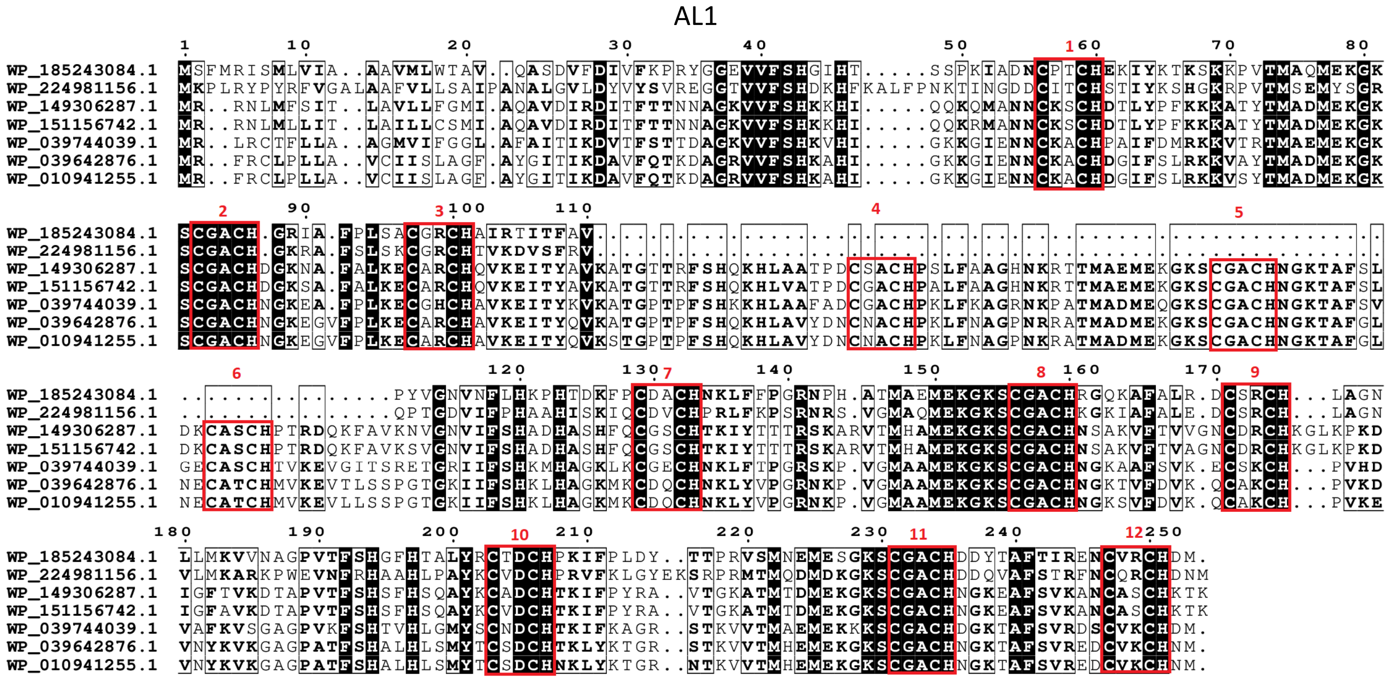
AL1
WP_185243084.1
WP_224981156.1
WP_149306287.1
WP_151156742.1
WP_039744039.1
WP_039642876.1
WP_010941255.1

AL2

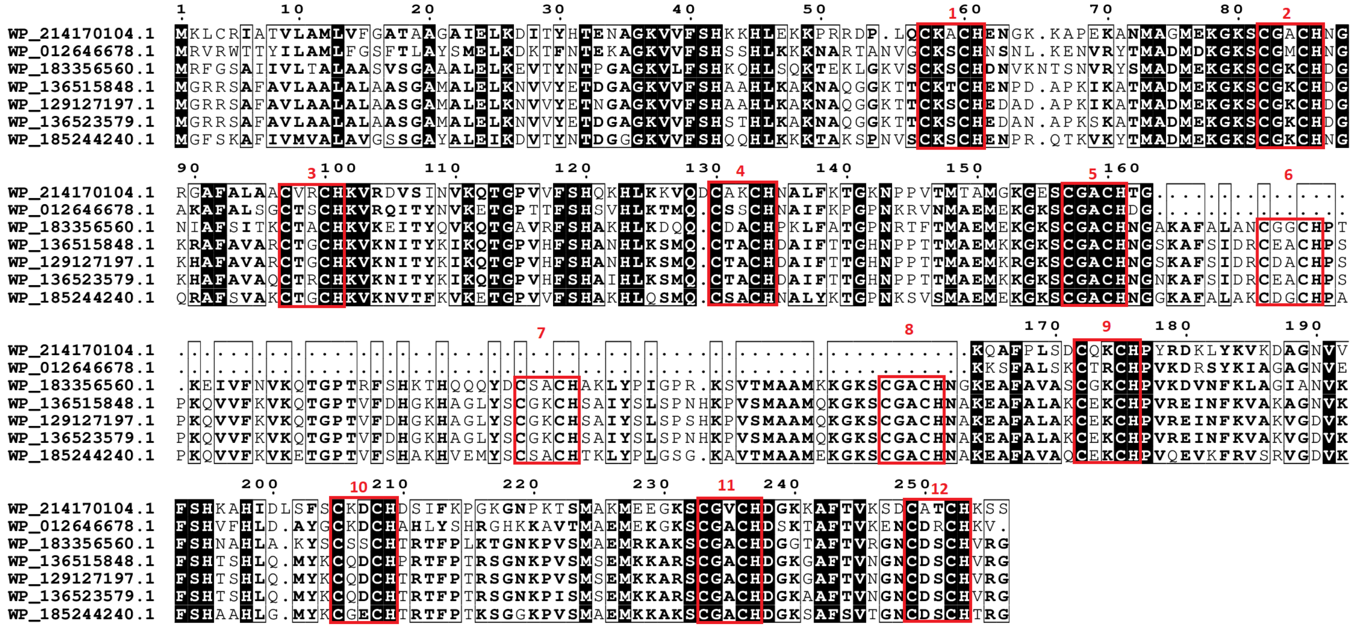


Figure S15. Maximum likelihood phylogenetic tree of GSU1996 family. Each presented tip is labeled with the RefSeq accession code, number of heme-binding motifs and if it contains a paralogue within this analysis and taxonomic class. Confidence values (expressed in %) of SH-aLRT/ultrafast bootstrap are presented near each node. Each heme-binding motif gain (green) / loss (blue) event was considered when confidence values are ≥ 80%/95%, respectively, otherwise only highlighted in gray. Confidence values below 70% are not shown. At the bottom of the tree, a subset of the aligned sequences is presented (those that are in bold in the phylogenetic tree) that are related to each event of heme-binding motif gain / loss (AL1 and AL2).

#### 16HmcA

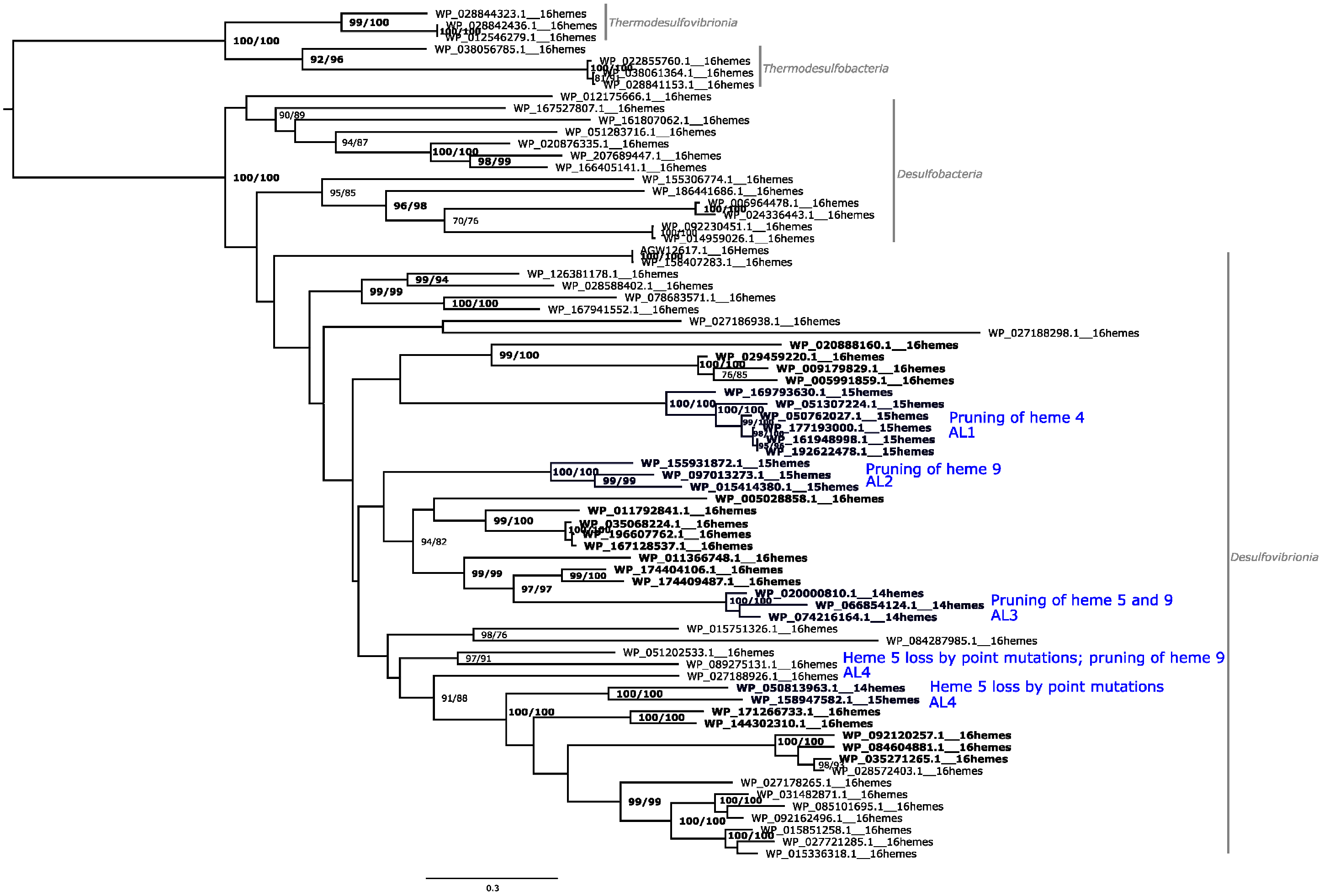

WP_028844323.1__16hemes
WP_028842436.1__16hemes
WP_012546279.1__16hemes
WP_038056785.1__16hemes
Thermodesulfovibrionia
WP_022855760.1__16hemes
WP_038061364.1__16hemes
WP_028841153.1__16hemes
Thermodesulfobacteria
WP_012175666.1__16hemes
WP_167527807.1__16hemes
WP_161807062.1__16hemes
WP_051283716.1__16hemes
WP_020876335.1__16hemes
WP_207689447.1__16hemes
WP_166405141.1__16hemes
WP_155306774.1__16hemes
WP_186441686.1__16hemes
WP_006964478.1__16hemes
WP_024336443.1__16hemes
WP_092230451.1__16hemes
WP_014959026.1__16hemes
Desulfobacteria
AGW12617.1__16Hemes
WP_158407283.1__16hemes
WP_126381178.1__16hemes
WP_028588402.1__16hemes
WP_078683571.1__16hemes
WP_167941552.1__16hemes
WP_027186938.1__16hemes
WP_027188298.1__16hemes
WP_020888160.1__16hemes
WP_029459220.1__16hemes
WP_009179829.1__16hemes
WP_005991859.1__16hemes
WP_169793630.1__15hemes
WP_051307224.1__15hemes
WP_050762027.1__15hemes
WP_177193000.1__15hemes
WP_161948998.1__15hemes
WP_192622478.1__15hemes
Pruning of heme 4
AL1
WP_155931872.1__15hemes
WP_097013273.1__15hemes
WP_015414380.1__15hemes
Pruning of heme 9
AL2
WP_005028858.1__16hemes
WP_011792841.1__16hemes
WP_035068224.1__16hemes
WP_196607762.1__16hemes
WP_167128537.1__16hemes
WP_011366748.1__16hemes
WP_174404106.1__16hemes
WP_174409487.1__16hemes
WP_020000810.1__14hemes
WP_066854124.1__14hemes
WP_074216164.1__14hemes
Pruning of heme 5 and 9
AL3
WP_015751326.1__16hemes
WP_084287985.1__16hemes
WP_051202533.1__16hemes
WP_089275131.1__16hemes
WP_027188926.1__16hemes
Heme 5 loss by point mutations; pruning of heme 9
AL4
WP_050813963.1__14hemes
WP_158947582.1__15hemes
Heme 5 loss by point mutations
AL4
WP_171266733.1__16hemes
WP_144302310.1__16hemes
WP_092120257.1__16hemes
WP_084604881.1__16hemes
WP_035271265.1__16hemes
WP_028572403.1__16hemes
WP_027178265.1__16hemes
WP_031482871.1__16hemes
WP_085101695.1__16hemes
WP_092162496.1__16hemes
WP_015851258.1__16hemes
WP_027721285.1__16hemes
WP_015336318.1__16hemes
Desulfovibrionia
0.3

AL1

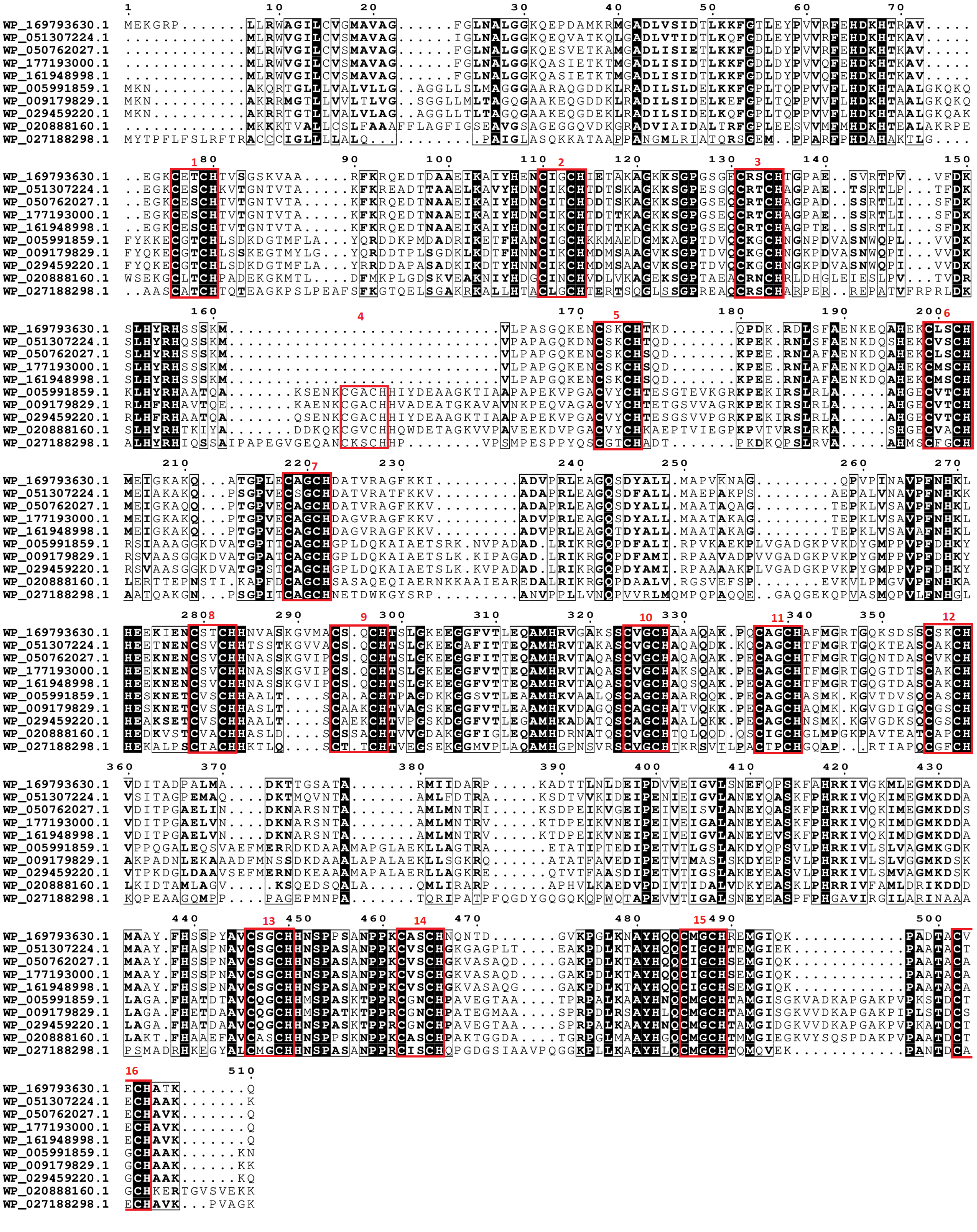

AL2

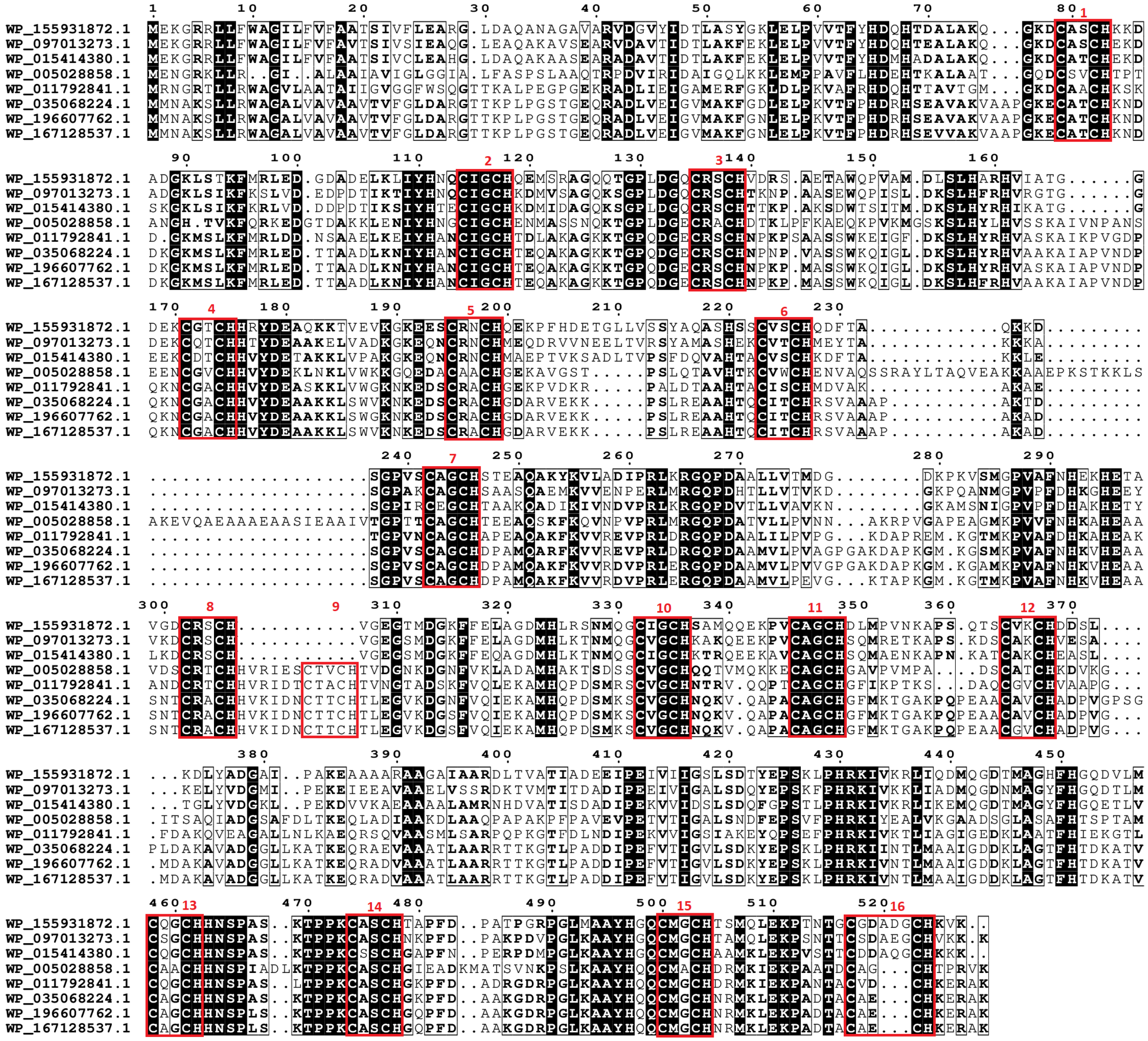
WP_155931872.1
WP_097013273.1
WP_015414380.1
WP_005028858.1
WP_011792841.1
WP_035068224.1
WP_196607762.1
WP_167128537.1

AL3

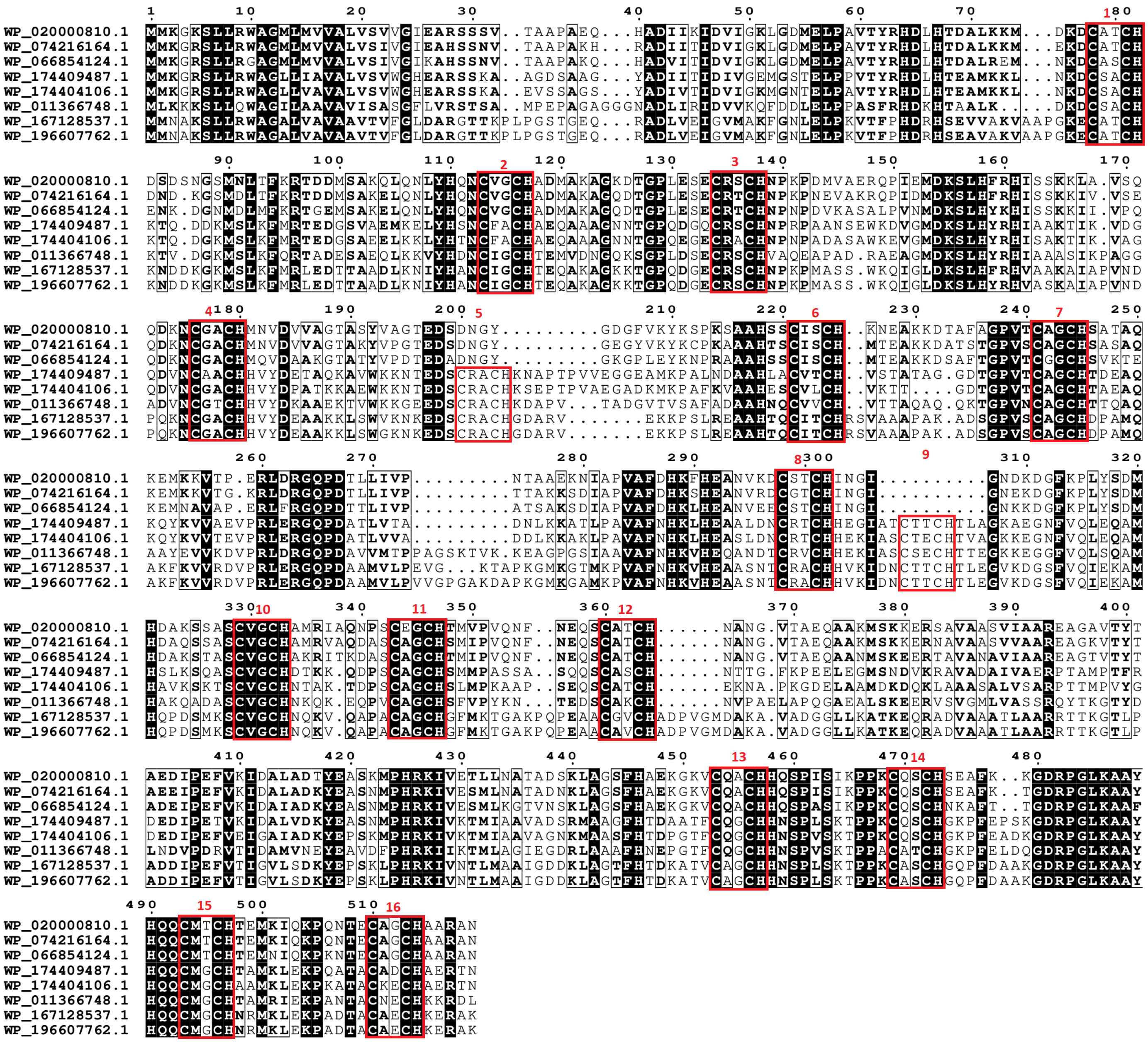
WP_020000810.1
WP_074216164.1
WP_066854124.1
WP_174409487.1
WP_174404106.1
WP_011366748.1
WP_167128537.1
WP_196607762.1

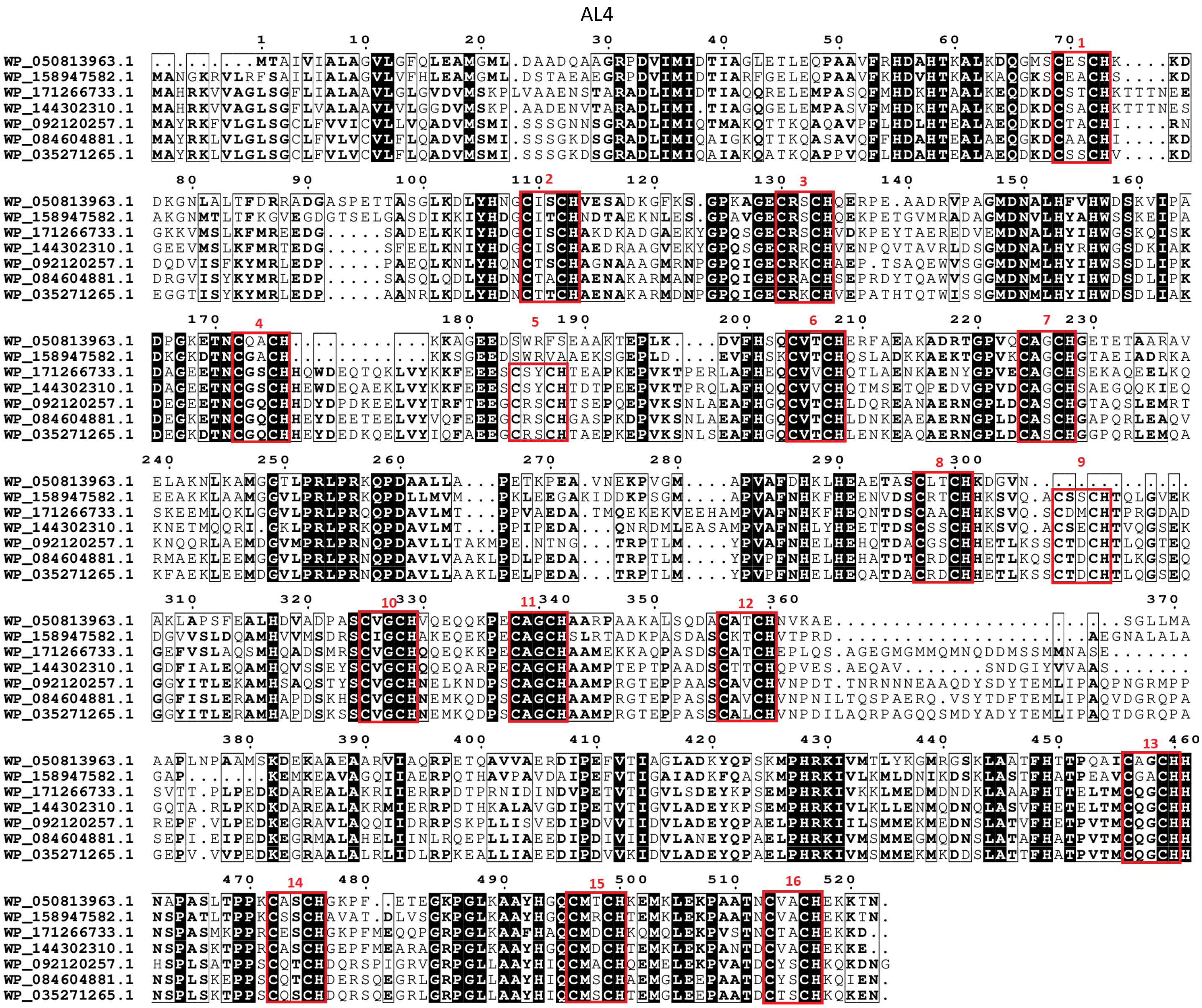


Figure S16. Maximum likelihood phylogenetic tree of UndA family. Each presented tip is labeled with the RefSeq accession code, number of heme-binding motifs and if it contains a paralogue within this analysis and taxonomic class. Confidence values (expressed in %) of SH-aLRT/ultrafast bootstrap are presented near each node. Each heme-binding motif gain (green) / loss (blue) event was considered when confidence values are ≥ 80%/95%, respectively, otherwise only highlighted in gray. Confidence values below 70% are not shown. At the bottom of the tree, a subset of the aligned sequences is presented (those that are in bold in the phylogenetic tree) that are related to each event of heme-binding motif gain / loss (AL1 to AL4).

a

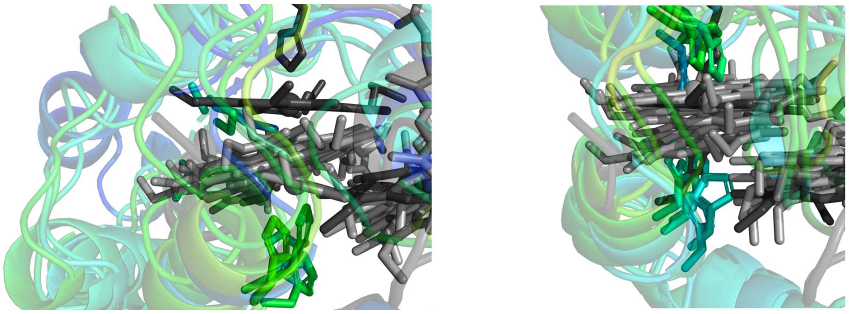

$Cx_3CH$ + Cys ligand $Cx_2CH$ + His ligand

b

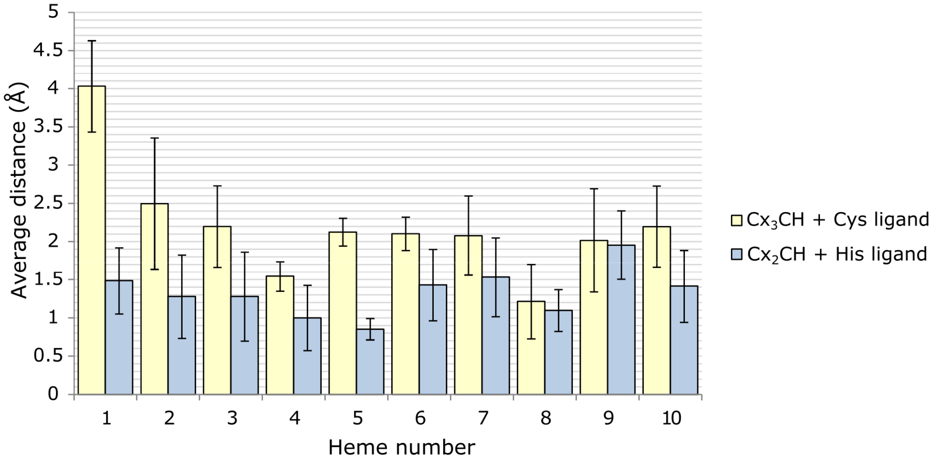


Figure S17. Structure prediction using RoseTTAFold2 for the hemecore alignment of the MtrA sequences containing the alternative heme 1 ($Cx_3CH$ + Cys ligand) and closely related sequences ($Cx_2CH$ + His ligand) against the reference MtrA structure (PDB: 6R2Q_A). **a**. Heme 1 arrangement for the MtrA sequences containing the alternative heme 1 ($Cx_3CH$ + Cys ligand) and close related sequences ($Cx_2CH$ + His ligand) when all hemes are aligned to the MtrA reference structure (in black). Polypeptide chain is colored from red to blue according to the pLDTT confidence values. **b.** Average distance between the heme rings (iron and nitrogen atoms) of the predicted structures ($Cx_3CH$ + Cys ligand and $Cx_2CH$ + His ligand) and the reference MtrA structure (PDB: 6R2Q_A). Error bars represent confidence interval at 95%.

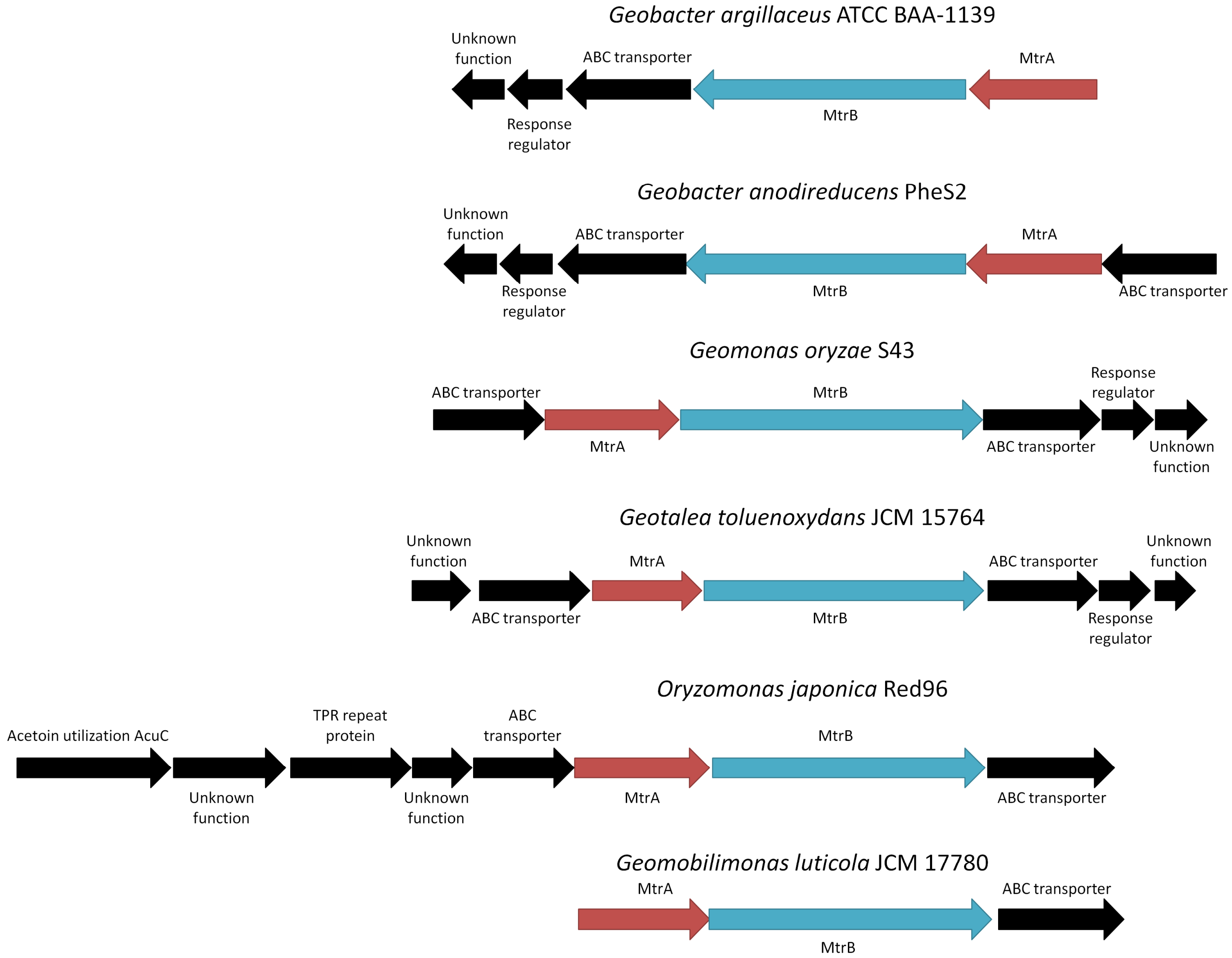


Figure S18. Operon organization for the MtrA homologue sequences of the $Cx_3CH$ + Cys ligand. Sizes of the arrows are relative with their gene length. Red arrows correspond to MHC (MtrA homologues), blue to MtrB homologues and black to other proteins that are part of the same operon.

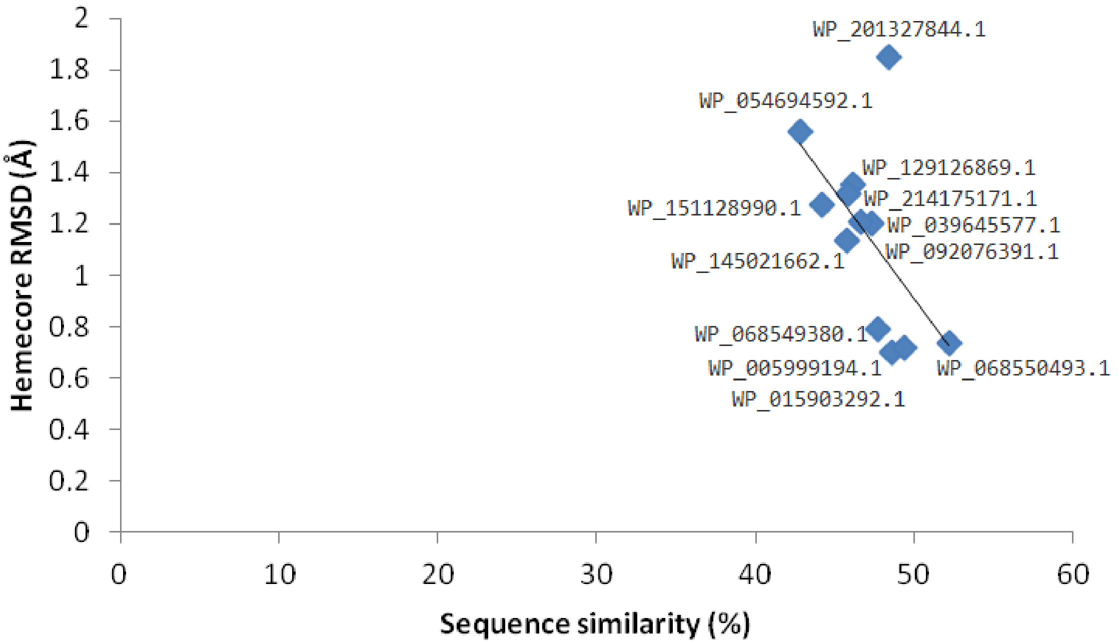


Fig. S19. Linear regression of the Hemecore RMSD of aligned predicted structures of the $Cx_2CH$ + His ligand and $Cx_3CH$ + Cys ligand MtrA homologs against MtrA reference structure (PDB: 6R2Q_A) and the sequence similarity. The predicted structure from the sequence with accession code WP_201327844.1 was removed from further structural comparison as it is plotted far from the trend of hemecore distance/sequence similarity.
